# Suppressor of quenching 1 functions as a methionine sulfoxide reductase in the chloroplast lumen for regulation of photoprotective qH in Arabidopsis

**DOI:** 10.1101/2024.11.01.621559

**Authors:** Jingfang Hao, Alexander Johansson, Johan Svensson Fall, Jianli Duan, Alexander P. Hertle, Matthew D. Brooks, Krishna K. Niyogi, Keisuke Yoshida, Toru Hisabori, Alizée Malnoë

## Abstract

Photosynthetic organisms must balance light absorption and energy dissipation to prevent photo-oxidative damage. Non-photochemical quenching (NPQ) dissipates excess light energy as heat, with the quenching component qH providing sustained photoprotection. However, the molecular mechanism underlying qH induction remains unclear. Our study focuses on the thylakoid membrane protein SUPPRESSOR OF QUENCHING 1 (SOQ1) and its inhibition of qH through interaction with LIPOCALIN IN THE PLASTID (LCNP) in *Arabidopsis thaliana*. Structural homology of SOQ1 lumenal domains with bacterial disulfide bond protein D suggested potential thiol-disulfide exchange activity. *In vitro* assays determined that both SOQ1 thioredoxin-like (Trx-like) and C-terminal (CTD) domains contain a redox-active cysteine pair and evidenced electron transfer from Trx-like to CTD. Importantly, we found that SOQ1 lumenal domains exhibit methionine sulfoxide reductase (Msr) activity converting oxidized methionine residues in LCNP back to methionine, which thereby inactivates LCNP and prevents qH formation. Mutational analyses identified cysteine residues in SOQ1-CTD and methionine residues in LCNP as critical for qH suppression, supporting their role in redox regulation. Additionally, we found that the redox state of SOQ1 *in vivo* is light-dependent, shifting from reduced to oxidized under stress conditions, indicating a dynamic regulation of its activity. We conclude that the Trx-like domain of SOQ1 provides reducing power to its CTD displaying Msr activity. SOQ1 is therefore an unusual example of a protein possessing both a disulfide reductase and Msr domain in tandem. Our findings elucidate the redox-regulation mechanism of qH involving SOQ1-mediated methionine reduction of LCNP, providing insights into the intricate control of photoprotective processes in chloroplasts and enhancing our understanding of plant resilience under environmental stress.

## Introduction

Photosynthesis is an essential biological process that converts light energy into chemical energy, supporting plant growth and food production. However, environmental stressors can significantly impact its efficiency. Excessive light intensity beyond photosynthetic capacity can lead to the formation of reactive oxygen species (ROS) in chloroplasts, causing photo-oxidative damage (Aro et al., 1993; Niyogi, 1999). To prevent this damage, plants have evolved photoprotective mechanisms. One such mechanism dissipates excess absorbed light energy as heat through a process called non-photochemical quenching (NPQ) (Horton et al., 1996). Energy dissipation is critical for survival and resilience of photosynthetic organisms while contributing to their overall environmental adaptability and productivity (De Souza et al., 2022).

NPQ appears as a decrease of chlorophyll fluorescence and is comprised of several components that differ in kinetics and molecular players involved (Malnoë, 2018; Bassi and Dall’Osto, 2021). Among these components, qE is the fastest, initiating within seconds after exposure to light and relaxing within one-two minutes after return to darkness. It relies on a pH gradient across the thylakoid membrane (ΔpH), the PSII subunit S (PsbS), and the conversion of violaxanthin to zeaxanthin catalyzed by violaxanthin de-epoxidase (VDE) (Li et al., 2000; Ruban et al., 2012; Jahns and Holzwarth, 2012). The protonation of PsbS and VDE allows for qE formation by altering the PsbS lumenal loop conformation (Fan et al., 2015; Krishnan-Schmieden et al., 2021) and promoting VDE dimerization (Arnoux et al., 2009); the photosystem II (PSII) light-harvesting antenna also undergoes small conformational changes in low pH (Ruan et al., 2023). Alternatively, light harvesting can be regulated through phosphorylation of stromal residues in the antenna by the STN7 kinase (Bellafiore et al., 2005). This NPQ component, known as qT (state transitions), redistributes the antenna from PSII to photosystem I (PSI), and its relaxation takes 5-10 minutes (Quick and Stitt, 1989). The redox status of quinone in the thylakoid membrane is read by STN7; the reduction of quinone to quinol is the signal for its activation (Shapiguzov et al., 2016). Recently, a slower NPQ component, qH, was identified in *Arabidopsis thaliana* (Brooks et al., 2013; Malnoë et al., 2018), which takes hours to deactivate (thus referred to as “sustained”). The molecular signal for qH induction is unknown and serves as the motivation for this study.

qH limits lipid peroxidation, providing photoprotection (Malnoë et al., 2018). It occurs in both major and minor antenna complexes (Bru et al., 2022, Bru et al. in preparation) and requires a lumenal protein, LIPOCALIN IN THE PLASTID (LCNP), for initiation (Malnoë et al., 2018), and a stromal protein, RELAXATION OF QH1 (ROQH1), for termination (Amstutz et al., 2020). The thylakoid membrane-anchored protein SUPPRESSOR OF QUENCHING 1 (SOQ1) suppresses qH formation by inhibiting LCNP through its lumenal domains (Brooks et al., 2013; Malnoë et al., 2018; Yu et al., 2022). SOQ1 contains a haloacid dehalogenase-like hydrolase (HAD) domain, a transmembrane helix (TM), and lumenal domains including a thioredoxin-like domain with an atypical CCINC motif (Trx), a β-propeller NHL repeat (NHL), and a C-terminal domain (CTD) (Brooks et al., 2013; Yu et al., 2022) (Fig. 1A). SOQ1 domains can accumulate as truncated forms: TM-Trx-NHL-CTD (TM-TNC) in the thylakoid membrane and Trx-NHL-CTD (TNC), NHL-CTD (NC) and CTD in the thylakoid lumen (Yu et al., 2022). Whether these truncated forms of SOQ1 are functional is investigated here. Mutations in conserved cysteine residues within the SOQ1 Trx-like motif abolish its ability to suppress NPQ, indicating a redox function (Brooks et al., 2013).

**Fig. 1.**
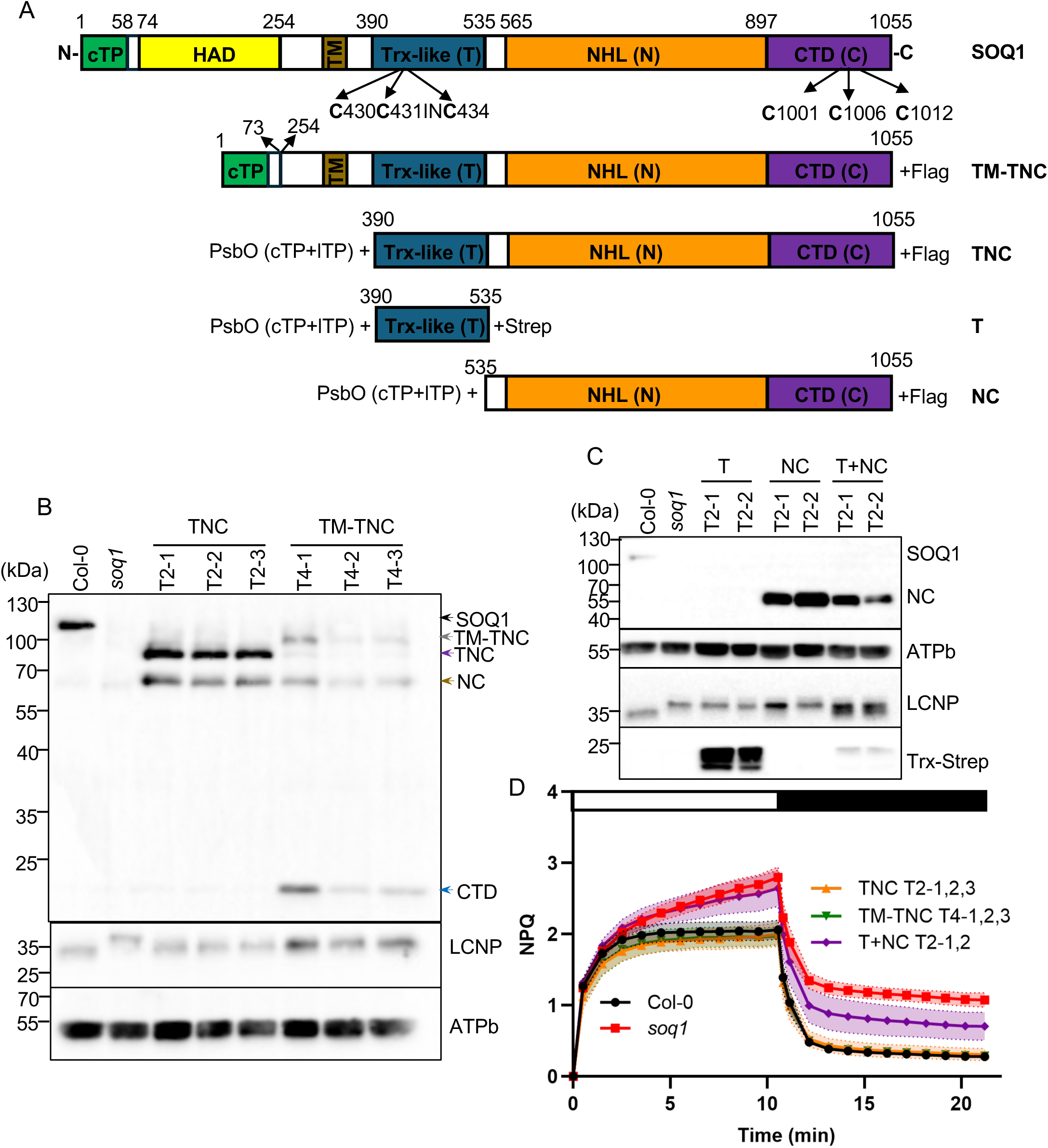
SOQ1 lumenal domains rescue a wild-type NPQ phenotype in the *soq1* mutant. A, Schematic of SOQ1 constructs transformed into the *soq1* mutant. SOQ1 contains a chloroplast transit peptide (cTP; green), HAD (yellow), transmembrane (TM; dark yellow), Trx-like (T, blue), NHL (N, orange) and CTD (C, purple) domains. The chloroplast (cTP) and lumenal transit peptide (lTP) sequences of PsbO were fused to the N-terminal of TNC, T or NC. Numbers indicate amino acid positions. The arrows show the cysteines in the CCINC motif of Trx-like domain and CTD domain. B, C, Immunoblot analyses of two or three independent T2 or T4 *Arabidopsis* variant lines containing SOQ1 lumenal domains Trx-NHL-CTD (TNC) or the transmembrane domain (TM) with the lumenal domains TNC (TM-TNC) or Trx (T) or NHL-CTD (NC) or T+NC expressed as separate entities. The constructs were transformed in the *soq1* knock-out mutant background. Wild type (Col-0) is used as a reference. Samples were loaded at the same amount of total protein (15 µg). Proteins were separated by SDS-PAGE and analyzed by immunodetection with antibodies against SOQ1_CTD_, LCNP, Strep-tag or ATPb. ATPb is shown as loading control. Representative immunoblot from two independent biological experiments is shown. D, NPQ kinetics of Col-0, *soq1*, TNC, TM-TNC and T+NC transformants. Five-week-old plants grown at 120 μmol photons m^−2^ s^−1^ were dark acclimated for 20 min and NPQ was induced at 1,200 μmol photons m^−2^ s^−1^ (white bar) and relaxed in the dark (black bar). Data represent mean ± SD (n = 8 to 12; four individuals from two or three independent T2 or T4 lines).

The role of redox regulation in the thylakoid lumen is understood for only a few proteins, and the reducing or oxidizing partners of these redox-regulated proteins are not all known (Hoh et al., 2023). Recent studies have identified thiol-disulfide exchange reactions involving HCF164 for reduction of target proteins such as cytochrome *f* whose thiol is required for heme assembly (Motohashi and Hisabori, 2006), and lumen thiol oxidoreductase 1 (LTO1) for oxidation of target proteins such as PSII subunit O (PsbO) (Karamoko et al., 2011) or STN7 (Wu et al., 2021) whose disulfides enable proper folding for activity. Exploring whether SOQ1 lumenal domains are involved in redox regulation could provide insights into the broader redox network in the thylakoid lumen.

Understanding SOQ1 functional role not only reveals its regulatory function in photosynthetic light harvesting efficiency but also holds implications for human health. NHL repeat-containing protein 2 (NHLRC2), a mammalian homolog of SOQ1, shares structural similarities and potential functional roles (Biterova et al., 2018; Nishi et al., 2017; Yu et al., 2022). Variants of NHLRC2 are associated with the fatal FINCA (fibrosis, neurodegeneration, cerebral angiomatosis) disease in early childhood (Uusimaa et al., 2018; Hiltunen et al., 2020; Badura-Stronka et al., 2022; Tallgren et al., 2023) and with neural tube defects in bovines (Nicholas et al., 2022). NHLRC2 is involved in phagocytosis and cytoskeleton rearrangement, but its biochemical function is yet to be determined (Paakkola et al., 2018; Haney et al., 2018; Yeung et al., 2019).

The structure of SOQ1 lumenal domains unveiled an additional domain (CTD), which we found indispensable for suppressing qH (Yu et al., 2022). Importantly, SOQ1 domains have homology to disulfide bond protein D (DsbD). DsbD, a bacterial inner membrane protein, comprises three domains: nDsbD which has similarity to SOQ1-CTD, a central transmembrane domain (tDsbD), and cDsbD which has similarity to SOQ1-Trx (Yu et al., 2022). Each domain contains two catalytic cysteines that enable electron transfer through a cascade of thiol–disulfide exchange reactions, from tDsbD to cDsbD and finally to nDsbD (Missiakas et al., 1995; Stewart et al., 1999; Katzen and Beckwith, 2000). DsbD then transfers reducing power to downstream proteins such as DsbC, DsbG, and CcmG, which promote protein folding (Missiakas et al., 1994; Shevchik et al., 1994; Rietsch et al., 1996), oxidative stress defense (Depuydt et al., 2009; Denoncin et al., 2014), and c-type cytochrome maturation (Fabianek et al., 1998; Stirnimann et al., 2005). Remarkably, SOQ1 cysteine pairs (C431-C434 in Trx-like domain and C1006-C1012 in the CTD) structurally align with those of *Escherichia coli* DsbD (C461-C464 in cDbsD and C103-C109 in nDsbD), suggesting potential thiol-disulfide exchange between SOQ1 lumenal domains that may be important for SOQ1 function. The higher NPQ and slower relaxation, coupled with the slower electrophoretic mobility of LCNP, in the *soq1* knock-out mutant compared to wild-type plants, indicate that LCNP is post-translationally modified when SOQ1 does not function, and this modification activates LCNP for qH formation (Malnoë et al., 2018). However, the mechanism by which LCNP activates for qH formation and how SOQ1 inhibits LCNP to suppress qH is not known.

In this study, we aimed to investigate the function of SOQ1 lumenal domains in their soluble form, the potential thiol-disulfide exchange between SOQ1-Trx and CTD domains and their role in qH regulation. Our findings demonstrate that SOQ1 lumenal domains can function as soluble and separate entities, are redox-active and possess methionine sulfoxide reductase activity, a process not previously recognized in the lumen of chloroplast. We conclude that LCNP methionine oxidation acts as a signal for qH induction, providing another elegant way for cells to activate energy dissipation using oxidative stress for feedback.

## Results

### SOQ1 lumenal domains in their soluble form can prevent qH

We previously found that truncated forms of SOQ1 can accumulate in the thylakoid membrane (TM-TNC) and lumen (TNC, NC and CTD) in both non-stress conditions (growth light (GL)) and stress conditions (6 h cold (4□) and 1,600 µmol photons m^−2^ s^−1^ high light (CHL)) (Yu et al., 2022). Next, we investigated whether these truncated forms are functional by transforming constructs that encode TM-TNC, TNC, Trx (T) or NC separately or together (T+NC) into the *soq1* mutant background (Fig.1A). Wild type listed as Col-0 (Columbia accession) was used as a reference in the following experiments. To target these SOQ1 variants into the thylakoid lumen, the chloroplast and lumenal targeting sequences of PsbO were fused to the N-terminus of TNC, T or NC. We isolated three independent lines for TM-TNC and TNC, and two for T, NC and T+NC. We confirmed that the variants accumulated (Fig. 1B, 1C) and localized in the chloroplast thylakoid (TM-TNC) or were properly targeted to the lumen (TNC, T, NC, T+NC) in non-stress and stress conditions (Fig. S1A, S1B). Moreover, TM-TNC, TNC and NC can be cleaved between domains yielding NC and CTD accumulating in the lumen and also CTD’ (Fig. S1A, S1B), which is a smaller form of CTD with a possible different cleavage site in the NC loop (Yu et al., 2022). All the forms were detected using antibodies against the C-terminal peptide of SOQ1 (anti-SOQ1_CTD_), except for SOQ1-Trx to which we added a Strep-tag to its C-terminus for immunodetection. SOQ1-Trx was also found in several forms possibly due to different processing of its C-terminal loop and accumulated to some extent in the membrane fraction (Fig. S1B).

To test the quenching suppression capacity of these variants, we measured chlorophyll fluorescence of leaves under high light (1,200 μmol photons m^−2^ s^−1^) for 10 min followed by 10 min in the dark from plants grown under non-stress conditions. This high light intensity triggers qH in the *soq1* mutant observed by a higher NPQ induction in the light followed by a slower relaxation in the dark (Fig. 1D). We confirmed that the transformants had similar maximum fluorescence (F_m_) to *soq1* (Fig. S2A, S2B, Table S1) and compared their NPQ kinetics. Notably, both TM-TNC and TNC were able to rescue wild-type NPQ kinetics (Fig. 1D), indicating that the soluble TNC is sufficient to prevent qH formation. In addition, the electrophoretic mobility of LCNP in TM-TNC and TNC was the same as in wild type (Fig. 1B), further supporting that the slower mobility of LCNP in the *soq1* mutant contributes to qH formation.

SOQ1-Trx (T) or NC expressed separately failed to rescue wild-type NPQ kinetics (Fig. S1C) and the mobility of LCNP in Trx and NC remained the same as in *soq1* (Fig. 1C), suggesting that both Trx and NC are required to inhibit LCNP. Indeed, the T+NC plants that co-expressed Trx and NC displayed an intermediate level of NPQ between that of wild type and *soq1* which is statistically different from *soq1* (Fig. 1D, Table S2). Interestingly, the electrophoretic mobility of LCNP in T+NC plants was also intermediate and partially restored (Fig. 1C).

To examine whether these variants can also function under stress conditions, we performed a 6h-CHL treatment which induces qH in wild type (Malnoë et al., 2018). TM-TNC and TNC displayed a similar NPQ level as wild type whereas T, NC and T+NC reached similar NPQ level as *soq1* (Fig. S1D, S1E, Table S2). Although T+NC could inactivate some LCNP under non-stress conditions, that was insufficient under cold and high light conditions likely due to a high level of LCNP (Levesque-Tremblay et al., 2009). Taken together, these results demonstrate that SOQ1 lumenal domains (Fig. 1D) in their soluble form (TNC, and T+NC to some extent) can prevent qH possibly by restoring LCNP modification (indicated by the rescued electrophoretic mobility) thereby inactivating LCNP.

### SOQ1 Trx domain and CTD are redox active and the redox state of SOQ1 is light-dependent

The structural homology of SOQ1 Trx and CTD domains to cDsbD and nDsbD, respectively, raises the question of whether Trx and CTD are redox active. To address this question, we tested the redox activity of the recombinant Trx (SOQ1-Trx) and CTD (SOQ1-CTD) purified from *E. coli*. We probed the redox state of SOQ1-Trx and SOQ1-CTD in the presence of different concentrations of the reductant dithiothreitol (DTT), or the oxidant diamide, using the 4-acetamido-4′-maleimidylstilbene-2,2′-disulfonic acid (AMS) labeling assay. AMS is a 0.5 kDa thiol-reactive agent that can alkylate reduced cysteine residues, resulting in a slower migration on SDS-PAGE of the reduced protein compared to the oxidized protein if the cysteines are involved in a disulfide bond (Inaba et al., 2005). After AMS labeling, both SOQ1-Trx and SOQ1-CTD displayed slower mobility under reducing conditions (band annotated as “Red” for reduced) compared to oxidizing conditions (band annotated as “Ox” for oxidized) (Fig. 2A), indicating that one disulfide bond can be formed in SOQ1-Trx and SOQ1-CTD, respectively. The slower mobility of the Ox band in the AMS-labeled control (Cont) compared to the non-treated sample (-AMS) is due to an additional lone cysteine present in each domain. Indeed, Trx-*f*1, a protein containing a cysteine pair involved in a disulfide bond and an additional lone cysteine (Yoshida et al., 2015), was purified and probed for reference, and showed a similar banding pattern as Trx and CTD. Additionally, we followed the redox state of SOQ1-Trx or SOQ1-CTD along a concentration gradient of reducing and oxidizing DTT (Fig. 2B) and calculated their midpoint redox potential (*Em*) values as −274 mV and −252 mV (pH 7.0, 25□), respectively (Fig. 2C), implying that SOQ1-Trx could transfer electrons to SOQ1-CTD.

**Fig. 2.**
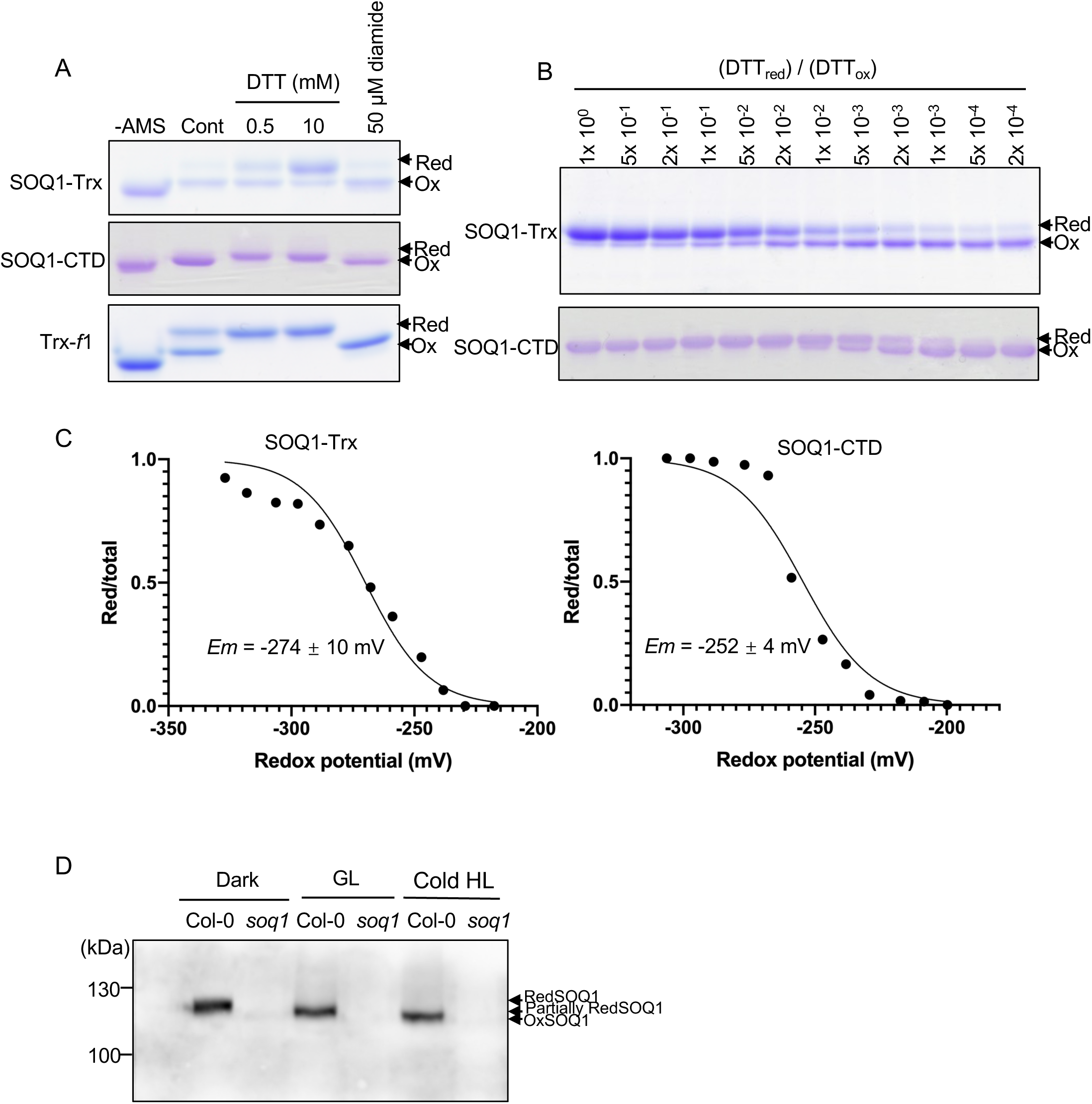
The cysteines of SOQ1-Trx and CTD are redox active *in vitro* and SOQ1 is oxidized under stress conditions *in vivo*. A, AMS labeling of recombinant SOQ1-Trx or CTD under different redox conditions: no add (Cont), reducing (DTT) or oxidizing (diamide). Thioredoxin-*f*1 (Trx-*f*1) was used as a positive control. Proteins were TCA-precipitated, labeled with AMS, loaded on a nonreducing SDS-PAGE, and stained with CBB. The reduced (Red) form migrates slower than the oxidized (Ox) form due to AMS modification of reduced cysteines. Difference in MW between unlabeled (-AMS) and the oxidized form is due to the labeling of free non redox-active cysteines. B, Trx and CTD were equilibrated with various ratios of redox buffers (reduced DTT (DTTred))/(oxidized DTT (DTTox)). C, The midpoint redox potential of SOQ1-Trx and SOQ1-CTD. The reduction level quantified as the ratio of the reduced form to the total was plotted against the redox potential of the DTT buffer. The data were fitted to the Nernst equation. The Em values are the mean of three (SOQ1-Trx) and five (SOQ1-CTD) independent experiments. D, Total Arabidopsis proteins were rapidly isolated using SDS buffer with or without 8 mM MM(PEG)24 to label reduced cysteines from Col-0 and *soq1* plants grown under standard growth conditions and exposed to dark (samples were taken at the end of the night period), growth light (GL) for 6 h or a 6 h-cold and high light treatment (Cold HL). The samples were loaded at the same amount of total proteins (30 µg) on a 8% non-reducing SDS-PAGE and analyzed by immunodetection using anti-SOQ1_CTD_ antibody. Representative immunoblot from three independent biological experiments is shown.

The presence of cysteine pairs in SOQ1-Trx and CTD which can form disulfides prompted us to analyze the redox state of SOQ1 *in vivo*. We chose methyl-PEG-Maleimide (MM(PEG)24) to alkylate reduced cysteine residues due to its larger size (1.2 kDa), which makes it better suited for proteins with high molecular weight. We labelled whole cell proteins to investigate SOQ1 full-length redox state and not lumenal proteins for the truncated forms, as the redox state of lumenal proteins could be changed during the long isolation procedure. In the wild type, SOQ1 migration is slower in non-stress conditions (dark and growth light) compared to stress conditions (cold and high light), indicating that SOQ1 is reduced in non-stress conditions and oxidized in stress conditions (Fig. 2D).

### SOQ1 Trx domain exhibits disulfide reductase activity

We determined that SOQ1-Trx and SOQ1-CTD are redox-active, and we hypothesized that thiol-disulfide exchange occurs between these domains. To test this hypothesis, we first attempted a classical insulin turbidity assay but did not observe reduction of insulin by SOQ1-Trx (Fig. S3A,B). We then performed electron transfer assays between reduced SOQ1-Trx (obtained by treatment with 100 mM DTT for 1 h at 25 °C, followed by DTT removal) and its likely native substrate instead, oxidized SOQ1-CTD (obtained by treatment with 50 mM CuCl_2_ for 1 h at 25 °C, followed by CuCl_2_ removal), at different temperatures (25□, 37□ and 42□) with a low concentration of DTT to function as electron donor. We determined that 0.2 mM DTT was an appropriate concentration as it only leads to partial reduction (∼20 %) of SOQ1-CTD (Fig. 3A, C). We monitored the redox state of SOQ1-CTD upon addition of DTT alone or upon incubation with DTT and different concentrations of reduced SOQ1-Trx (1-12 μM) and found that SOQ1-CTD can be reduced by SOQ1-Trx (Fig. 3B, C, S3C). In addition, we tested the reverse reaction using different concentrations of reduced SOQ1-CTD (0.5-4 μM) (obtained by treatment with 100 mM DTT for 1 h at 25 °C, followed by DTT removal) and oxidized SOQ1-Trx (obtained by treatment with 50 mM CuCl_2_ for 1 h at 25 °C, followed by CuCl_2_ removal; 10 % of Trx is in the Red form and 0.2 mM DTT leads to ∼20 % Trx Red) and found that the ratio of reduced SOQ1-Trx to total SOQ1-Trx did not change between addition of DTT alone or together with reduced SOQ1-CTD (Fig. 3D, E, S3D). These results suggest that electron transfer occurs between SOQ1-Trx and SOQ1-CTD and that the electron flow is unidirectional from SOQ1-Trx to SOQ1-CTD.

**Fig. 3.**
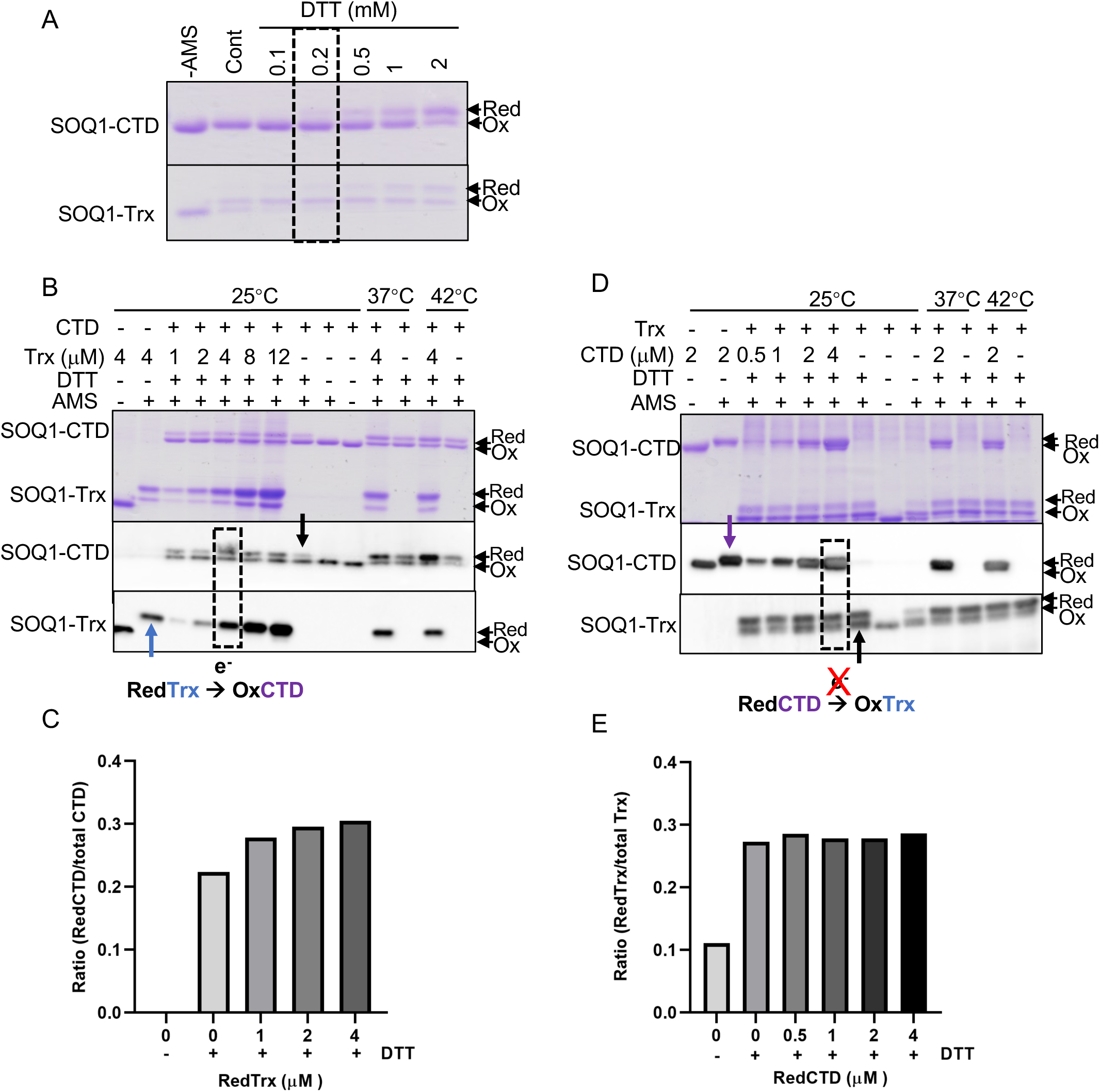
Electron transfer between SOQ1 Trx and its CTD. A, Determination of the maximum amount of DTT to mediate the electron transfer reaction. 2 µM SOQ1-CTD or SOQ1-Trx was treated with different concentrations of DTT (0-2 mM) at 25°C for 10 min. B, Electron transfer reaction assay between partially reduced SOQ1-Trx (blue arrow) and oxidized SOQ1-CTD using DTT as an electron donor. 2 µM oxidized SOQ1-CTD was incubated with different concentrations of reduced SOQ1-Trx (1-12 µM) in the presence of 0.2 mM DTT at 25°C, 37°C and 42°C for 10 min. The reaction was stopped by TCA precipitation and samples were labeled with AMS, loaded on a nonreducing SDS-PAGE stained with CBB (upper, 200 pmol samples were loaded) or transferred and analyzed by immunodetection (0.2 pmol samples were loaded) using anti-SOQ1_CTD_ (middle) or anti-His for SOQ1-Trx (bottom). DTT alone leads to a small reduction of CTD (black arrow) whereas addition of both DTT and Trx reduces CTD to a higher amount (dashed box). Representative gel and immunoblot from three independent biological experiments are shown. C, The ratio of reduced SOQ1-CTD (RedCTD) to the total (RedCTD + OxCTD) is calculated from the CBB gel in B. D, Electron transfer reaction assay between reduced SOQ1-CTD (purple arrow) and partially oxidized SOQ1-Trx using DTT as an electron donor. 2 µM partially oxidized SOQ1-Trx was incubated with various concentration of reduced SOQ1-CTD (0.5, 1, 2 and 4 µM) in the presence of 0.2 mM DTT at 25°C, 37°C and 42°C for 10 min. The method is the same as in B, here middle panel is using anti-His to probe for SOQ1-CTD (same results with anti-SOQ1_CTD_). The redox state of Trx does not change between addition of DTT alone (black arrow) or together with reduced CTD (dashed box). The anti-His antibody affinity for the AMS-labeled Ox form of SOQ1-Trx is lower than that of the Red form of SOQ1-Trx for unknown reasons (same result with anti-SOQ1_Trx_). Representative gel and immunoblot from two independent biological experiments are shown. E, The ratio of reduced SOQ1-Trx (RedTrx) to the total (RedTrx + OxTrx) calculated from CBB gel in D.

### Two cysteines in CTD are required for qH suppression

Previously, we established that the CTD and the conserved cysteines (C431 and C433) in the Trx domain were essential for SOQ1 to suppress qH (Yu et al., 2022; Brooks et al., 2013). Using structural homology to nDsbD, we determined that C1006 and C1012 in CTD may be involved in a disulfide bond (Yu et al., 2022) making them good candidates for the thiol-disulfide exchange reaction with SOQ1-Trx. However, three cysteines are present in SOQ1-CTD (Yu et al., 2022) namely C1001, C1006 and C1012. To assess which cysteine residues are required for the suppression function of SOQ1, we transformed the *soq1* mutant with constructs containing either cysteine to serine mutation (C1006S, C1012S) or cysteine to threonine (C1001T). We verified that the mutated proteins were expressed and localized at the thylakoid membrane and observed similar cleavage of lumenal domains as in Col-0 (Fig. 4A, S1, S4). The maximum fluorescence F_m_ values of the transformants were similar to *soq1* (Fig. S2C, Table S1). Interestingly, C1006S and C1012S displayed similar NPQ kinetics to *soq1*, whereas C1001T rescued wild-type NPQ kinetics (Fig. 4B). After a cold and high light-treatment, the NPQ level in C1006S and C1012S reached similar values as *soq1*, whereas C1001T displayed wild-type NPQ level (Fig. 4C). Furthermore, C1001T restored LCNP mobility, whereas C1006S and C1012S exhibited slower mobility of LCNP as in *soq1* (Fig. 4A). These results demonstrate that the two cysteine residues, C1006 and C1012, are required for qH suppression, whereas C1001 is dispensable for SOQ1 function.

**Fig. 4.**
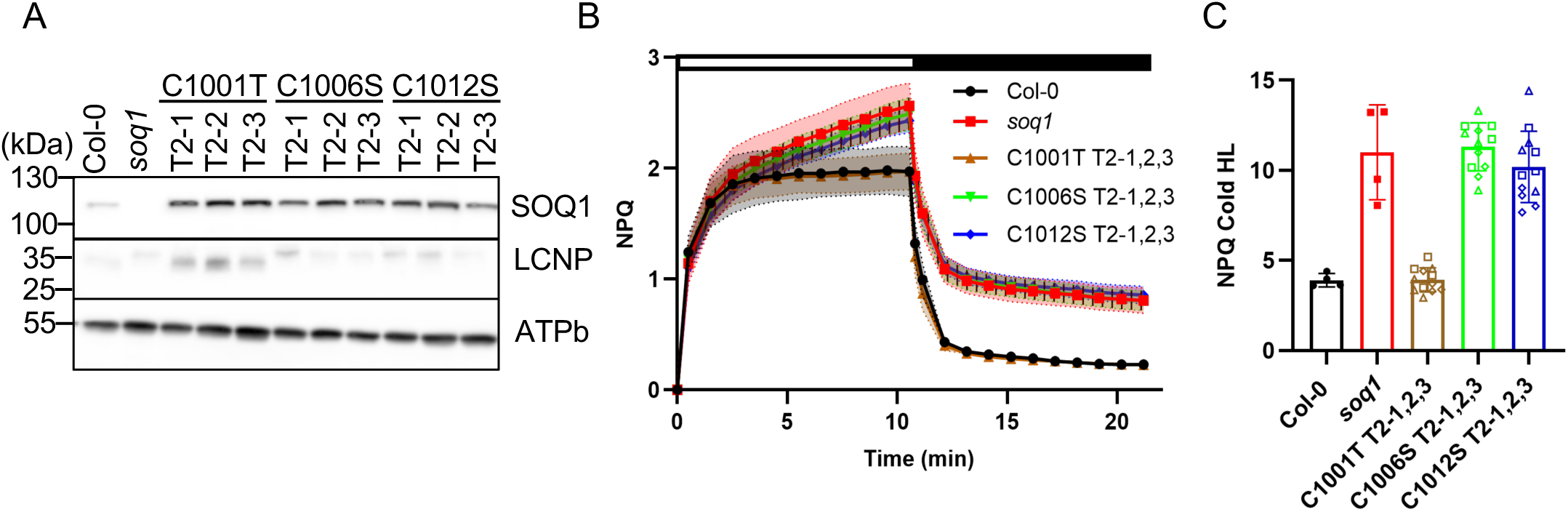
C1006 and C1012 in CTD are required for qH suppression. A, Protein accumulation in Col-0, *soq1* and three individuals from independent T2 lines transformed in *soq1* mutant background with full length SOQ1 containing mutation C1001T, C1006S or C1012S. 15 µg total leaf proteins were loaded. Proteins were separated by SDS-PAGE and analyzed by immunodetection with antibodies against SOQ1, LCNP or ATPb. ATPb is shown as loading control. B, NPQ kinetics of Col-0, *soq1* and C1001T, C1006S and C1012S transformants. Five-week-old plants grown at 150 μmol photons m^−2^ s^−1^ were dark-acclimated for 20 min and NPQ was induced at 1200 μmol photons m^−2^ s^−1^ (white bar) and relaxed in the dark (black bar). C, NPQ levels of Col-0, *soq1* and C1001T, C1006S and C1012S transformants after 6h CHL treatment calculated as (Fm before treatment – Fm after treatment)/(Fm after treatment); Fm after treatment was measured following dark-acclimation for 5 min. (B-C) Data represent means ± SD (n = 12; four individuals from three independent T2 lines with square, triangle and diamond symbols denoting −1, −2 and −3 respectively).

### Recombinant TNC possesses methionine sulfoxide reductase activity

On reducing SDS-PAGE followed by immunoblot, we have observed that LCNP electrophoretic mobility is slower in the *soq1* mutant. Decreased mobility on reducing SDS-PAGE can be due to methionine oxidation to methionine sulfoxide and sulfone or cysteine oxidation to cysteine sulfenic, sulfinic and sulfonic acid (Liang et al., 2012). LCNP contains 13 methionine residues and 7 cysteine residues (6 of which are involved in disulfide bridges, Hao et al. in preparation). LCNP mobility is not restored by addition of reducing reagents (Malnoë et al., 2018); therefore a different content in disulfides or cysteine sulfenic acids is not causal for this behavior (Rudyk and Eaton, 2014), so we hypothesized that the slower mobility of LCNP is due to methionine oxidation. We purified and incubated recombinant LCNP with N-ethyl maleimide (NEM) to alkylate free cysteine residues followed by treatment with hydrogen peroxide (H_2_O_2_) to oxidize methionine residues (Costa et al., 2007; Muthuramalingam et al., 2013). The recombinant LCNP_L87_ (starting at Leucine 87, mature protein predicted by Target P-2.0) was used, which contains the seven conserved cysteines. The oxidation efficiency of H_2_O_2_ was titrated (Fig. S5A), and finally the optimum condition was determined as 50 mM of H_2_O_2_ for three days at 4 in which a slower mobility of LCNP, reminiscent of that observed in the *soq1* mutant, was now apparent (Fig. 5A, Fig. S5D).

**Fig. 5.**
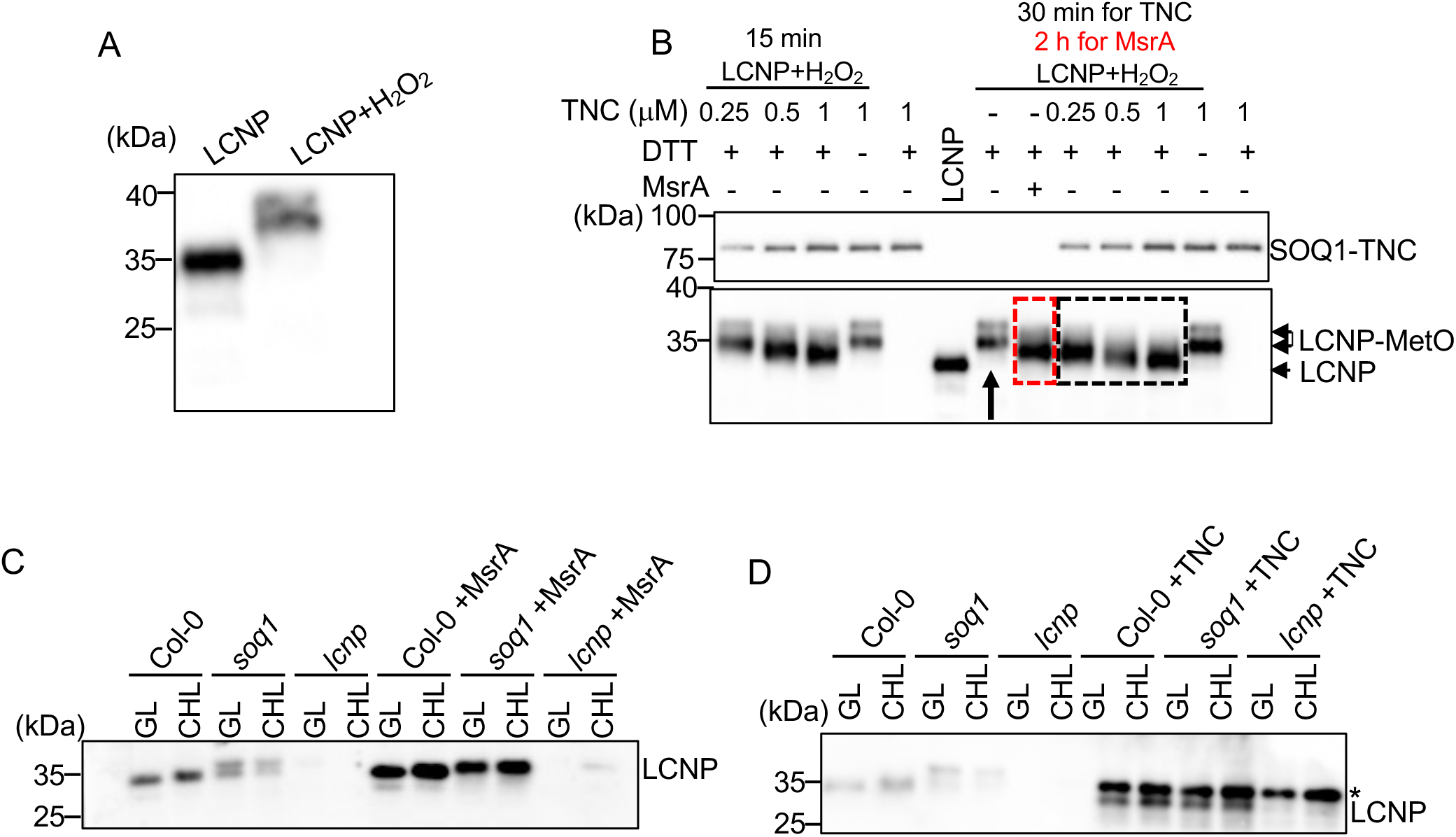
LCNP electrophoretic mobility is restored upon addition of methionine sulfoxide reductase or TNC. A, H_2_O_2_-treatment of recombinant LCNP. Immunoblot analysis of purified recombinant LCNP_L87_ treated with NEM to alkylate free cysteines (LCNP) followed by treatment with H_2_O_2_ (LCNP+H_2_O_2_). B, Addition of methionine sulfoxide reductase A (MsrA) or SOQ1-TNC to recombinant LCNP. 1 µM LCNP+H_2_O_2_ (black arrow) displays a slower mobility compared to LCNP which is restored upon incubation with different amount of TNC in the presence of 5 mM DTT at 25℃ for 15 or 30 min (black dashed box) or with 3.8 µM MsrA at 37℃ for 2 h (red dashed box). The reaction was stopped by TCA precipitation. The samples were separated by SDS-PAGE and analyzed by immunodetection using anti-LCNP and anti-SOQ1_CTD_ antibodies. Representative immunoblot from two independent biological experiments is shown. C, Incubation of lumenal proteins with MsrA. 30 µg lumenal proteins from Col-0, *soq1* and *lcnp* grown under standard growth conditions and exposed to growth light (GL) or cold high light (CHL) conditions were incubated with or without 3 µM recombinant MsrA for 2h at 37℃. D, Incubation of lumenal proteins with 4 µM recombinant TNC for 30 min at room temperature under 5 mM DTT. Proteins were denatured by adding 1x SDS sample buffer and analyzed by immunodetection using anti-LCNP antibody. Star symbol (*) represents a nonspecific band, based on its presence in *lcnp,* detected by the anti-LCNP antibody likely originating from the TNC recombinant protein preparation.

To test whether the methionines in LCNP are oxidized by H_2_O_2_, we incubated LCNP+H_2_O_2_ with 4 μM recombinant methionine sulfoxide reductase A (MsrA) from human in the presence of 5 mM DTT for 2 h at 37 □ and found that LCNP mobility was partially restored (Fig. 5B). In parallel, we incubated LCNP+H_2_O_2_ with increasing concentrations of SOQ1-TNC in the presence of 5 mM DTT for 15 min or 30 min at 25 □ and found that LCNP mobility was fully restored with 1 μM TNC (Fig. 5B). Similar results were obtained using the recombinant LCNP_A104_ (starting at Alanine 104 based on a previous prediction of the mature protein by Target P) which does not contain the first conserved cysteine and without NEM (Fig. S5B). We then examined LCNP slower mobility *in vivo* by incubating the lumen fraction from Col-0 and *soq1* isolated under non-stress or stress conditions with recombinant MsrA and found that LCNP mobility was partially restored (Fig. 5C). Strikingly, the slower mobility of LCNP in *soq1* was fully restored by adding recombinant TNC (Fig. 5D, S5C). These results strongly suggest that LCNP slower mobility is due to methionine oxidation and that SOQ1-TNC possesses methionine sulfoxide reductase activity.

### The methionines of LCNP are required for inactivation by SOQ1

To investigate further whether the slower mobility of LCNP in *soq1* is due to oxidized methionine residues, we generated LCNP constructs replacing methionine by alanine. Of the 13 methionine residues in Arabidopsis LCNP (AtLCNP), seven are highly conserved (M125, M126, M129 in the N-terminal domain, M219, and M309, M310, M315 in the C-terminal domain). An enrichment in dsed methionines in both N-terminal and C-terminal domains seems to be a feature of LCNP proteins; AtLCNP contains a total of seven methionines in its N-terminal domain (M_121_MMMMMRGM_129_) as well as four methionines in its C-terminal domain (M_309_MSMPGM_315_) (Fig. 6A). We generated two LCNP variants: one with ten Met-to-Ala mutations (all the N- and C-terminal domain methionines but for M129 that we were unable to obtain) named 10MTA (M121 122 123 124 125 126 309 310 312 315) and one with twelve Met-to-Ala mutations named 12MTA (10MTA plus M172 and M219). We transformed these LCNP constructs in both *lcnp* and *soq1 lcnp* mutants to ensure LCNP mobility could be compared (i.e. not solely due to a change of molecular weight from the introduced alanines) and isolated three independent lines for each. We observed a slightly slower mobility of LCNP in the *soq1 lcnp* LCNP-10MTA lines compared to the *lcnp* LCNP-10MTA lines, whereas LCNP mobility no longer differed between the *soq1 lcnp* LCNP-12MTA lines and the *lcnp* LCNP-12MTA lines (Fig. 6B, Fig. S6A). Cold and high light treatment did not lead to further slowing down of LCNP mobility in the MTA lines (Fig. S6C,D). These results clearly indicate that LCNP slower mobility in *soq1* is due to methionine residues, possibly several of them in their oxidized form.

**Fig. 6.**
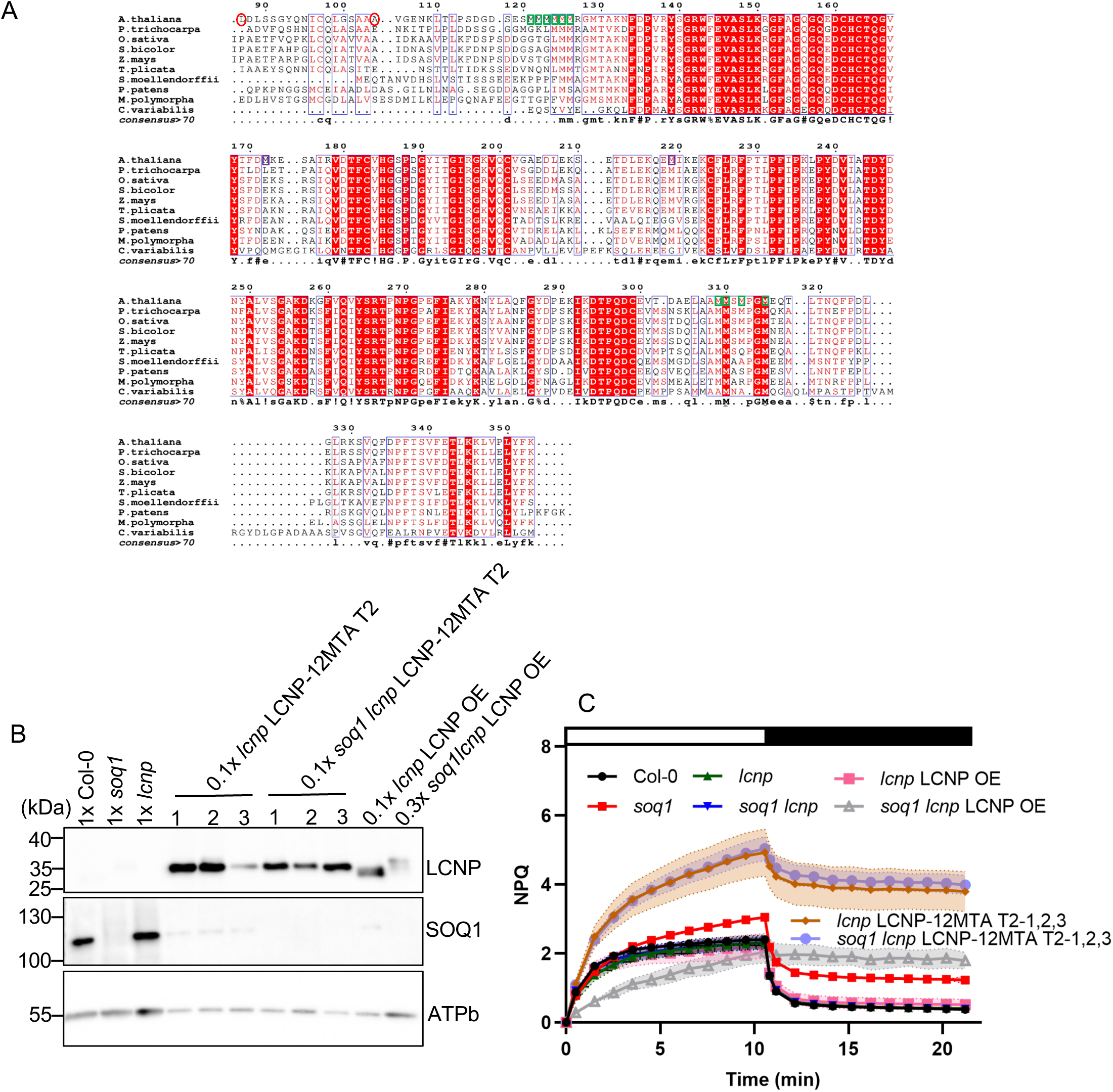
Methionines of LCNP are required for inactivation by SOQ1. A, Multiple sequence alignment of LCNP proteins in photosynthetic eukaryotes. It includes mature LCNP protein sequences from *Arabidopsis thaliana* (At3g47860), *Populus trichocarpa* (Potri.001G420800), *Oryza sativa* (Os04g53490), *Sorghum bicolor* (Sobic.006G222700), *Zea mays* (Zm00001d002240), *Thuja plicata* (Thupl.29377109s0015), *Selaginella moellendorffii* (98590), *Physcomitrium patens* (Pp6c9_3320), *Marchantia polymorpha* (Mapoly0115s0028) and *Chlorella variabilis* (CHLNCDRAFT_145104). A 100% conservation is marked with red background. The 10 mutated methionines are marked with a green box and the additional two in the 12 mutated mutant with a purple box. L87 and A104 are marked with a red circle. B, Immunoblot analyses of three independent T2 *Arabidopsis* variant LCNP lines containing 12 methionine to alanine (MTA) mutations transformed in *lcnp* and *soq1 lcnp* mutant background. LCNP overexpressed in *lcnp* (*lcnp* LCNP OE) and *soq1 lcnp* (*soq1 lcnp* LCNP OE) mutant background were used as the LCNP mobility control. Samples were loaded at the same amount of total protein (1x corresponds to 15 µg), less proteins were loaded for the transformants as LCNP is overexpressed. ATPb was used as the loading control. C, NPQ kinetics of Col-0, *soq1,* three independent *lcnp* and *soq1 lcnp* T2 variant lines, *lcnp* LCNP OE and *soq1 lcnp* LCNP OE lines. Five-week-old plants grown at 150 μmol photons m^−2^ s^−1^ were dark-acclimated for 20 min and NPQ was induced at 1,200 μmol photons m^−2^ s^−1^ (white bar) and relaxed in the dark (black bar). Data represent means ± SD (n = 12; four individuals from three independent T2 lines).

The maximum fluorescence F_m_ values of the transformants were similar to Col-0 or *lcnp* LCNP OE (Fig. S7, Table S1), but the NPQ levels of the *lcnp* LCNP-MTA lines were approximately two times higher than the controls and *soq1* as well (Fig. 6C, Fig. S6B). The methionine mutations to alanine in LCNP therefore results in a more active form of LCNP. In addition, the *soq1 lcnp* LCNP-MTA lines now showed similar NPQ to the *lcnp* LCNP-MTA lines, so the lack of SOQ1 no longer led to additional NPQ (Fig. 6C, Fig. 6B). These data suggest that the methionine residues of LCNP are required for its inactivation by SOQ1; in other words, SOQ1 cannot prevent qH formation upon high light exposure if the methionines of LCNP are mutated. Of note, the *soq1 lcnp* LCNP OE has a largely decreased F_m_ (80% lower than Col-0) and F_o_, whereas *soq1 lcnp* LCNP-MTA does not (Fig. S7A,B). This “pre-quenched” F_m_ can likely be explained by the overexpression of LCNP which can become activated in growth light as SOQ1 is missing. This is consistent with the high levels of the slower mobility form of LCNP observed in *soq1 lcnp* LCNP OE (Fig. 6B and Fig. S6A). In contrast the MTA variants require high light intensity to become activated.

## Discussion

Here, we have uncovered that the lumenal domains of SOQ1 in their soluble form can prevent qH (Fig. 1), that both Trx and CTD are redox-active and that the *in vivo* redox state of SOQ1 is light-dependent with a more oxidized state upon light stress (Fig. 2). We demonstrated that the SOQ1 Trx domain can reduce CTD *in vitro* (Fig. 3), and that two cysteine residues in the CTD are required for the suppression of qH (Fig. 4). Importantly, we found that the recombinant lumenal domains of SOQ1 possess methionine sulfoxide reductase (Msr) activity (Fig. 5), and that the methionine residues of LCNP are required for its inactivation by SOQ1 (Fig. 6).

### Dithiol-disulfide exchange between SOQ1 Trx and CTD

The structural similarities between DsbD and SOQ1 lumenal domains indicated that dithiol-disulfide exchange may occur between SOQ1 Trx and CTD (Yu et al., 2022). We provide experimental evidence that a cysteine pair can form a disulfide in both SOQ1-Trx and SOQ1-CTD (Fig. 2A). Because C1006 and C1012 but not C1001 were essential to prevent qH in complementation assays (Fig. 4), we conclude that C431 and C434 (Brooks et al., 2013) and C1006 and C1012 are the cysteines pairs involved in disulfide bond formation in SOQ1 Trx and SOQ1 CTD, respectively. We measured the midpoint redox potential of SOQ1-Trx (*Em* = −274 mV) and SOQ1-CTD (*Em* = −252 mV) at pH 7.0 (Fig. 2B, 2C) (of note, as lumenal pH acidifies in the light (down to <6.0), these potentials may become more positive (Hoh et al., 2023) and determined that dithiol-disulfide exchange occurs between these two domains in the direction of increasing potential from SOQ1-Trx to SOQ1-CTD (Fig. 3, S3). In contrast, *Neisserial* cDsbD (*Em* = −242mV) and nDsbD (*Em* = −246mV), as well as *E. coli* cDsbD (*Em* = −235 mV) and nDbsD (*Em* = −232mV) display similar midpoint potentials (Rozhkova et al., 2004; Smith et al., 2019). The difference in redox potential may govern the unidirectional electron transfer between SOQ1-Trx and SOQ1-CTD, whereas electron transfer in DsbD is based on the exposure of the active site in the oxidized nDsbD, which contains the Phenylalanine (F66) cap in an open conformation; if nDsbD is reduced, the cap is closed to prevent reverse electron transfer (Smith et al., 2019). Interestingly, CTD does contain a conserved Phenylalanine (F950), which might correspond to F66 (Yu et al., 2022) and could serve to regulate substrate(s) access to CTD to prevent a futile redox cycle. Whether F950 is involved in forming an open conformation to expose the active site of CTD needs more investigation.

### Regulation of SOQ1 activity

SOQ1 lumenal domains in their soluble form (TNC, and even T+NC to some extent) can prevent qH (Fig. 1, S1), suggesting that the stromal and transmembrane (TM) domains are not required for SOQ1 function in suppressing qH. This result is at variance with Brooks et al. (2013) in which the ΔHAD/TM construct did not complement, but this is likely due to mistargeting; here we used the lumenal targeting sequence from PsbO to ensure lumenal localization. The implication is that unlike DsbD periplasmic domains which obtain reducing power from the transmembrane domain tDsbD (Katzen and Beckwith, 2000), SOQ1 lumenal domains would receive electrons from another protein. A good candidate is cytochrome c deficiency A (CcdA), a homolog of tDsbD (Katzen et al., 2002; Page et al., 2004; Motohashi and Hisabori, 2010). Remarkably, the suppression capacity of soluble TNC was similar to that of TM-TNC and Col-0, indicating that TNC is able to accept electrons at a similar rate from the thylakoid membrane, despite not being anchored by its TM domain. When TNC is further separated into its T and NC domains (T+NC), the suppression capacity is much lower (Fig. 1, S1), so T and NC can interact in the lumen as truncated forms, but the efficiency of electron transfer is decreased. Interestingly, the redox state of SOQ1 *in vivo* changes from reduced to oxidized upon stress (Fig. 2D) indicating an imbalance in the flux of electrons with possibly less electrons coming from the stromal donors to CcdA and/or more electrons going to SOQ1 target(s). Future work will focus on identifying the stromal electron donors and how they may transfer less electrons to SOQ1 so that LCNP can be activated under stress.

### Possible catalytic mechanism of SOQ1

Methionines and cysteines contain sulfur atoms which are prone to oxidation. The sulfoxide form of methionine (MetO) and sulfenic and sulfinic forms of cysteine (CysSOH, CysSO_2_H) are reversible whereas the further oxidized derivative (sulfone, MetO_2_ and sulfonic acid, CysSO_3_H) are not (Costa et al., 2007). Two enantiomers of MetO can form upon oxidation depending on their configuration in the protein: S- and R-MetO, and they can be reversed by specific reductases: methionine sulfoxide reductase A (MsrA) and MsrB, respectively (Lee and Gladyshev, 2011). The treatment of LCNP with H_2_O_2_ led to oxidation of its methionines (alkylation of cysteines prevented their oxidation) and upon incubation with MsrA, LCNP electrophoretic mobility was partially restored (Fig.5, S5) indicating that some of LCNP-MetO was in the S-form and reversed (a mixture of both epimers should form upon H_2_O_2_ treatment (Toennies and Kolb, 1939)). Strikingly, TNC completely restored LCNP mobility suggesting that SOQ1 possesses methionine sulfoxide reductase activity with either higher efficiency than MsrA possibly due to LCNP being its native substrate, and/or it may be that SOQ1 possesses both A and B activity.

Typically, Msr proteins obtain reducing power from thioredoxins (Brot et al., 1981) or glutaredoxins (Vieira Dos Santos et al., 2007) and these reductants are separate proteins from the Msr protein. The PilB protein present in some bacteria, however, contains both activities with its Trx-like N-terminal domain reducing the central MsrA and C-terminal MsrB domains (Wu et al., 2005; Brot et al., 2006; Boschi-Muller, 2018). Similarly, here we found that the Trx-like domain of SOQ1 provides reducing power to the CTD displaying Msr activity. MsrA and MsrB can form a single fused protein (MsrAB) in many bacteria increasing the catalytic efficiency compared to the separated MsrA and MsrB (Han et al., 2016; Kim et al., 2009). In the case of SOQ1, there is no clear structural homology between CTD and Msr, as it is similar instead to nDsbD. Could nDsbD also display Msr activity? nDsbD is known as an electron donor to PilB (Quinternet et al., 2009) but to our knowledge has not been tested for Msr activity. Although it may be unlikely that nDsbD possesses Msr activity as only a few residue changes could have led to the Msr activity of SOQ1-CTD. Indeed a few residue changes can convert the sulfenic acid reductase activity of DsbG (structurally similar to DsbC) into a disulfide bond isomerase (Chatelle et al., 2015). The β-stranded domain of SOQ1 CTD would constitute a novel fold for Msr activity. The reaction mechanism of SOQ1 CTD Msr activity includes two cysteines, C1006 and 1012 (Fig. 4), which may function similarly to a subclass of MsrA or to MsrB (Rouhier et al., 2007; Lowther et al., 2002; Lourenço Dos Santos et al., 2018). It would proceed as follows: one cysteine performs a nucleophilic attack on the methionine sulfoxide substrate; the generation of methionine leads to a sulfenic intermediate on that cysteine, which reacts with the second cysteine forming an intramolecular disulfide, and the disulfide would then be reduced by the Trx-like domain of SOQ1.

NHLRC2, the mammalian homolog of SOQ1, did not display thioredoxin activity in the insulin turbidity assay (Uusimaa et al., 2018; Yeung et al., 2019) but neither did cDsbD (Smith et al., 2019) or SOQ1 (Fig. S3), therefore it is also possible that NHLRC2 Trx-like domain can perform dithiol-disulfide exchange with its CTD. Indeed, the CCINC motif of NHLRC2 and SOQ1 Trx-like domains adopt almost identical conformations (Yu et al., 2022), and two cysteine residues in the CTD of NHLRC2, C689 and C696, are conserved (Biterova et al., 2018) and would correspond to C1006 and C1012 of SOQ1, respectively. Next, it will be exciting to explore whether NHLRC2 exhibits Msr activity and determine if the FINCA disease is due to dysregulation of its targets by MetO.

In Arabidopsis, the Msr family consists of 14 members which have a role in both protection against oxidative damage and signaling; none are however localized to the thylakoid lumen (Rey and Tarrago, 2018). SOQ1 is therefore the first known enzyme of the thylakoid lumen to display Msr activity. The source of ROS generating MetO in the lumen under excess light could be singlet oxygen (^1^O_2_) or H_2_O_2_ (Triantaphylidès and Havaux, 2009; Khorobrykh et al., 2020; Li and Kim, 2022) as a two-electron oxidation is reversible (Schöneich, 2005). Oxidation could be passive but it may be too slow to be biologically relevant, and likely not stereospecific, although it could explain that qH in wild type takes several hours of intense cold temperature and light stress to be induced (Malnoë et al., 2018). Alternatively, it may be enzymatically-assisted by monooxygenases (Manta and Gladyshev, 2017) but none have been characterized in the lumen (Järvi et al., 2013). An intriguing possibility may be that SOQ1 also possesses methionine oxidase activity similarly to what has been found for mouse MsrA (Lim et al., 2011) where the availability of Trx regulates the oxidation or reduction function of MsrA i.e. if the Trx does not have access to MsrA then MsrA functions as an oxidase. That may be the case under certain conditions (especially as truncated T and NC domains accumulate (Yu et al., 2022)), however LCNP-MetO forms in absence of SOQ1 under standard conditions so its oxidation can occur without SOQ1.

### Methionine oxidation for LCNP activation

The electrophoretic mobility of LCNP is slower in the *soq1* mutant compared to wild type and this is linked to higher qH (Malnoë et al., 2018). Here we have consistently observed a correlation between the slower mobility of LCNP and the presence of qH (*soq1* transformed with SOQ1-Trx, -NC, -C1006S or -C1012S (Fig. 1, S1, 4)) suggesting that LCNP is activated through a post-translational modification. We found that H_2_O_2_-treated LCNP slows down its electrophoretic mobility similarly to the *in vivo* migration of LCNP in the *soq1* mutant (Fig. 5, S5D). Incubation with MsrA or SOQ1-TNC partially or completely restores the migration of LCNP both *in vivo* and *in vitro* (Fig. 5), and the LCNP-12MTA line rescued LCNP mobility fully (Fig. 6B). Altogether these results demonstrate that the slower mobility is methionine-dependent and that LCNP-MetO would be the activated form to induce qH. Which exact methionine residues of LCNP are oxidized to MetO and how oxidation activates LCNP will be the focus of future work. The size difference in SDS electrophoresis of approximately 2 kDa between the oxidized and non-oxidized form is more pronounced than expected if all 13 Met are oxidized to MetO (+208 Da). The slower migration on SDS-PAGE is a general feature of methionine-rich proteins and likely a consequence of protein conformational alterations due to methionine oxidation modifying the ability of SDS to bind and/or denature the protein (Vougier et al., 2004; Liang et al., 2012). Methionine oxidation has generally been associated with the loss of protein function for which Msr enzymes play a role in the repair of oxidized proteins, but it has now been increasingly recognized as a mode of protein activation (Drazic and Winter, 2014). Here, LCNP is a new example of a protein function modulated by the redox status of its methionine residues, with SOQ1 as a dedicated enzyme for the reduction of MetO in LCNP to regulate its function.

Of note, the LCNP-MTA lines resulted in a form of LCNP no longer inhibited by SOQ1 (Fig. 6C, Fig. S6B), the methionines of LCNP are therefore required for its inactivation by SOQ1. However in contrast to the wild type form of LCNP overexpressed in the *soq1* background, the MTA variants required high light for their activation which merits further investigation. Indeed the nitrate-dependent transcription factor NirA is activated by oxidation of Met169 and the loss of hydrophobic interaction from the substitution of methionine to alanine mimicking the more hydrophilic MetO in a M169A mutant rendered it into a permanently active form (Gallmetzer et al., 2015). Next, substitution of methionines to isoleucines to mimic reduced methionines may prevent activation of LCNP.

Our study elucidated the redox-regulation mechanism of photoprotective qH by SOQ1 on LCNP (Fig. 7), providing novel insights into the molecular interplay between these proteins within the thylakoid lumen. Specifically, we have demonstrated that SOQ1 suppresses qH by inhibiting LCNP through its lumenal domains, exhibiting both thioredoxin disulfide reductase and methionine sulfoxide reductase activities that convert activated LCNP-MetO to its inactive LCNP-Met form (Fig. 7). These findings advance our understanding of the photoprotective mechanisms that safeguard photosynthetic efficiency highlighting a feedback regulation of light harvesting through oxidation of methionine residues.

**Fig. 7.**
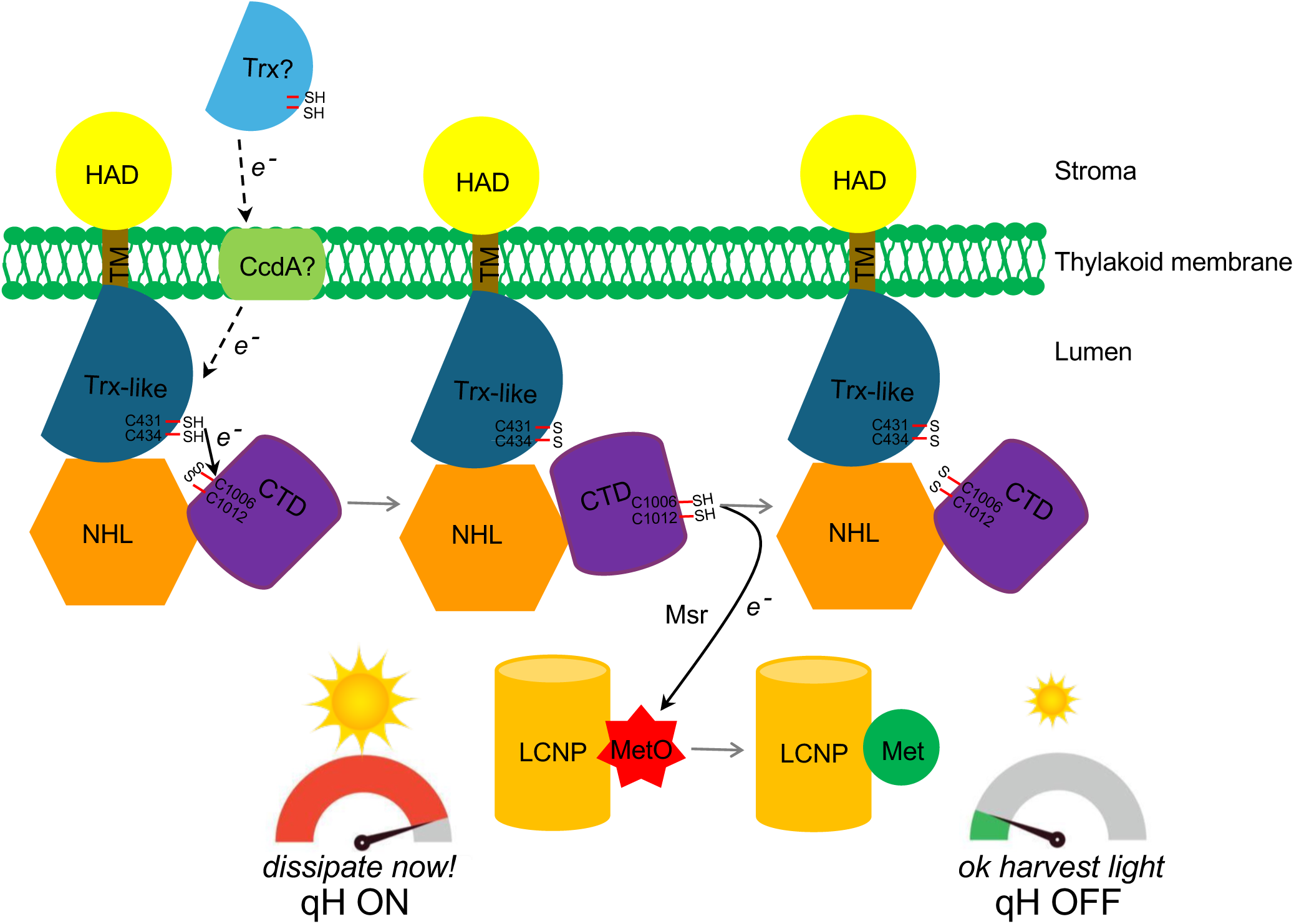
Model for SOQ1 regulation of qH-energy dissipation: LCNP-methionine oxidation as an oxidative stress meter. The Trx-like domain of SOQ1 may obtain electrons (dashed arrows) from reduced proteins in the stroma, such as thioredoxins, through the membrane protein, cytochrome c deficiency A (CcdA). The reduced Trx-like domain transfers electrons to the CTD and the reduced CTD displays methionine sulfoxide reductase (Msr) activity which converts LCNP-MetO to LCNP-Met (solid arrows, showed in this work). Under non-stress conditions, SOQ1 maintains LCNP-Met reduced. Under stress conditions, accumulation of LCNP-MetO enables LCNP activation for qH. Adapted from (Brooks et al., 2013) and (Yu et al., 2022).

## Materials and methods

### Plant material and growth conditions

Wild-type *Arabidopsis thaliana* and the derived mutants studied here are of the Col-0 ecotype. Knock-out (KO) mutant *soq1* is *soq1-1* described by Brooks et al., (2013) and mutant *lcnp* is T-DNA KO *lcnp-1* described by Levesque-Tremblay et al., (2009). All plant materials in this study were grown in a growth chamber with 120 μmol photons m^−2^ s^−1^ light intensity and 60% humidity at 20 °C during the day for 8 h and 18 °C during the night for 16 h. For cold and high light treatment, the plants were illuminated in the cold room (4 °C) for 6 h at 1,600 µmol photons m^−2^ s^−1^ light intensity using a custom-designed LED panel built by JBeamBio with cool white LEDs BXRA-56C1100-B-00 (Farnell).

### Cloning

The cDNAs encoding SOQ1 Trx-NHL-CTD (TNC, K390-R1055), Trx-like domain (Trx, K390-L535), and CTD (E898-R1055) were amplified from pET151/D-TOPO-SOQ1 vector (Brooks et al., 2013) through PCR. Then the PCR products were cloned into pET151/D-TOPO vector that contains at the N-terminal 6xHis tag DNA sequence following the manufacturer’s instructions. The TNC, Trx, and CTD constructs were transformed into *Escherichia coli* strain BL21 (DE3) for protein purification. Rare codons of *LCNP* cDNA without the sequence encoding the chloroplast transit peptide were mutated to codons that are frequently used in *E. coli* and synthesized in pUC57 vector (GenScript USA Inc.). *LCNP* cDNA encoding LCNP_L87_ and LCNP_A104_ were amplified from pUC57-LCNP and cloned into pET24d vector with a N-terminal 10xHis-SUMO-Flag-fusion tag. The construct expressing LCNP was transformed into *E. coli* strain Origami (DE3).

### Protein expression and purification

The clones were cultured in LB medium containing 100 µg·ml^−1^ ampicillin for TNC, Trx and CTD or 50 µg·ml^−1^ kanamycin for LCNP at 37 °C and shaken at 220 rpm until the OD600 reached 0.2-0.3, then the protein expression was induced by adding 1 mM IPTG overnight at 21 °C. Cells were harvested by centrifugation at 5,500g for 15 min at 4 °C. The pellets were resuspended in 3 ml of lysis buffer (25 mM Tris-HCl, pH7.5, 10 mM imidazole for TNC, Trx and CTD, 50 mM Tris-HCl, pH7.5, 10 mM imidazole for LCNP) and disrupted by sonication (20 s on, 10 s off, 45 cycles). After centrifugation at 17,000g for 30 min at 4 °C, the supernatant was harvested and incubated with 1 ml of pre-washed ProBond Nickel Cleaning Resin (Invitrogen, Cat. No. R80101) for 1h at 4 °C. After washing successively with 50 ml of 20 mM, 15 ml of 40 mM and 15 ml of 60 mM imidazole in lysis buffer (for LCNP purification, 50 ml of 20 mM imidazole for washing), protein was eluted with 5 ml of lysis buffer containing 250 mM imidazole. The protein was concentrated with Amicon Ultra-15 Centrifugal Filter (Sigma Aldrich, Cat. No. UFC900308) and the concentration was measured using the Bradford assay.

### Redox characterization of Trx and CTD *in vitro*

To determine whether Trx and CTD each contain one disulfide bond, 2 μM recombinant Trx and CTD were respectively incubated with and without DTT (0.5 mM and 10 mM) and 20 μM diamide in 50 mM phosphate buffer (pH 7.0). The total reaction volume is 200 μl. After incubation for 1 h at 25 °C, proteins were precipitated with 10 % (v/v) trichloroacetic acid (TCA) and washed two times with 100 % ice-cold acetone. Precipitates were dissolved in 20 μl of 1x non-reducing SDS buffer (62.5 mM Tris-HCl (pH 6.8), 2 % (w/v) SDS, 7.5 % (v/v) glycerol and 0.01 % (w/v) bromophenol blue) containing 2 mM of thiol-modifying reagent AMS (Invitrogen, Cat. No. A485). After labeling for 1 h at 25 °C in the dark, protein samples were loaded onto 12 % nonreducing SDS-PAGE and stained with Coomassie Brilliant Blue R-250 (ThermoFisher, Cat. No. 24615).

The putative disulfide reduction activity of SOQ1-Trx domain was tested using similar conditions as described previously (Motohashi and Hisabori, 2006). Reduction of insulin is measured as a change in turbidity of the solution due to precipitation of free insulin B chain (Holmgren, 1979). Insulin reduction assay was carried out at 25 °C in a 1.6 ml reaction mixture containing 50 mM potassium phosphate (pH 7.0), 2 mM EDTA, 130 µM bovine insulin and 1, 5 or 10 µM recombinant protein. The reaction was initiated by adding 500 µM DTT and the change in turbidity was monitored at 650 nm. Nonenzymatic reduction of insulin by DTT was monitored in the absence of added recombinant protein. Arabidopsis stromal Trx-m1 was used as a positive control (Yoshida et al., 2015).

To study the electron transfer process between Trx and CTD, each protein was treated with 100 mM DTT or 50 μM CuCl_2_ at 1h at 25 to obtain the reduced and oxidized forms. The 100 mM DTT or 50 μM CuCl_2_ reagents were removed by running through a PD-10 column (GE Healthcare, Cat. No. GE17-0851-01). The optimum concentration of DTT was determined by incubating the different concentrations of DTT (0.1, 0.2, 0.5, 1 and 2 mM) with oxidized CTD or oxidized Trx for 10 min. Reduced Trx (1, 2, 4, 8, 12 μM) was incubated with oxidized CTD (2 μM) and oxidized Trx (2 μM) was incubated with reduced CTD (0.5, 1, 2, 4 μM) in 50 mM potassium phosphate buffer (pH 7.0) and 200 μM DTT for 10 min incubation at 25 °C, 37 °C and 42 °C. The total reaction volume above is 200 μl. Proteins were TCA-precipitated, labeled with AMS, and loaded onto a nonreducing SDS-PAGE and immunodetection analysis using the rabbit-specific antibodies against a C-terminal peptide of SOQ1 (anti-SOQ1_CTD_), a peptide of SOQ1 Trx-like domain (anti-SOQ1_Trx_) (Malnoë et al., 2018; Yu et al., 2022) or anti-His tag (R&D Systems, Cat. No. MAB050H).

### Determination of the midpoint redox potential of Trx and CTD

The midpoint redox potential of Trx and CTD were determined as described previously (Motohashi and Hisabori, 2006; Yoshida et al., 2015) with modifications. Recombinant protein (1 µM) was incubated in 50 mM potassium phosphate buffer (pH 7.0), 50 mM oxidized DTT, and various concentrations of reduced DTT (10 µM to 50 mM). After incubation for 3 h at 25 °C, proteins were precipitated with 10 % (v/v) TCA and washed with ice-cold acetone, alkylated with AMS and subjected to nonreducing SDS-PAGE as described above. The *E_m_* values of both proteins were calculated by fitting the titration data of the reduction level of proteins to the Nernst equation. A value of −327 mV was used as the *E_m_*of DTT at pH 7.0 (Singh et al., 1995).

### Determination of SOQ1 redox state *in vivo*

The protein redox state *in vivo* was determined as described by Yoshida and Hisabori (Yoshida and Hisabori, 2019) with modifications. Col-0, *soq1* and *lcnp* plants were grown in a growth chamber until five-week-old. For dark conditions, samples were harvested before the light was turned on in the growth chamber. For standard light conditions, samples were harvested after 6 h light illumination at 120 µmol photons m^−2^s^−1^. For cold and high light treatment, the plants were illuminated in the cold room (4 °C) for 6 h at 1600 µmol photons m^−2^ s^−1^. Sample collection was performed as follow: two fully expanded leaves were excised and immediately ground using liquid nitrogen in a mortar with a pestle. The powdered plant leaves were dissolved in 300 μl of 1 x non-reducing SDS buffer with or without 8 mM methyl-PEG-maleimide reagent (Thermo Scientific, Cat. No. 22713). After incubation at 25 °C for 1 h in the dark with shaking, protein samples were heated at 95 °C for 5 min. After centrifugation at 14,000 rpm for 10 min, the supernatant was transferred to a new 1.5 ml Eppendorf tube. The protein concentration was measured by Pierce™ BCA Protein Assay Kit (Thermo Scientific, Cat. No. 23225). 30 µg of proteins were loaded on an 8 % SDS-PAGE coupled with immunoblot analysis using a rabbit-specific antibody against the C-terminal peptide of SOQ1 (anti-SOQ1_CTD_) (Malnoë et al., 2018).

### Generation and identification of *Arabidopsis* transformants

*C1001T*, *C1006S*, *C1012S*, *TNC*, *Trx* and *NC* cDNAs were obtained using site-specific PCR mutagenesis (Follo and Isidoro, 2008) using pENTR/D-TOPO-SOQ1 C-Flag vector as template (Brooks et al., 2013). These constructs were cloned either into pEarleyGate100 or pEarleyGate100 that was modified by adding a N-terminal cDNA encoding the chloroplast and lumenal transit peptides of *Arabidopsis* photosystem II subunit O (PsbO, At5g66570), which target TNC, Trx and NC to the thylakoid lumen. LCNP methionine mutation constructs were obtained via QuickChange Lightning Multi Site-Directed Mutagenesis kit (Agilent, Cat. No. 210513) and cloned into the pCambia1300 under the ubiquitin 10 promoter with a C-terminal Flag tag of LCNP. All the primers used for this study are listed in Table S3. The constructs above were transferred into *soq1* mutants (SOQ1 constructs) or *lcnp* and *soq1 lcnp* (LCNP methionine mutation constructs) by the floral dip method (Clough and Bent, 1998) using *Agrobacterium tumefaciens GV3101* cell cultures. The wild-type LCNP overexpression (OE) lines are described in Hao et al. in preparation. T1 seeds were selected on Murashige and Skoog (MS) plates containing 15 µg·ml^−1^ basta or 30 μg·ml^−1^ hygromycin. Selected plants were propagated to the T2 generation and used for experiments. The protein accumulation of the transformants was determined by immunoblot analysis using anti-SOQ1_CTD_ (Malnoë et al., 2018), anti-Flag (Sigma-Aldrich, Cat. No. F7425), anti-Strep (Agrisera, Cat. No. AS21 4682) or rabbit-specific antibody against a peptide of LCNP (Yu et al., 2022).

### Total protein extraction

Around 100 mg of leaves from five-week-old plants were ground in liquid nitrogen with a mortar and pestle. 400 μl of extraction buffer (50 mM HEPES pH 7.2, 150 mM NaCl, 15 mM MgCl2, 1 mM EDTA, 10 % glycerol, 1 % Triton X-100, 10 mM β-mercaptoethanol, 2 mM PMSF and Roche cOmplete Protease Inhibitor Cocktail)) were added in the ground leaves and incubated for 30 min at 4 °C. After centrifugation at 14,000 rpm for 15 min, the supernatant was collected to measure the protein concentration using Bradford reagent.

### Immunoblot analysis

Samples were loaded at the same total protein, chlorophyll or lumenal protein, separated on 8, 12 or 15 % SDS–PAGE, transferred to PVDF membrane (Sigma-Aldrich, Cat. No. IPVH00005), blocked with 5 % (w/v) non-fat dry milk and incubated for 1 h with the following antibodies: anti-SOQ1_CTD_, (1:200 dilution) (Malnoë et al., 2018), against a peptide of SOQ1 Trx-like domain (anti-SOQ1_Trx_) (GVHSAKFDNEKDLDAIR) produced by ThermoFisher (1:1,000 dilution), and against a peptide of LCNP (AEDLEKSETDLEKQ) produced and purified by peptide affinity by Biogenes (1:200 dilution) (Yu et al., 2022), His-Tag HRP-conjugated antibody (1:1,000 dilution, R&D Systems, Cat. No. MAB050H), mouse anti-Strep-tag (1:1,000 dilution, Agrisera, Cat. No. AS21 4682), anti-light harvesting complex b4 (Lhcb4) (1:7,500 dilution, Agrisera, Cat. No. AS04 045), anti-ATP synthase subunit beta (ATPb) (1:2,000 dilution, Agrisera, Cat. No. AS05 085) and anti-plastocyanin (PC) (1:2,000 dilution, Agrisera, Cat. No. AS06 141). (HRP)-conjugated anti-rabbit (diluted 1:10,000; Sigma-Aldrich, Cat. No. A6154) or anti-mouse (diluted 1:10,000; Agrisera, Cat. No. AS11 17724) secondary antibodies were used for detection by chemiluminescence with ECL substrate (Agrisera, Cat. No. AS16 ECL-N-100).

### Chlorophyll fluorescence measurement

Chlorophyll fluorescence was measured at room temperature from detached, fully expanded leaves using a fluorescence imaging system Speedzen 200 (JBeamBio). Leaves were dark-acclimated for 20 min and NPQ was induced using 1,200 µmol photons m^−2^ s^−1^ (red actinic light) for 10 min and relaxed in the dark for 10 min. Maximum fluorescence levels after dark acclimation (F_m_) and throughout measurement (F_m_’) were recorded after applying a saturating pulse of light. NPQ was calculated as (F_m_-F_m_’)/F_m_’. NPQ level after cold high light treatment was calculated as (F_m_ before treatment-F_m_ after treatment)/F_m_ after treatment.

### Chloroplast sub-fraction isolation

Chloroplast sub-fractions were isolated as described by Yu et al. (2022) with modifications. Briefly, 10 g leaves from 6-week-old wild type, *soq1*, *lcnp* and SOQ1 variants were blended in 90 ml of extraction buffer containing 50 mM Tricine-NaOH pH 7.8, 330 mM sorbitol, 1 mM EDTA, 10 mM KCl, 0.15% (w/v) bovine serum albumin, 4 mM sodium ascorbate and 7 mM L-cysteine, two times for 5 sec using a Multi Blender (Coline). Homogenate was collected by filtrating through 4 layers miracloth (Cat. No.475855, Calbiochem) and centrifuged at 3,000 rpm for 5 min at 4 °C. After washing with the resuspension buffer (50 mM sodium phosphate pH 7.8, 330 mM sorbitol, 10 mM NaF), chloroplasts were ruptured by osmotic shock in lysis buffer (10 mM sodium phosphate pH 7.8, 5 mM MgCl_2_, 10 mM NaF). Following centrifugation at 7,500 rpm for 5 min at 4 °C, thylakoids were collected in the pellets. The thylakoids were washed briefly with washing buffer (50 mM sodium phosphate pH 7.8, 100 mM sorbitol, 10 mM NaF, 5 mM MgCl_2_) and resuspended in sonication buffer (30 mM sodium phosphate (pH 7.8), 50 mM NaCl, 5 mM MgCl2 and 100 mM sucrose) at a chlorophyll concentration of 0.5 mg ml^−1^. Intact thylakoids were broken by sonication method adapted from (Levesque-Tremblay et al., 2009) (Sonics Vibra cell VCX130). Thylakoid membrane and lumen were separated by ultracentrifugation for 2 h at 50,000 rpm at 4 °C. The lumen proteins were concentrated at 14,000 rpm for 10 min at 4 °C using Amicon ultra-0.5 centrifugal filters (Cat. No. UFC500324, Merck millipore). All solutions contained protease inhibitors with final concentration of 1 mM benzamidine, 5 mM ε-aminocaproic acid and 0.2 mM PMSF and/or Roche cOmplete™ protease inhibitor cocktail tablet. See also (Hao and Malnoë, 2023) for detailed protocol.

### Determination of SOQ1-TNC activity *in vitro*

The recombinant LCNP and TNC were purified as described above. The free thiols of ∼1 mg of LCNP were blocked using 100 mM N-ethyl maleimide (NEM) (Thermo Fisher, Cat. No. 23030) in darkness for 1 h (Muthuramalingam et al., 2013). After removing the excess NEM with PD-10 desalting columns, LCNP was treated with 50 mM H_2_O_2_ for 3 days at 4 °C to obtain LCNP-MetO. The excess H_2_O_2_ was removed by PD-10 desalting columns and LCNP-MetO was concentrated using Amicon ultra-0.5 centrifugal filters. 1 μM LCNP-MetO was incubated with 0.25, 0.5 and 1 μM TNC or 3.8 μM MsrA in the presence or in the absence of 5 mM DTT at 25 °C for 15 min and 30 min or 2 h (MsrA). The total reaction volume is 200 μl. The reaction was stopped by adding 10 % TCA for protein precipitation. After two times wash by 100 % acetone, the pellets were resuspended with 1x SDS sample buffer (containing 100 mM DTT) and analyzed by immunodetection using anti-LCNP and anti-SOQ1_CTD_ antibodies.

### Recombinant TNC and MsrA-dependent reaction

MsrA (Human) recombinant protein was purchased from Abnova company (Cat. No. P4550). 3 μM MsrA or 4 μM TNC was incubated with 30 μg lumenal proteins of wild type, *soq1* and *lcnp* plants in the presence of 5 mM DTT for 2 h at 37 (MsrA) and 30 min at 25 (TNC). The same amount of lumenal proteins without adding recombinant proteins was used as the control. The total reaction volume is 50 μl. The reaction was stopped by adding SDS sample buffer containing 62.5 mM Tris-HCl pH 6.8, 2 % SDS, 0.002 % Bromophenol Blue, 10 % glycerol, 100 mM DTT. Proteins were loaded in 12 % SDS-PAGE as the same amount and analyzed by immunodetection using anti-LCNP antibody.

### Multiple sequence alignment

Putative orthologs of AtLCNP were obtained from Phytozome (www.phytozome.net) or NCBI using a BLAST search. The results were manually curated for correctness of the gene models and exclusion of non-orthologous genes. Plastid and lumenal transit peptides were predicted by TargetP2.0 (Almagro Armenteros et al., 2019) and not included in the alignment. The protein sequences were aligned using Clustal Omega from EMBL-EBI (Madeira et al., 2024) and visualized using the ESPript 3.0 server (Robert and Gouet, 2014).

## Data availability

The authors declare that all data supporting the findings in this study are included in the paper and its supplementary information file and are available from the corresponding author upon request. Source data for all figures are provided with the paper. Sequence data from this article can be found in the Arabidopsis Genome Initiative under accession numbers At1g56500 (SOQ1) and At3g47860 (LCNP).

## Acknowledgement

We thank the Japanese Photosynthesis Consortium for funding A.M.’s stay in the Hisabori Lab at Tokyo Tech in Japan and Félix Buchert for his suggestion that LCNP may display methionine oxidation. We thank Mikael Lindberg for assistance in plasmid construction and protein expression, Pierrick Bru for technical assistance, and Aurélie Crepin, Maria Paola Puggioni and Pierrick Bru for critical discussions and Ulrich Bergmann for critical reading of the manuscript. The project was funded by European Commission Marie Skłodowska-Curie Actions Individual Fellowship (to A.M.) Reintegration Panel (845687). This research (J.H and A.M.) was supported by grants to UPSC from the Knut and Alice Wallenberg Foundation (2016.0341 and 2016.0352), the Swedish Governmental Agency for Innovation Systems (2016-00504), by a starting grant to A.M. from the Swedish Research Council Vetenskapsrådet (2018-04150) and by a consortium grant from the Swedish Foundation for Strategic Research (ARC19-0051). A.P.H., M.D.B., and K.K.N. were supported by the U.S. Department of Energy, Office of Science, through the Photosynthetic Systems program in the Office of Basic Energy Sciences. K.K.N. is an investigator of the Howard Hughes Medical Institute. A.P.H. received funding by Deutsche Forschungsgemeinschaft (DFG, German Research Foundation) under Germanýs Excellence Strategy – EXC-2048/1 – project ID 390686111.

## Author contributions

J.H. and A.M. designed the research; J.H. performed research with assistance from A.P.H., A.J. and J.S.F. for midpoint redox potential and technical support from A.M., K.Y. and T.H. for AMS labeling assay. A.M. and M.D.B. constructed the plasmids encoding truncated SOQ1 forms and J.H. constructed the plasmids encoding Cys-mutated SOQ1. A.M. generated the TM-TNC line and J.H. generated other transgenic lines of SOQ1. J.H. and J.D. constructed the plasmids encoding Met-mutated LCNP and generated the transgenic lines. All the authors discussed the data, and J.H. and A.M. wrote the paper with input from all authors.

