## Supplemental Figures for "Suppressor of quenching 1 functions as a methionine sulfoxide reductase in the chloroplast lumen for regulation of photoprotective qH in Arabidopsis"

**Running title: The methionine sulfoxide reductase activity of SOQ1**

Jingfang Hao[a], Alexander Johansson[a], Johan Svensson Fall[a], Jianli Duan[a], Alexander P. Hertle[bcdef], Matthew D. Brooks[bcg], Krishna K. Niyogi[bcd], Keisuke Yoshida[h], Toru Hisabori[h], Alizée Malnoë[ai]*

**Supplemental Data**: Fig. S1 to S7 and Table S1 to S3

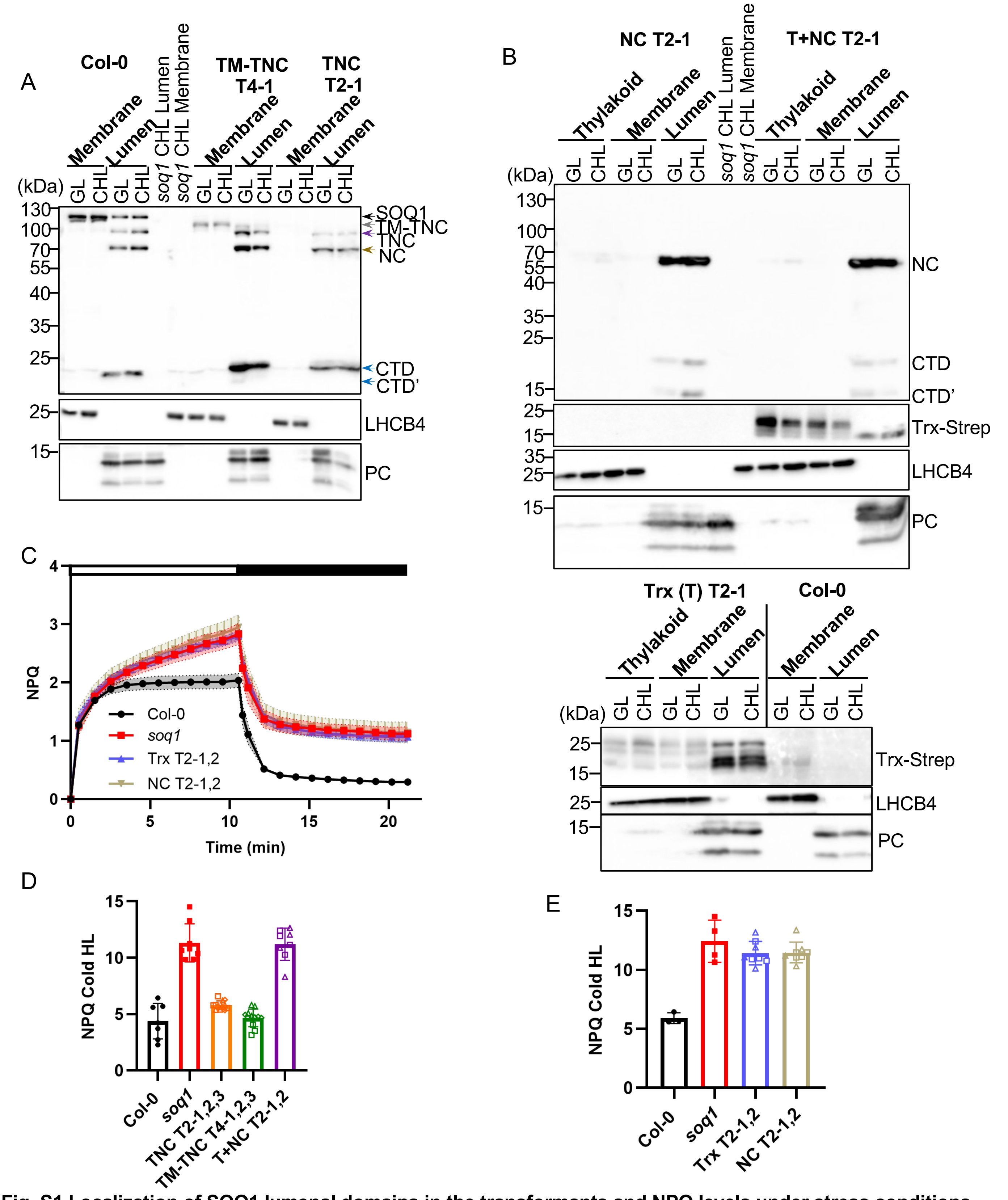

**Fig. S1 Localization of SOQ1 lumenal domains in the transformants and NPQ levels under stress conditions.** A, B, Localization of TNC, TM-TNC, Trx, NC and T+NC in chloroplast sub-fractions (Membrane: thylakoid membrane after having separated the lumen fraction, and Lumen) in growth light (GL) and after 6h cold and high light treatment (CHL). Samples were loaded at the same amount of chlorophyll (1.5 µg) for thylakoid and membrane or lumenal proteins (5 µg). Proteins were separated by SDS-PAGE and analyzed by immunodetection with antibodies against SOQ1-CTD, Strep-tag, LHCB4 (thylakoid membrane marker) and PC (lumen marker). Representative immunoblots from two independent biological experiments are shown. C, NPQ kinetics of Col-0, *soq1*, Trx and NC lines. Five-week-old plants grown at 120 µmol photons $m^{-2}$ $s^{-1}$ were dark acclimated for 20 min and NPQ was induced at 1,200 µmol photons $m^{-2}$ $s^{-1}$ (white bar) and relaxed in the dark (black bar). Data represent mean ± SD (n = 8; four individuals from two independent T2 lines). D, E, NPQ levels of Col-0, *soq1*, TNC, TM-TNC, T+NC, Trx and NC plants after 6h CHL treatment. NPQ was calculated as (Fm before treatment – Fm after treatment)/(Fm after treatment); Fm after treatment was measured following dark-acclimation for 5 min. Data represent means ± SD (n = 8 to 12; four individuals from two or three independent T2 or T4 lines with square, triangle and diamond symbols denoting -1, -2 and -3 respectively).

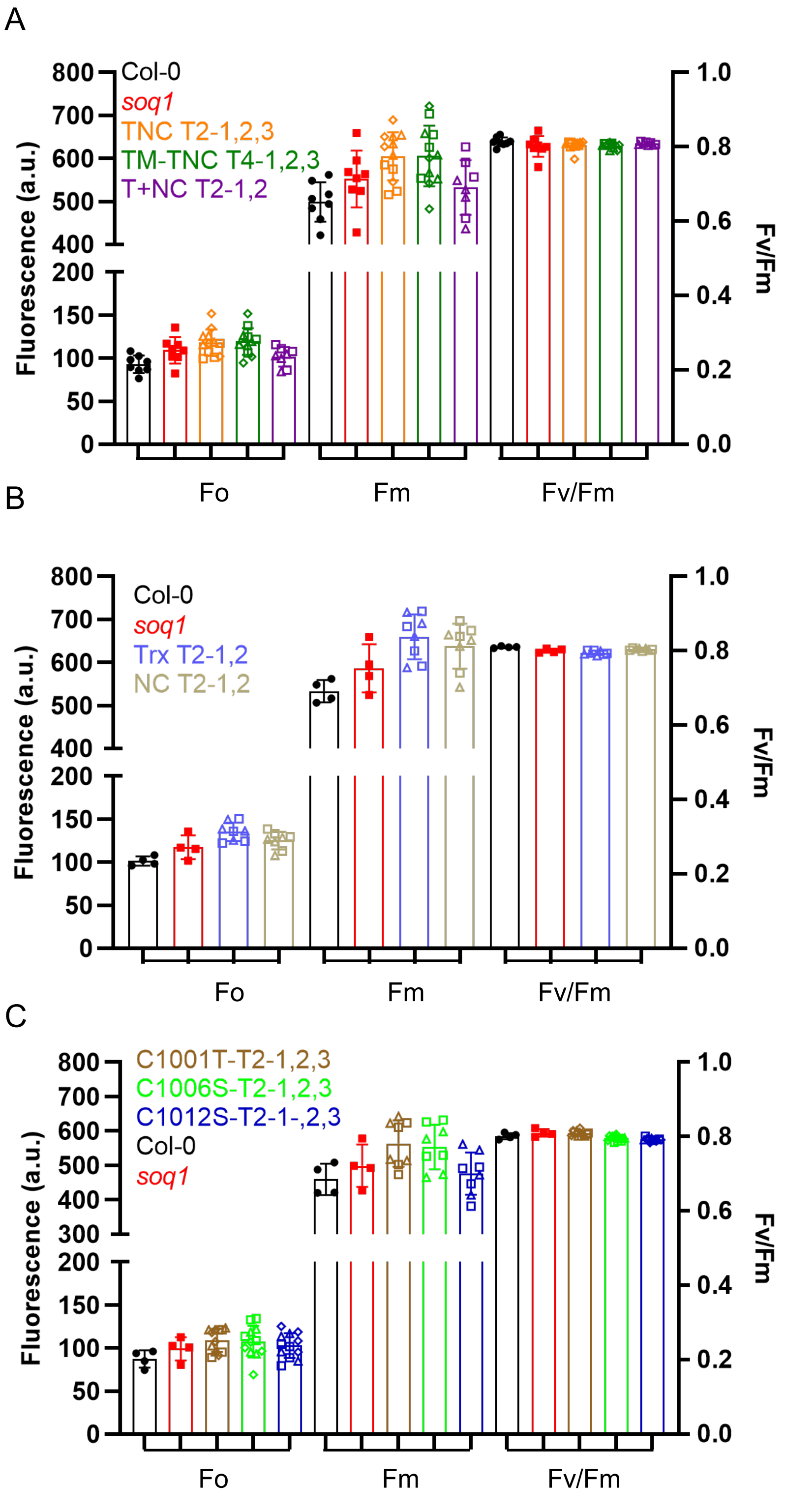

**Fig. S2 Photosynthetic parameters of the transformant lines containing SOQ1 variants.**
Photosynthetic parameters Fo, Fm and Fv/Fm (at time 0 of NPQ kinetics shown in Fig. 1C, 4C and S1C) of Col-0, *soq1*, TNC T2-1,2,3, TM-TNC T4-1,2,3, T+NC T2-1,2 (A), Trx T2-1,2 and NC T2-1,2 (B) and C1001T T2-1,2,3, C1006S T2-1,2,3 and C1012S T2-1,2,3 (C). Parameters are not statistically different from *soq1* by Tukey's multiple comparisons test (Table S1).

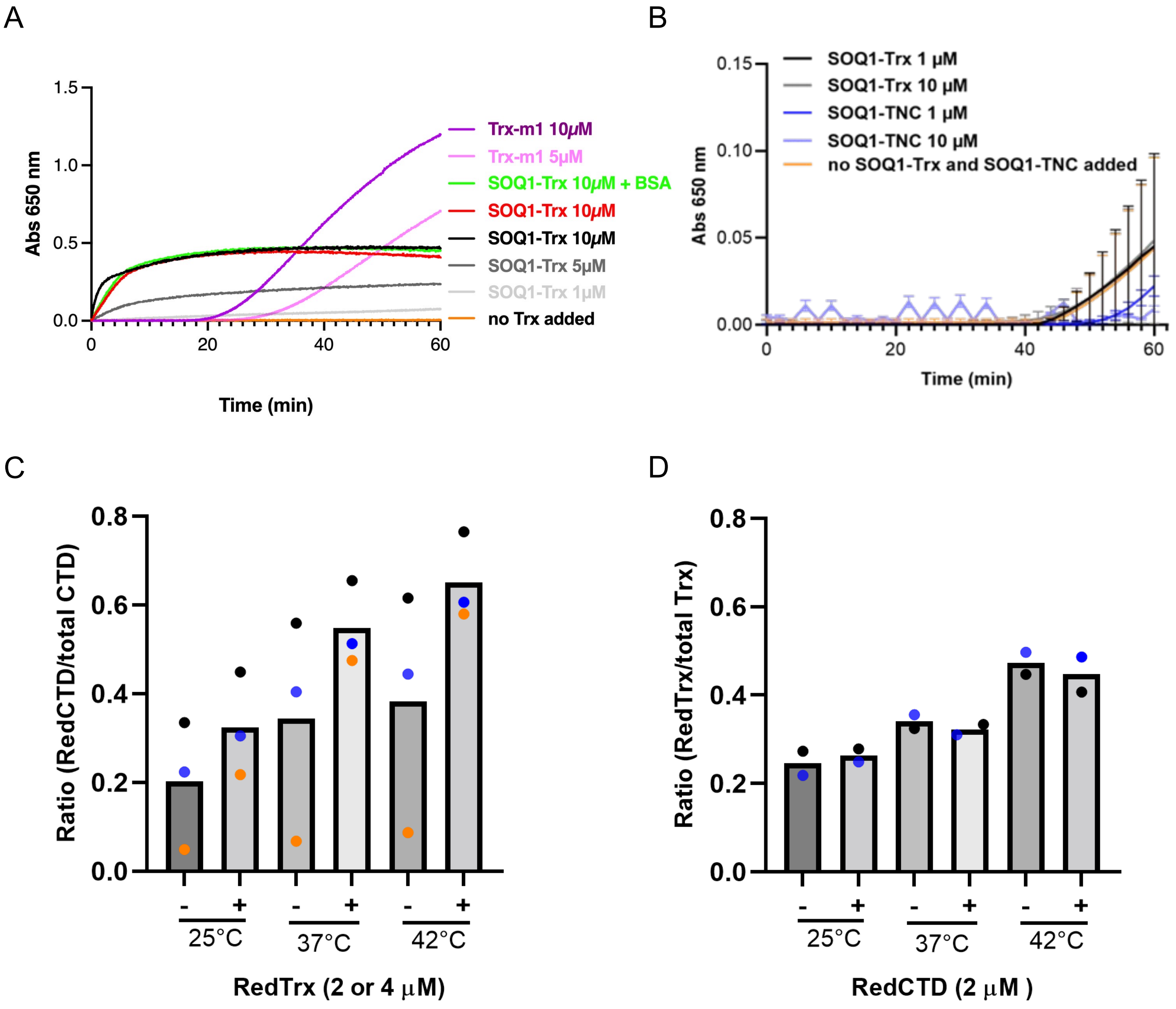

**Fig. S3 Electron transfer between SOQ1 Trx and its CTD (Figure 3).**
(A-B) SOQ1-Trx or SOQ1-TNC do not reduce insulin. Reduction of insulin is measured as a change in turbidity of the solution, due to the precipitation of free insulin B chain, at 650 nm. Potassium phosphate (KPi, pH7.0) was used for the reaction buffer. The reaction was initiated by adding 500 μM DTT. A, Trx-m1 was used as a positive control. SOQ1-Trx precipitates in this assay, independently of insulin presence: no insulin in samples plotted in red or green (here bovine serum albumin (BSA) was added to attempt preventing precipitation without success). B, We cloned a new version of SOQ1-Trx without the V5 epitope tag and TEV site and precipitation no longer occurred, neither did insulin in a time frame expected for disulfide reductase activity: see in the Trx-m1 control in (A), insulin starts to precipitate after approximately 20 min. C, The ratio of reduced SOQ1-CTD (RedCTD) to the total (RedCTD + OxCTD) is calculated from a CBB gel. It increases with addition (+) of RedTrx (2 μM, orange points or 4 μM, blue and black points) and increasing temperature. 2 μM partially oxidized SOQ1-CTD was used. D, The ratio of reduced SOQ1-Trx (RedTrx) to the total (RedTrx + OxTrx) calculated from a CBB gel does not change with addition (+) of RedCTD (2 μM) and increasing temperature. 1 μM (blue points) or 2 μM (black points) partially oxidized SOQ1-Trx was used. 0.2 mM DTT is present in all assays.

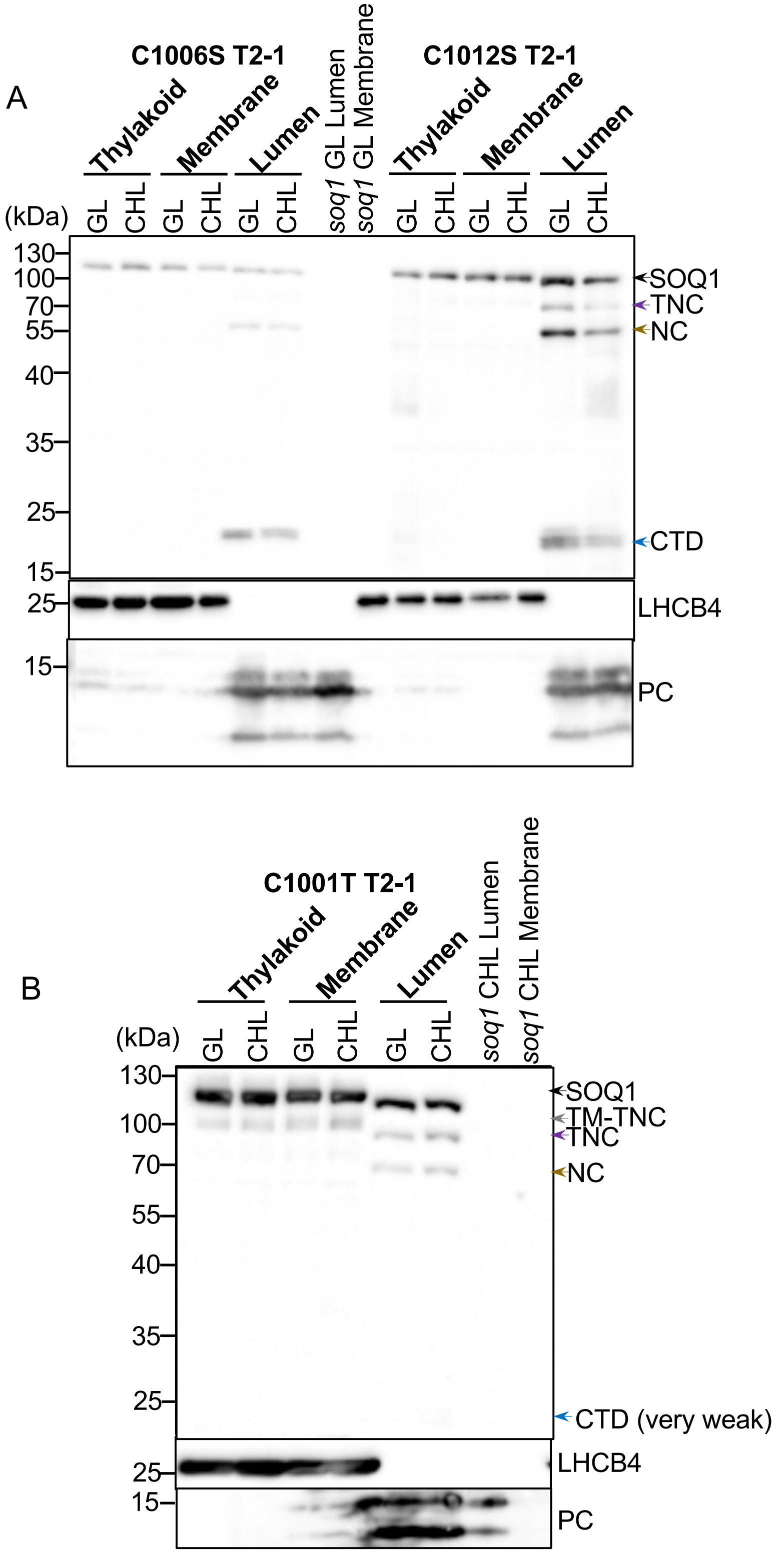

**Fig.S4 The localization of C1001T, C1006S and C1012S in chloroplast sub-fractions (Figure 4).**
A, Localization of C1006S and C1012S in chloroplast sub-fractions (Thylakoid, Membrane: thylakoid membrane after having separated the lumen fraction, and Lumen) in growth light (GL) or after 6h in cold high light (CHL). B, Localization of C1001T in chloroplast sub-fractions. In (A-B) Samples were loaded at the same amount of chlorophyll (1.5 µg) for thylakoid and membrane or lumenal proteins (5 µg). Proteins were separated by SDS-PAGE and analyzed by immunodetection with antibodies against SOQ1, PC, LHCB4. Representative immunoblots from two independent biological experiments are shown.

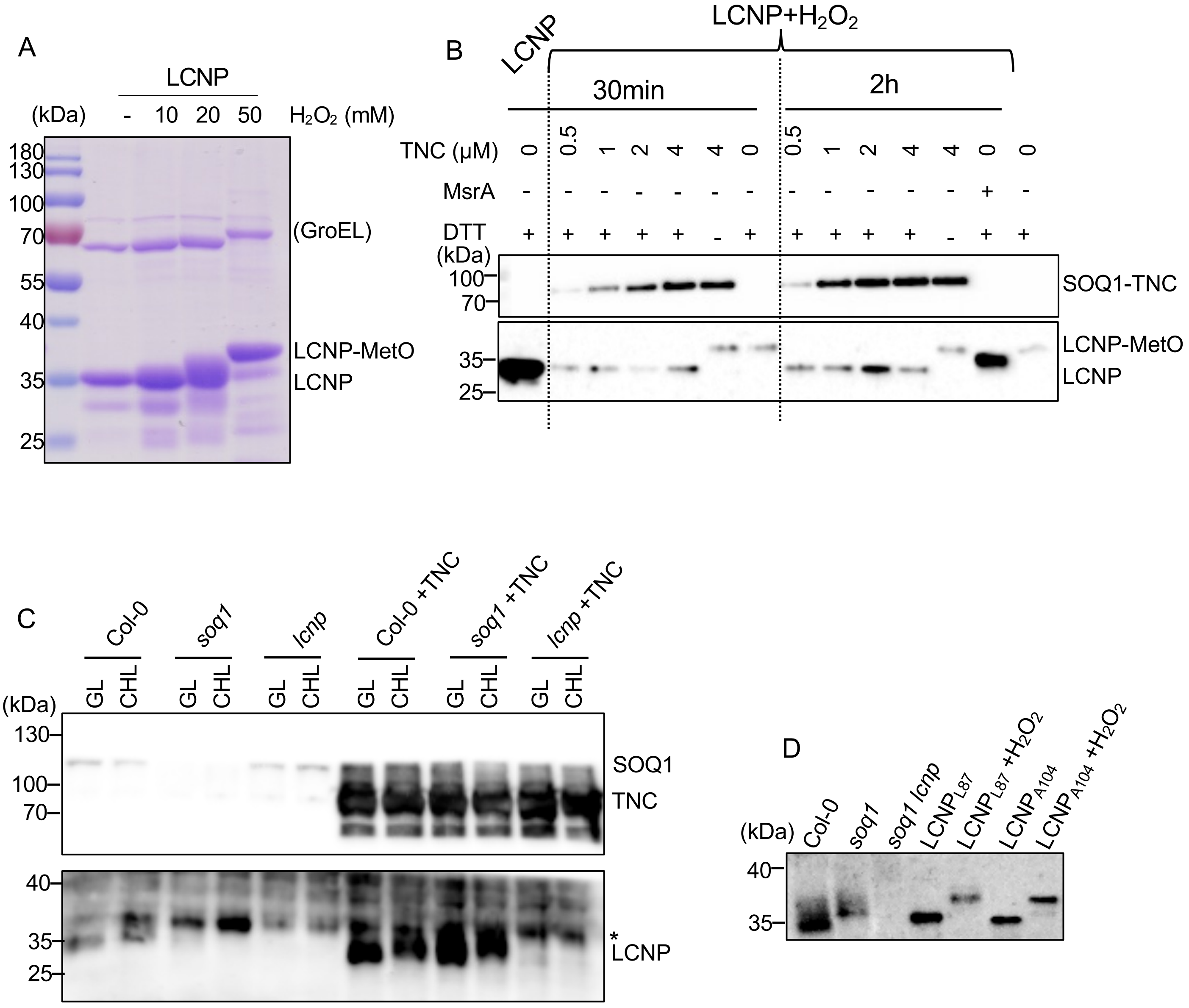

**Fig. S5 LCNP electrophoretic mobility is restored upon addition of methionine sulfoxide reductase or TNC.**
A, $H_2O_2$-treatment of recombinant LCNP. Coomassie stained gel of purified recombinant $LCNP_{A104}$ (LCNP) treated with increased concentrations of $H_2O_2$ (LCNP+$H_2O_2$). 50 mM was chosen. The upper band at ~70kDa is a contaminant protein from *E.coli* (GroEL determined by mass spectrometry). B, Addition of methionine sulfoxide reductase A (MsrA) or SOQ1-TNC to recombinant LCNP. 1 μM LCNP+$H_2O_2$ was incubated with different amount of TNC in the presence of 5 mM DTT at 25℃ for 30 min or with 3 μM MsrA at 37℃ for 2 h. The reaction was stopped by TCA precipitation. The samples were separated by SDS-PAGE and analyzed by immunodetection using anti-LCNP and anti-$SOQ1_{CTD}$ antibodies. Representative immunoblot from two independent biological experiments is shown. C, Incubation of whole cell proteins with TNC. 20 μg whole cell proteins from Col-0, *soq1* and *lcnp* grown under standard growth conditions and exposed to growth light (GL) or cold high light (CHL) conditions were incubated with or without 1 μM recombinant TNC for 30 min at room temperature under 5 mM DTT. Proteins were denatured by adding 1x SDS sample buffer and analyzed by immunodetection using anti-LCNP antibody. Star symbol (*) represents a nonspecific band, based on its presence in *lcnp,* detected by the anti-LCNP antibody likely originating from the TNC recombinant protein preparation. C, Thylakoid (2.5 μg chlorophyll) from Col-0, *soq1* or *soq1 lcnp* Arabidopsis mutants under cold and high light conditions and 1.5 ng recombinant $LCNP_{L87}$ or $LCNP_{A104}$, 3 ng recombinant $LCNP_{L87}$ +$H_2O_2$ or $LCNP_{A104}$ +$H_2O_2$ were separated by SDS-PAGE and analyzed by immunodetection using anti-LCNP. The recombinant proteins overall migrate slower than the native protein due to a Flag-tag in N-terminus (+1.01kD) there is also less crowding compared to thylakoid extracts. Native LCNP (expected size: 29.91kD), Flag-$LCNP_{L87}$ (expected size: 30.92kD) and Flag-$LCNP_{A104}$ (29.19kD). The L87 form of LCNP may correspond to the mature form of LCNP predicted by TargetP2.0.

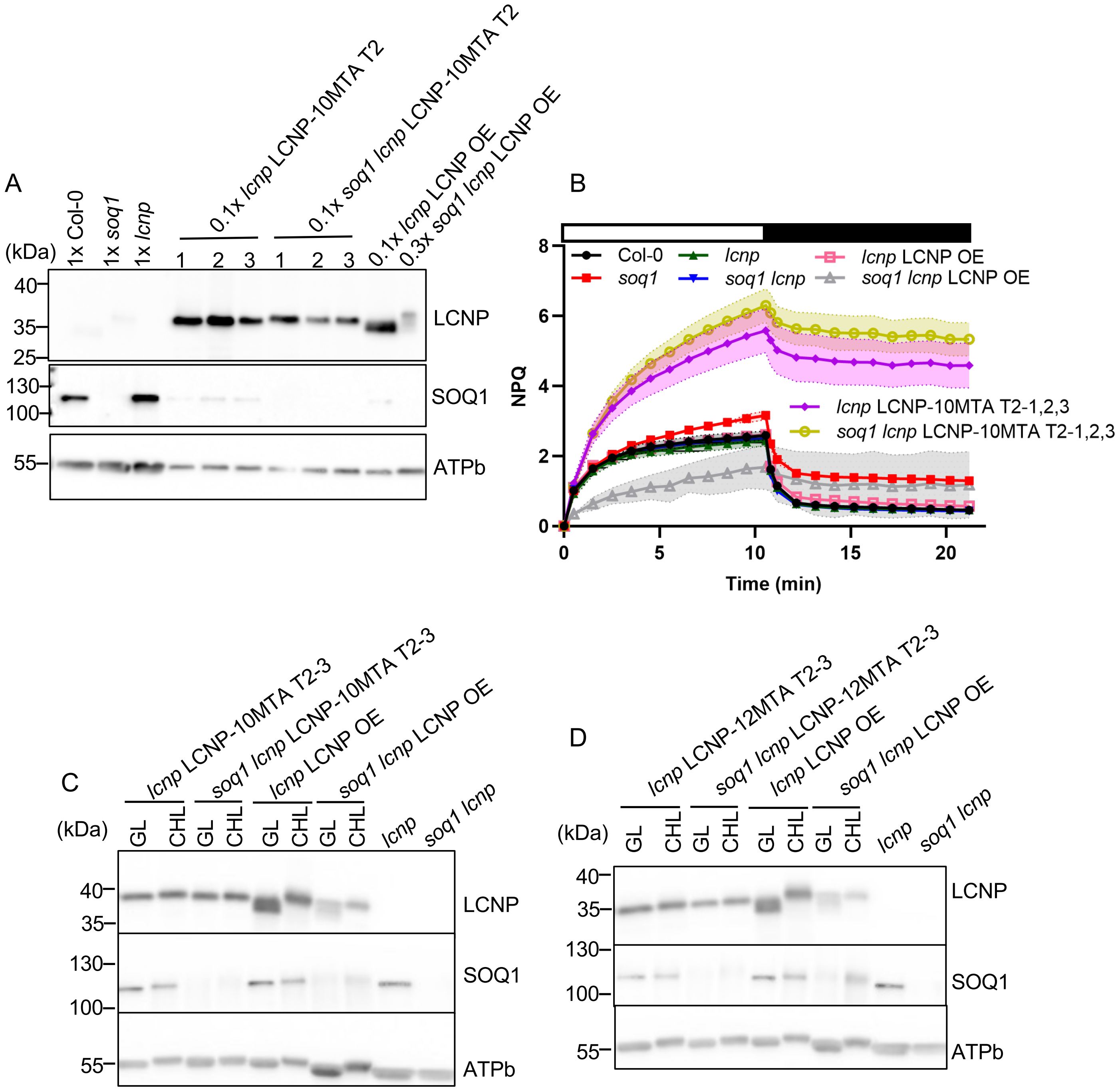

**Fig. S6 Slower electrophoretic mobility of LCNP is methionine-dependent**
A, Immunoblot analyses of three independent T2 *Arabidopsis* variant LCNP lines containing 10 methionine to alanine (MTA) mutations transformed in *lcnp* and *soq1 lcnp* mutant background. LCNP overexpressed in *lcnp* (*lcnp* LCNP OE) and *soq1 lcnp* (*soq1 lcnp* LCNP OE) mutant background were used as the LCNP mobility control. Samples were loaded at the same amount of total protein (1x corresponds to 15 µg), less proteins were loaded for the transformants as LCNP is overexpressed. ATPb was used as the loading control. B, NPQ kinetics of Col-0, *soq1,* three independent *lcnp* and *soq1 lcnp* T2 variant lines, *lcnp* LCNP OE and *soq1 lcnp* LCNP OE lines. Five-week-old plants grown at 150 µmol photons m$^{-2}$ s$^{-1}$ were dark-acclimated for 20 min and NPQ was induced at 1,200 µmol photons m$^{-2}$ s$^{-1}$ (white bar) and relaxed in the dark (black bar). Data represent means ± SD (n = 12; four individuals from three independent T2 lines). C, D, Immunoblot analyses of one independent T2 *Arabidopsis* variant LCNP line containing 10 or 12 methionine to alanine (MTA) mutations transformed in *lcnp* or *soq1 lcnp* mutant background under growth light (GL) or cold and high light (CHL) conditions. Here we can see that LCNP OE in *lcnp* background upon CHL migrates slower compared to GL as SOQ1 becomes limiting. Samples were loaded at the same amount of total protein (1.5 µg).

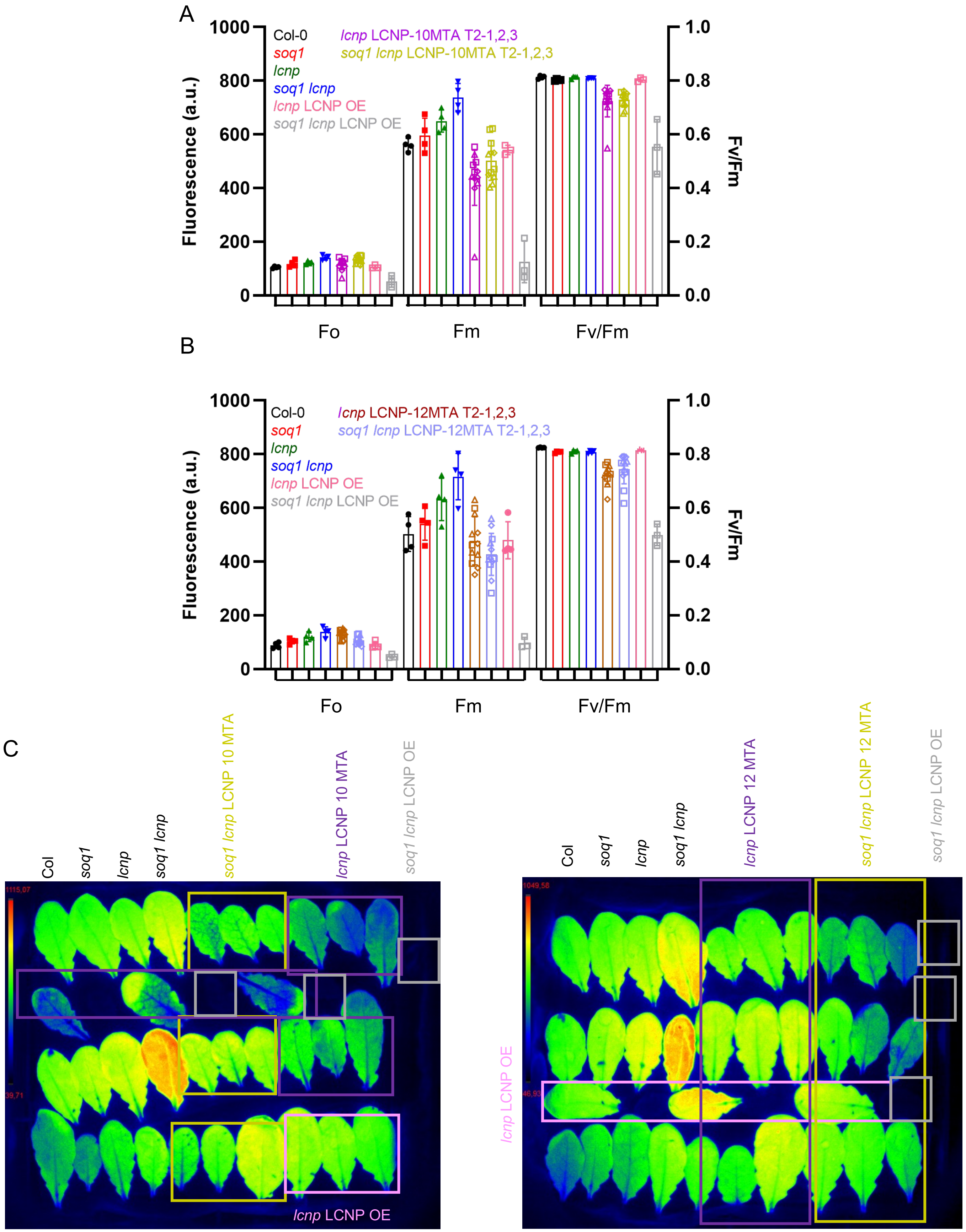

**Fig. S7 Photosynthetic parameters of the transformant LCNP lines containing 10 or 12 MTA**
Photosynthetic parameters Fo, Fm and Fv/Fm (at time 0 of NPQ kinetics shown in Fig. 6D,E) of Col-0, *soq1*, *lcnp* LCNP OE, *soq1 lcnp* LCNP OE, *lcnp* LCNP-10MTA T2-1,2,3, *soq1 lcnp* LCNP-10MTA T2-1,2,3 (A) and *lcnp* LCNP-12MTA T2-1,2,3, *soq1 lcnp* LCNP-12MTA T2-1,2,3 (B). Fm parameters are not statistically different in the MTA lines from Col-0 or *lcnp* LCNP OE by Tukey's multiple comparisons test (Table S1). C) The Fm of the *lcnp* and *soq1 lcnp* lines here are higher than usual and may be due to the leaves position under the imager for that experiment.

**Table S1 Statistical analyses of photosynthetic parameters in Fig. S2, S8**
Adjusted *p* values are displayed and were analyzed by Tukey's multiple comparisons test.
Parameters statistically different are denoted by * ($p < 0.05$).

| Parameter \ *soq1* vs | Col-0 | TNC T2-1,2,3 | TM-TNC T4-1,2-3 | T+NC T2-1,2,3 |
|---|---|---|---|---|
| $F_o$ | 0.1605 | 0.6272 | 0.5029 | 0.8178 |
| $F_m$ | 0.4202 | 0.3487 | 0.3277 | 0.9643 |
| $F_v/F_m$ | 0.4378 | 0.9770 | 0.9983 | 0.8247 |

| Parameter \ *soq1* vs | Col-0 | Trx T2-1,2 | NC T2-1,2 |
|---|---|---|---|
| $F_o$ | 0.1676 | 0.0519 | 0.6383 |
| $F_m$ | 0.4493 | 0.1134 | 0.3618 |
| $F_v/F_m$ | 0.0238* | 0.2192 | 0.5398 |

| Parameter \ *soq1* vs | Col-0 | C1001T T2-1,2,3 | C1006S T2-1,2-3 | C1012S T2-1,2,3 |
|---|---|---|---|---|
| $F_o$ | 0.8019 | 0.7886 | 0.8673 | 0.9932 |
| $F_m$ | 0.8971 | 0.4562 | 0.6141 | 0.9739 |
| $F_v/F_m$ | 0.4057 | 0.3722 | 0.5180 | 0.1494 |

| | Col-0 *vs* *lcnp* LCNP-10MTA | Col *vs* *soq1 lcnp* LCNP- 10MTA | *lcnp* LCNP OE vs *lcnp* LCNP-10MTA | *lcnp* LCNP OE vs *soq1 lcnp* LCNP-10MTA |
|---|---|---|---|---|
| $F_o$ | 0.8859 | 0.0234* | 0.9718 | 0.0888 |
| $F_m$ | 0.1246 | 0.8819 | 0.4201 | 0.9919 |
| $F_v/F_m$ | 0.0162* | 0.0266* | 0.0974 | 0.1398 |

| | Col-0 *vs* *lcnp* LCNP-12MTA | *Col vs* *soq1 lcnp* LCNP-12MTA | *lcnp* LCNP OE vs *lcnp* LCNP-12MTA | *lcnp* LCNP OE vs *soq1 lcnp* LCNP-12MTA |
|---|---|---|---|---|
| $F_o$ | 0.0009 | 0.4439 | 0.0012* | 0.5027 |
| $F_m$ | 0.9991 | 0.6981 | > 0.9999 | 0.9360 |
| $F_v/F_m$ | 0.0006* | 0.0096 | 0.0026* | 0.0343* |

**Table S2 Statistical analyses of NPQ levels in Fig. S1D, E**
Adjusted *p* values are displayed and were analyzed by Tukey's multiple comparisons test. Parameters statistically different are denoted by * ($p < 0.05$).

| Col-0 vs | *soq1* | TNC T2-1,2,3 | TM-TNC T4-1,2-3 | T+NC T2-1,2,3 |
|---|---|---|---|---|
| NPQ | < 0.0001* | 0.1046 | 0.9873 | < 0.0001* |

| *soq1* vs | Col-0 | Trx T2-1,2 | NC T2-1,2 |
|---|---|---|---|
| NPQ | < 0.0001* | 0.4499 | 0.4962 |

| *soq1* vs | Col-0 | C1001T T2-1,2,3 | C1006S T2-1,2-3 | C1012S T2-1,2,3 |
|---|---|---|---|---|
| NPQ | < 0.0001* | < 0.0001* | 0.9968 | 0.8812 |

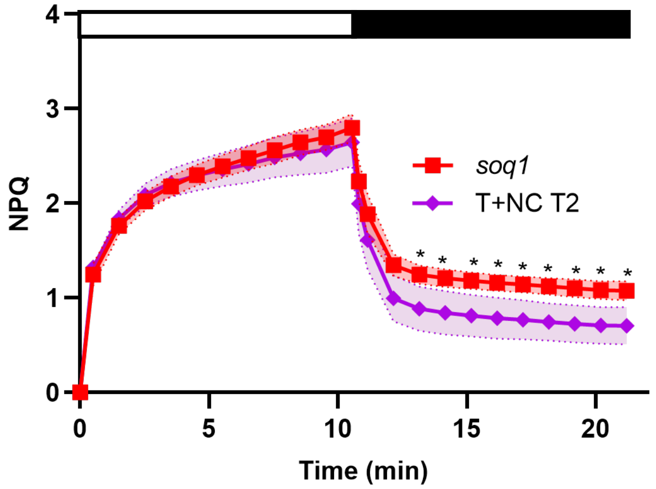

**Statistical analyses of NPQ kinetics of *soq1* and T+NC.**
Significant difference were analyzed by multiple unpaired t test. NPQ kinetics statistically different are denoted by * ($p < 0.05$).

### Table S3 Primers list

| Constructs | Primer name | Primer sequence | Plasmid | Comments |
|---|---|---|---|---|
| **For recombinant proteins** | | | | |
| | Forward primer | CAC CAA GCA AAC AGC TAC AAC TG | | |
| **Trx-NHL-CTD** | Reverse primer | TCA GCG AGT ACC TTG AAG CTG | pET151/D-TOPO | |
| | Forward primer | CAC CAA GCA AAC AGC TAC AAC TG | | |
| **Trx** | Reverse primer | TCA CAG AGC TGC CGC CAC CAC G | pET151/D-TOPO | |
| | Forward primer | CAC CCC GTT GAA ATT TCC GGG A | | |
| **NHL-CTD** | Reverse primer | TCA GCG AGT ACC TTG AAG CTG | pET151/D-TOPO | |
| | Forward primer | CACC GAGTTAAAAGGTGTTCAAC | | |
| **CTD** | Reverse primer | TCA GCG AGT ACC TTG AAG CTG | pET151/D-TOPO | |
| | Forward primer | CTG GAT CTG AGC AGC GGC TAC C | | |
| **LCNP** | Reverse primer | TTC AGG GTC TCG AAC ACG CTG G | pET24d fused SUMO tag | |
| **For SOQ1 transformants** | | | | |
| | Forward primer | tgagaactcgagATGGCAGCCTCTCTCCAA | | |
| **PsbO cTP/lTP** | Reverse primer | cagataactcgagCTTTGGAGCTCCCTCCG | n/a | Cloned into pEarleyGate100 via XhoI restriction site |
| | Forward primer | AACTGGAAGGCAATGCAATATG | | |
| **ΔHAD/TM** | Reverse primer | CGATACTTTCCCCCAATCATC | n/a | Cloned from Full-length SOQ1-C-FLAG |
| | Forward primer | **CAGTTTGAAAAA**GGATGACGACGATAAGTAGAAGG | | |
| **Trx (SOQ1 cTP)** | Reverse primer | **CGGATGGCTCCA**CGGAAATTTCAACGGAGAC | n/a | Cloned from ΔHAD/TM. Sequence in bold adds Strep-tag |
| | Forward primer | AACTGGAAGGCAATGCAATATG | | |
| **Trx-NHL-CTD (PsbO cTP)** | Reverse primer | TTTCAAAGCCATGGTGAAGG | pEarleyGate100 + PsbO cTP/lTP | Cloned from ΔHAD/TM |
| | Forward primer | AACTGGAAGGCAATGCAATATG | | |
| **Trx (PsbO cTP)** | Reverse primer | TTTCAAAGCCATGGTGAAGG | pEarleyGate100 + PsbO cTP/lTP | Cloned from Trx (SOQ1 cTP) |
| | Forward primer | GTATTAGACAGTACTCCGCTTCCAAC | | |
| **NHL-CTD (PsbO cTP)** | Reverse primer | TTTCAAAGCCATGGTGAAGG | pEarleyGate100 + PsbO cTP/lTP | Cloned from Full-length SOQ1-C-FLAG |
| | Forward primer | CAAAGAAGACGAGGTTTGCTTG | | |
| **C1001T** | Reverse primer | CAATAGTACACCTTcgtACTGATTTTC | pEarleyGate100 | Cloned from Full-length SOQ1-C-FLAG |
| | Forward primer | CAAAGAAGACGAGGTTTGCTTG | | |
| **C1006S** | Reverse primer | ctATAGTACACCTTGCAACTGATTTTC | pEarleyGate100 | Cloned from Full-length SOQ1-C-FLAG |
| | Forward primer | CAAAGAAGACGAGGTTagcTTG | | |
| **C1012S** | Reverse primer | CAATAGTACACCTTGCAACTGATTTTC | pEarleyGate100 | Cloned from Full-length SOQ1-C-FLAG |
| **For LCNP transformants** | | | | |
| Met121-122-123 to Ala | Multi site primer | tcttccatctgatggtgactcggaatcagcggcggcgatgatgatgagaggcatgactgctaag | pCambia1300 with UBQ10 promoter | |
| Met124-125-126 to Ala | Multi site primer | ctgatggtgactcggaatcagcggcggcggcggcgagaggcatgactgctaagaactttgacc | pCambia1300 with UBQ10 promoter | |
| Met309-310 to Ala | Multi site primer | tgatgctgagctagcagccgcggcgtccatgccaggtatggag | pCambia1300 with UBQ10 promoter | |
| Met312-315 to Ala | Multi site primer | ctagcagccgcggcgtccgcgccaggtgcggagcaaacactgaccaa | pCambia1300 with UBQ10 promoter | |
| M172A | | CAG GGA GTA TAC ACG TTT GAT GCG AAG GAA TCA GCC ATT AGA GT | pCambia1300 with UBQ10 promoter | |
| M219A | | GAG ACT GAC TTA GAA AAG CAA GAG GCG ATT AAA GAG AAG TGT TTC CTA CG | pCambia1300 with UBQ10 promoter | |
