## Supplementary material for "Suppressor of quenching 1 functions as a methionine sulfoxide reductase in the chloroplast lumen for regulation of photoprotective qH in Arabidopsis": Source data

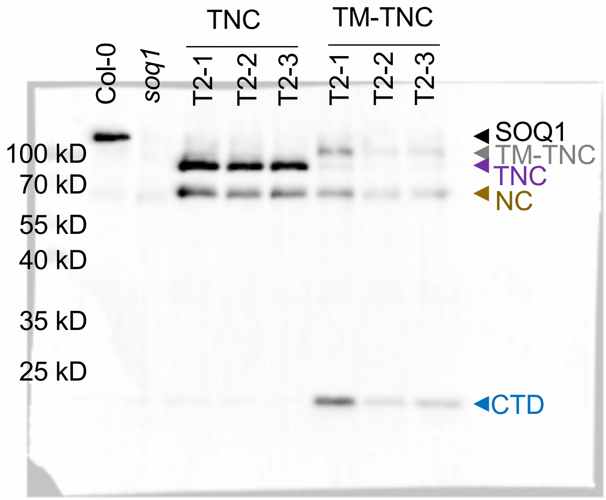
TNC
TM-TNC
Col-0
soq1
T2-1
T2-2
T2-3
T2-1
T2-2
T2-3
100 kD
70 kD
55 kD
40 kD
35 kD
25 kD
SOQ1
TM-TNC
TNC
NC
CTD

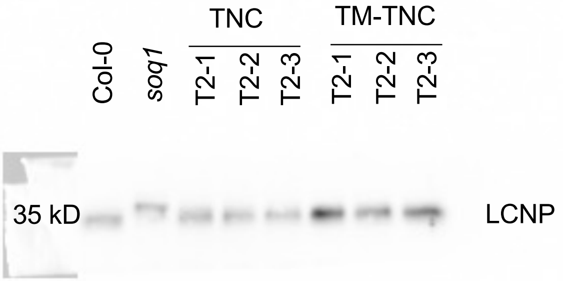
TNC
TM-TNC
Col-0
soq1
T2-1
T2-2
T2-3
T2-1
T2-2
T2-3
35 kD
LCNP

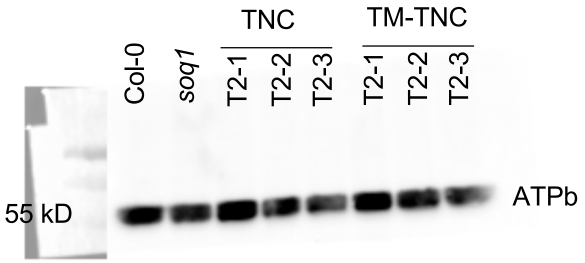
TNC
TM-TNC
Col-0
soq1
T2-1
T2-2
T2-3
T2-1
T2-2
T2-3
55 kD
ATPb

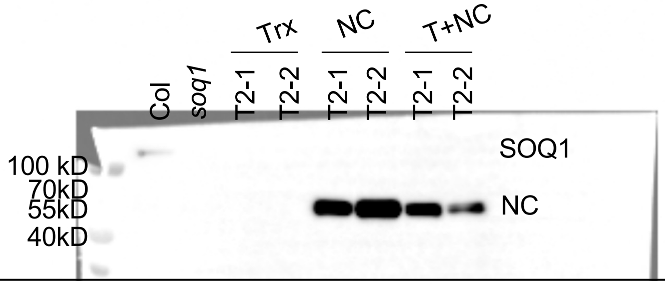


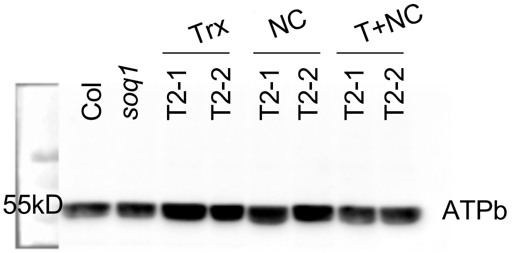


**Figure 1C Upper and 2nd Panel Source Data. Immunoblot analysis showed the acccumulation of Trx, NC and T+NC in two individual lines.** Uncropped version. PVDF Membrane was cut and probed with anti-SOQ1 and anti-ATPb antibodies. (Replicate number 1)

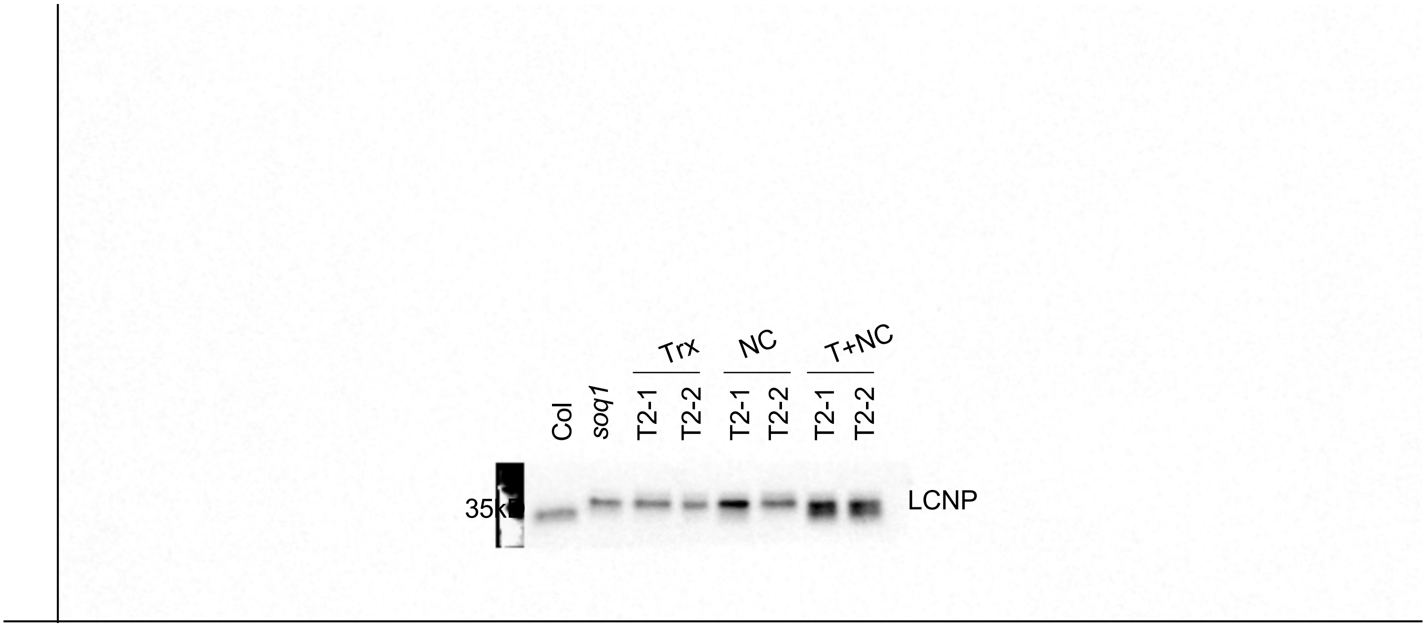


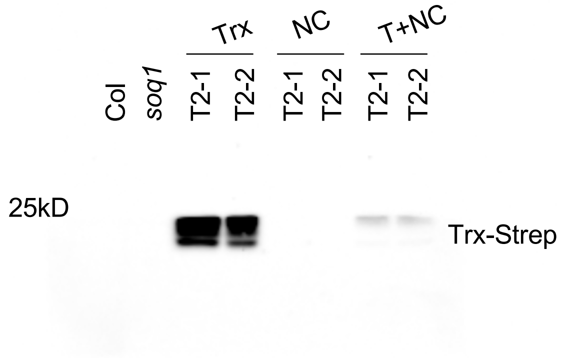


**Figure 1C 3rd and Bottom Panel Source Data. Immunoblot analysis showed the acccumulation of Trx, NC and T+NC in two individual lines.** Uncropped version. PVDF Membrane was cut and probed with anti-LCNP and anti-Strep antibodies. (Replicate number 1)

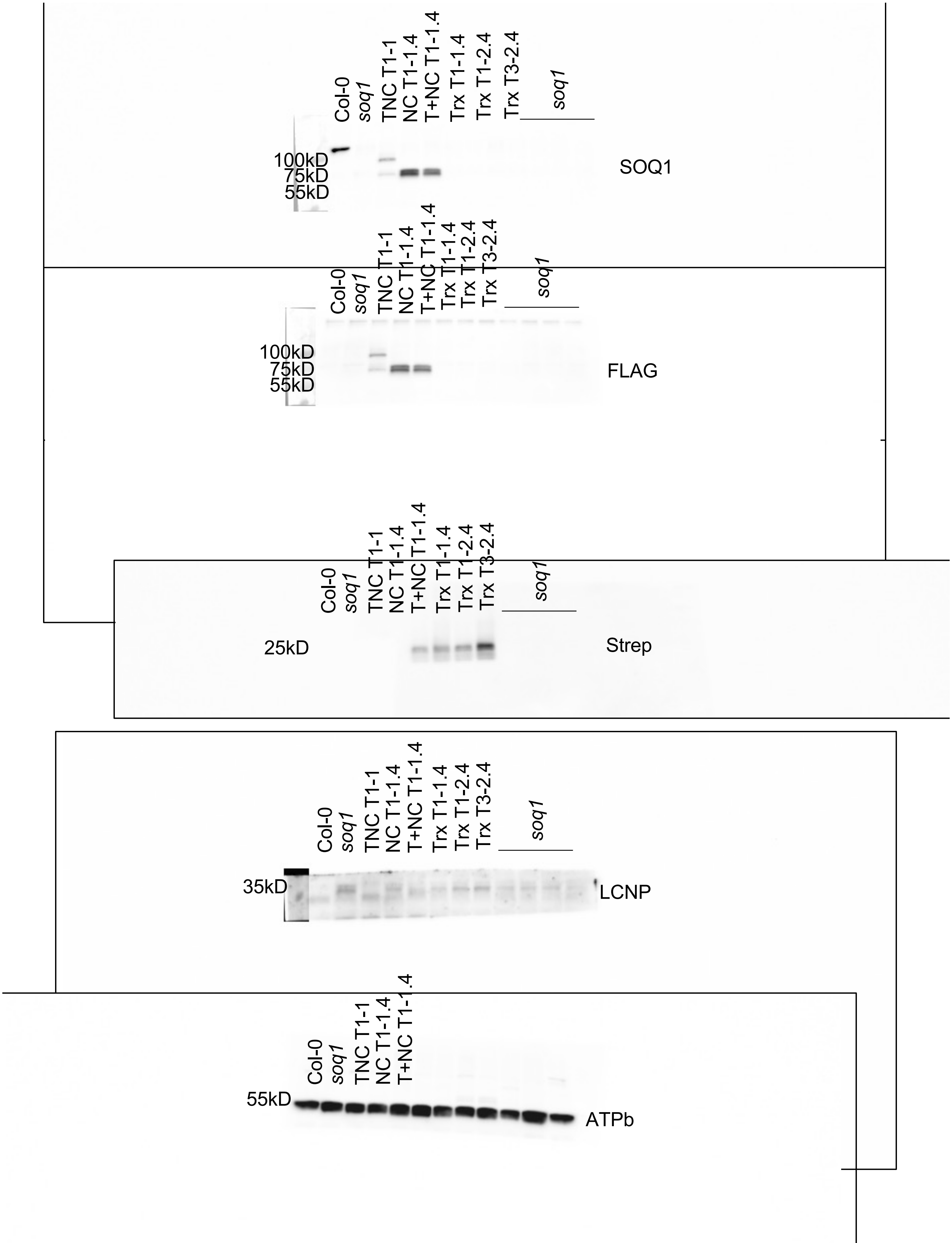


**Supporting Figure 1B and 1C Source Data. Immunoblot analysis showed the acccumulation of TNC, Trx, NC and T+NC in T1 lines.** Uncropped version. PVDF Membrane was cut and probed with anti-SOQ1, anti-Flag and anti-LCNP antibodies. (Replicate number 2)

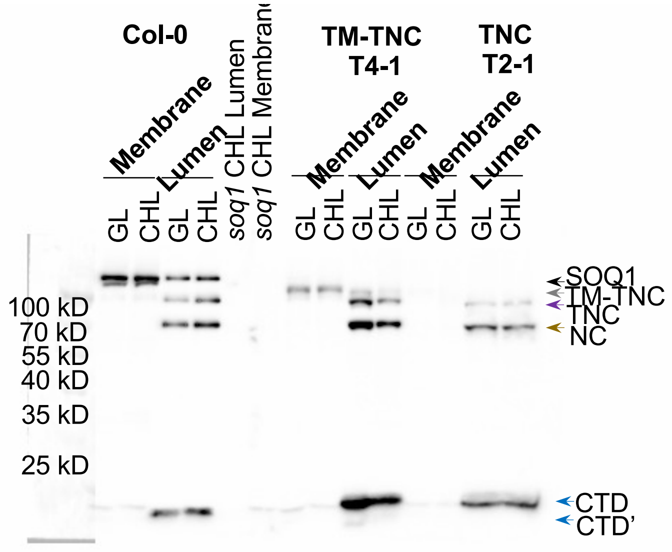


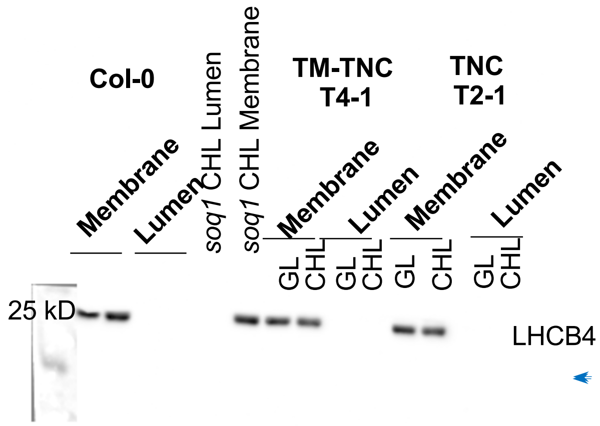


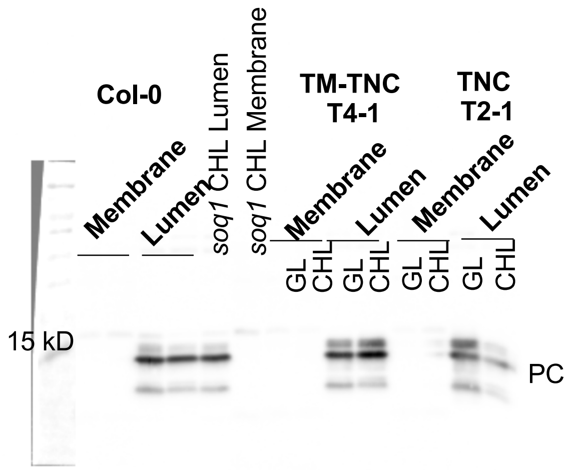


**Figure S1A Source Data. Immunoblot analysis showed the localization of TNC and TM-TNC.** Uncropped version. PVDF Membrane was probed with anti-SOQ1$_{CTD}$ and anti-LHCB4 and anti-PC antibodies. (Replicate number 1)

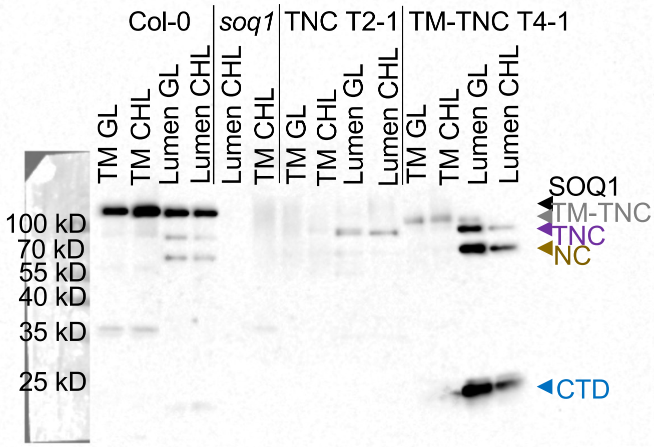


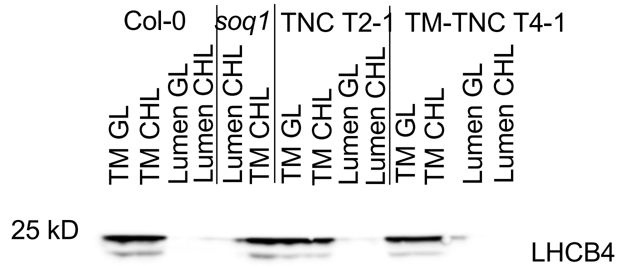


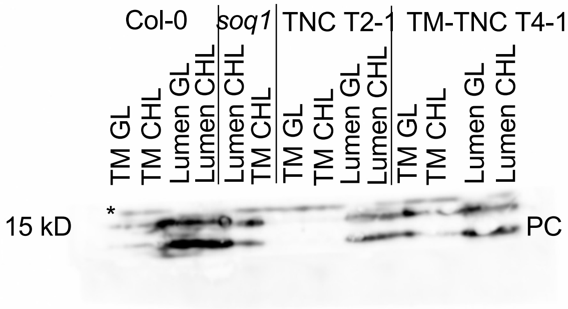


**Supporting Figure S1A Source Data. Immunoblot analysis showed the localization of TNC and TM-TNC in three individual lines.** Uncropped version. PVDF Membrane was probed with anti-SOQ1$_{CTD}$, anti-LHCB4 and anti-PC antibodies. (Replicate number 2)

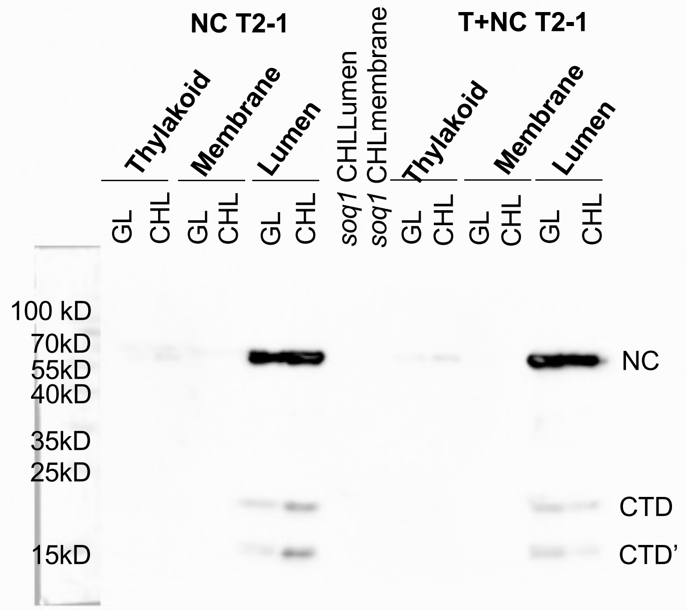


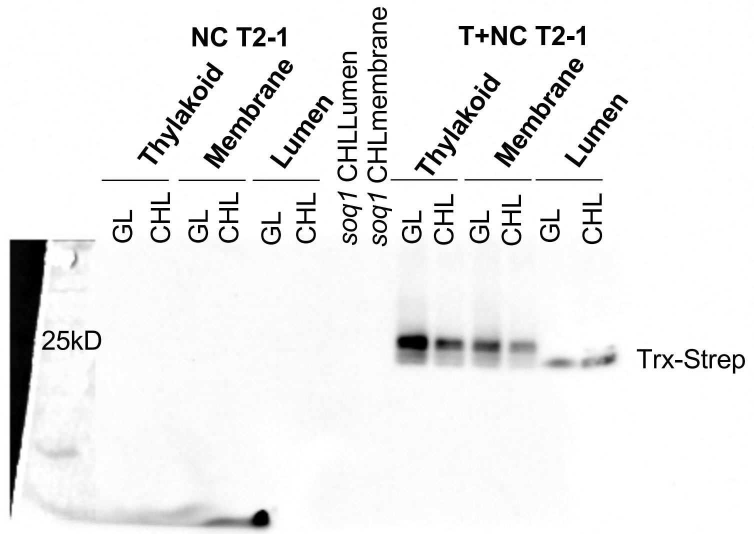


**Figure S1B NC and T+NC localization Source Data. Immunoblot analysis showed the localization of NC and T+NC.** Uncropped version. PVDF Membrane was probed with anti-$SOQ1_{CTD}$ and anti-Strep antibodies. (Replicate number 1)

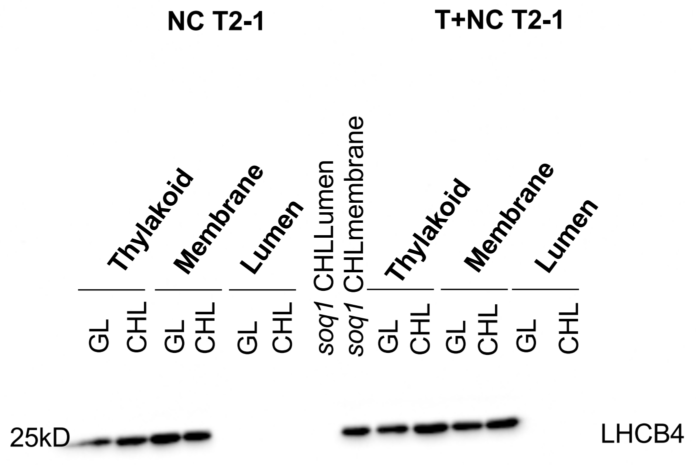


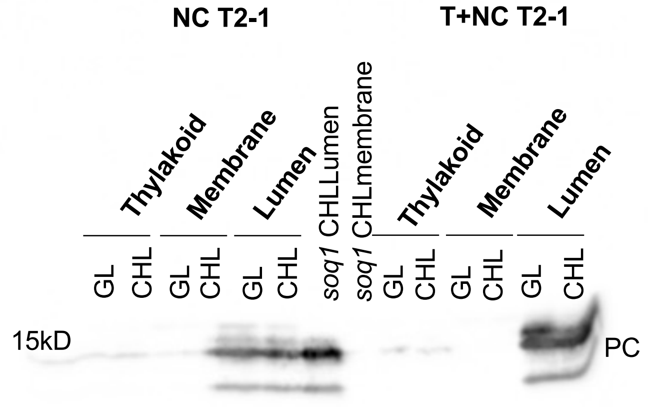


**Figure S1B NC and T+NC localization Source Data. Immunoblot analysis showed the localization of NC and T+NC.** Uncropped version. PVDF Membrane was probed anti-LHCB4 and anti-PC antibodies. (Replicate number 1)

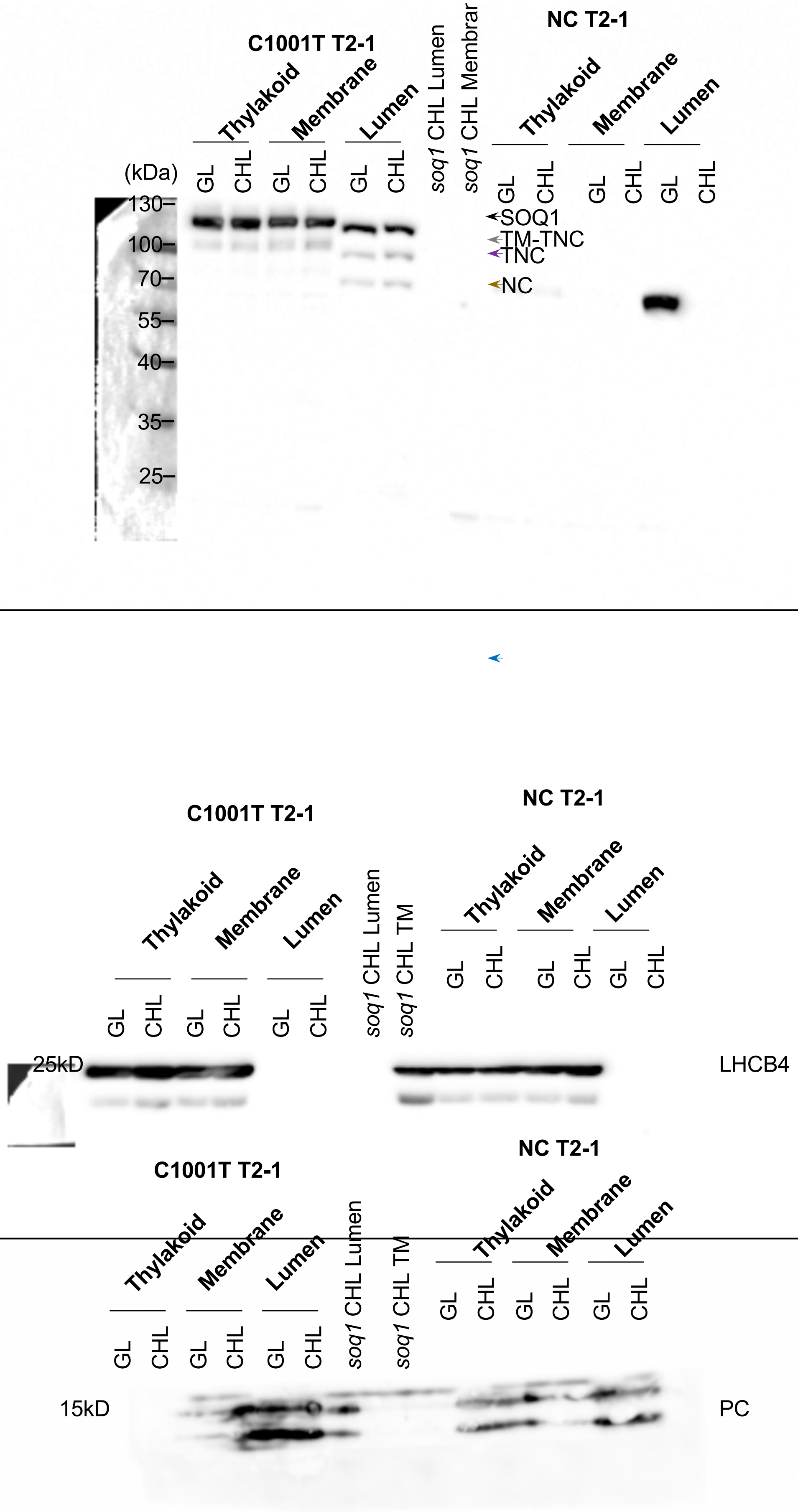


**Supporting Figure S1B NC localization Source Data. Immunoblot analysis showed the localization of NC.** Uncropped version. PVDF Membrane was probed $SOQ1_{CTD,}$ anti-LHCB4 and anti-PC antibodies. (Replicate number 2)

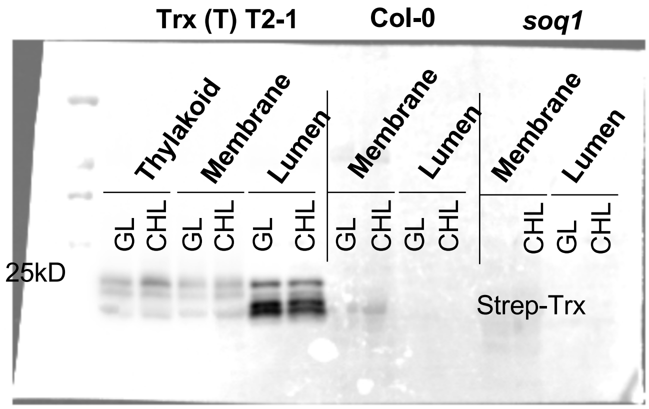


**Figure S1B Trx (T) localization Source Data. Immunoblot analysis showed the localization of Trx.** Uncropped version. PVDF Membrane was probed with anti-Strep antibodies.

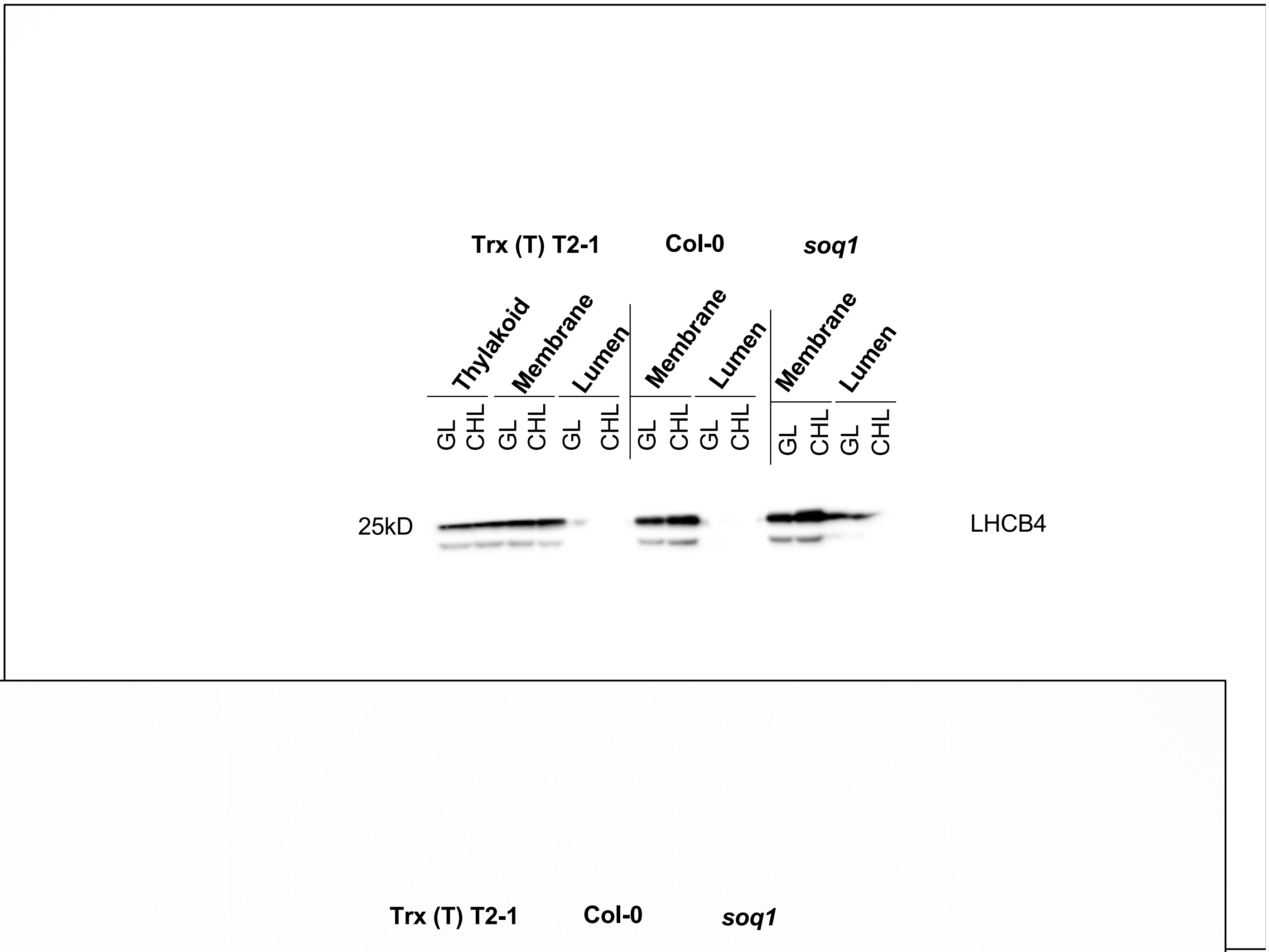


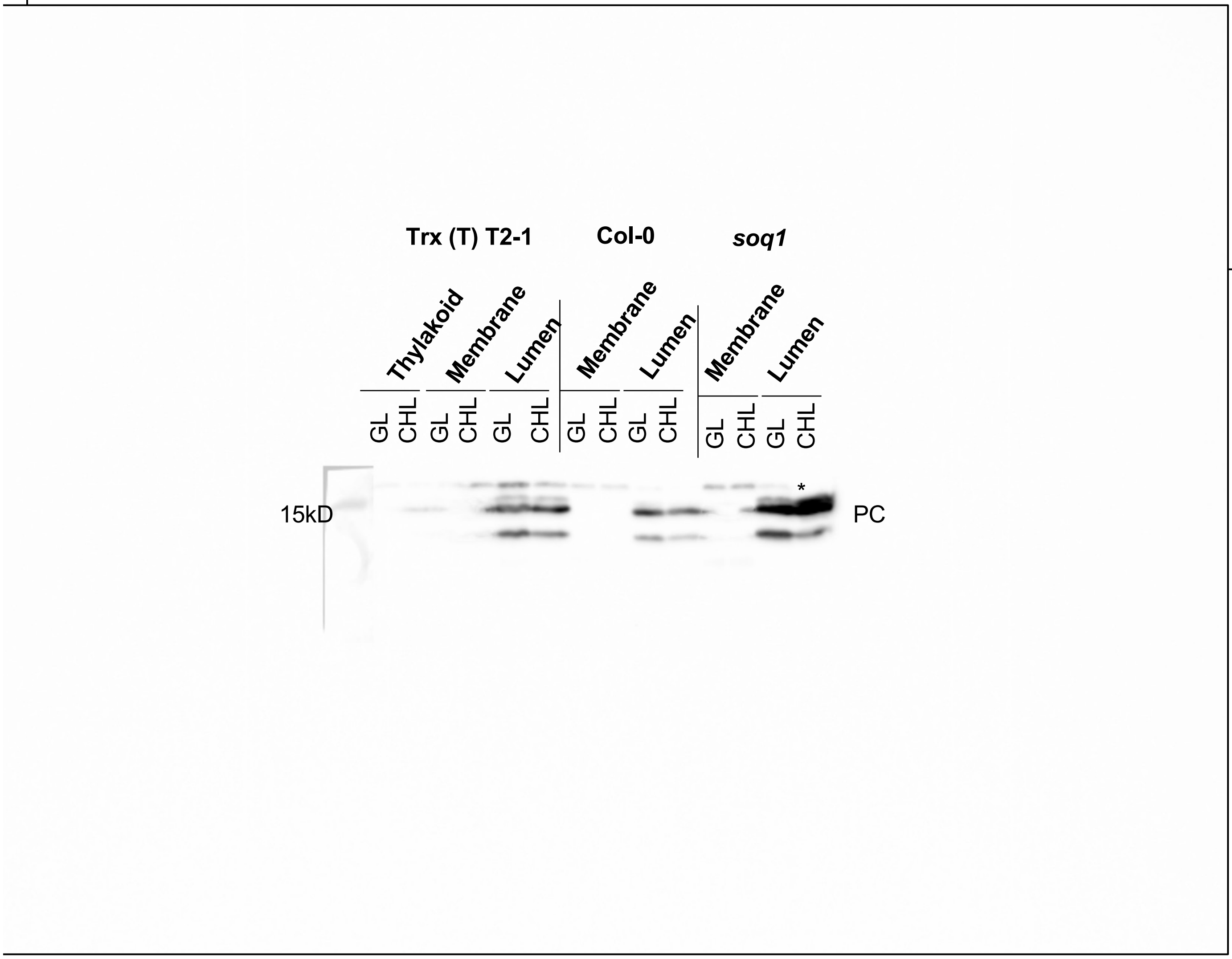


**Figure S1B Trx (T) localization Source Data. Immunoblot analysis showed the localization of Trx (T) in T2-1 line.** Uncropped version. PVDF Membrane was probed with anti-Strep, anti-LHCB4 and anti-PC antibodies. Star symbol (*) represents nonspecific band by anti-PC.

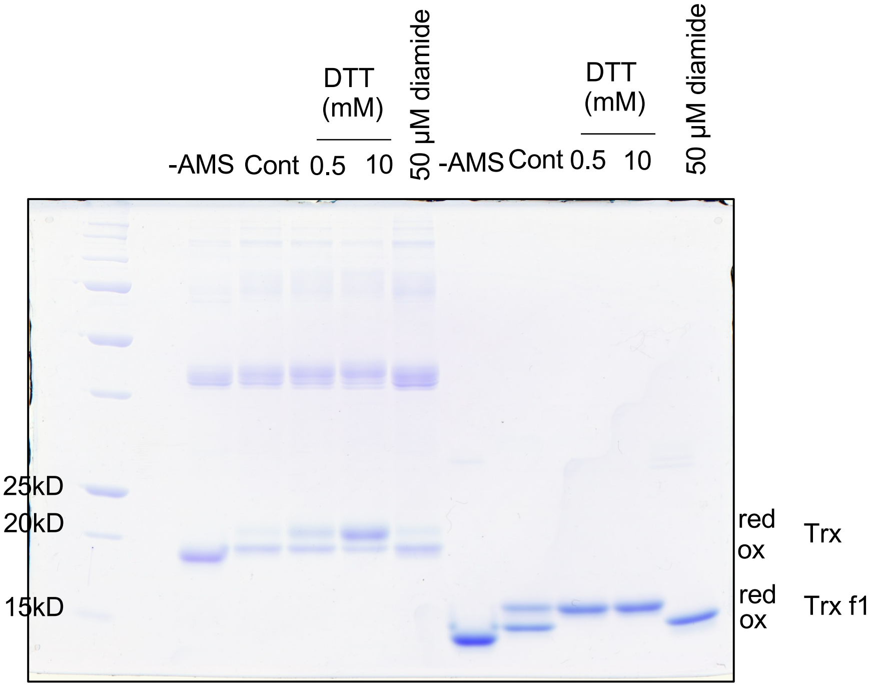


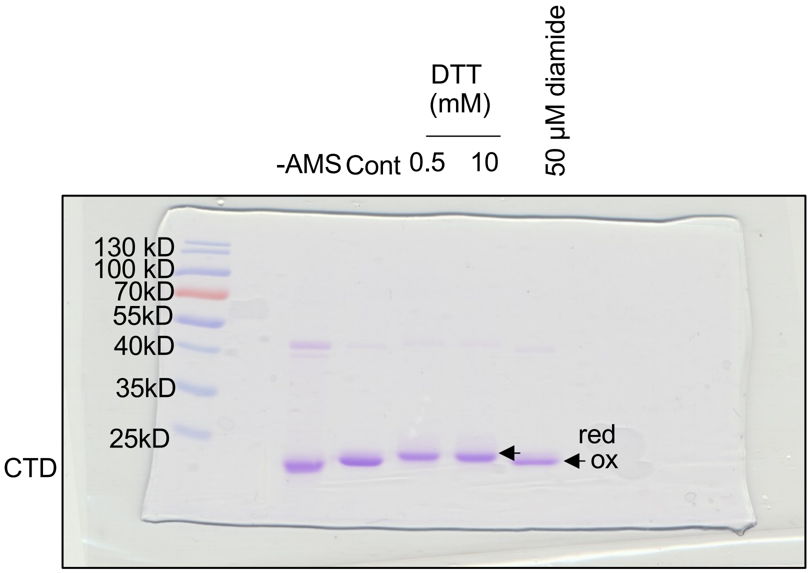


**Figure 2A Source Data. The redox state of Trx and CTD in the different redox conditions. Three Cys in the Trx and CTD, respectivity. Trx f1 was used as the control.**Uncropped version. (Replicate number 1)

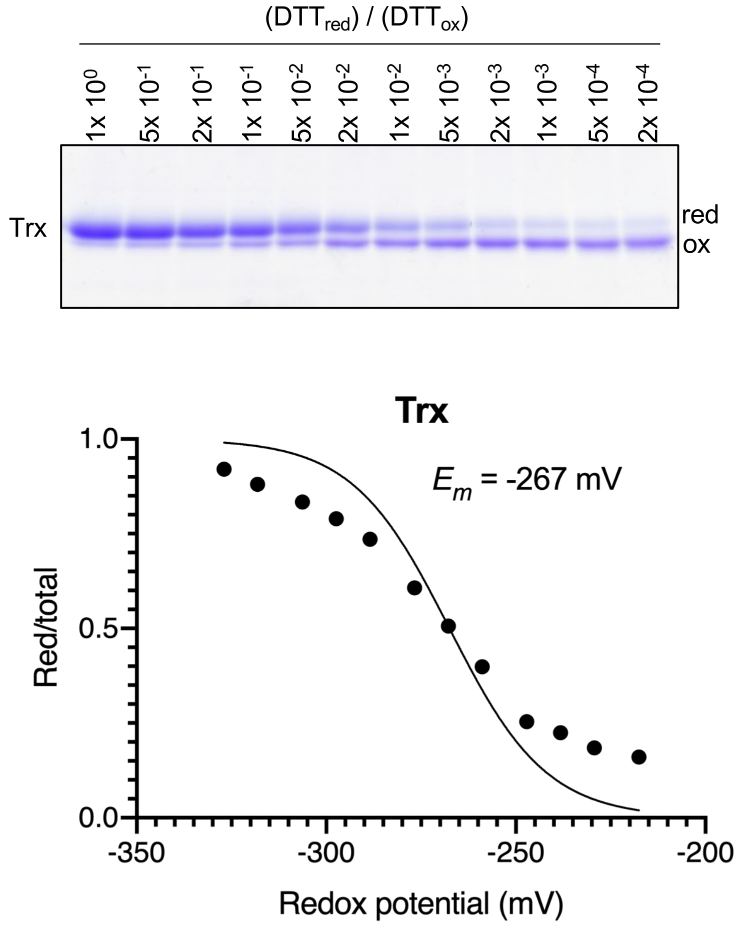


**Figure 2B-C Source Data. The SDS-polyacrylamide gel images of redox titration of Trx. Three Cys in the Trx.** (Replicate number 1)

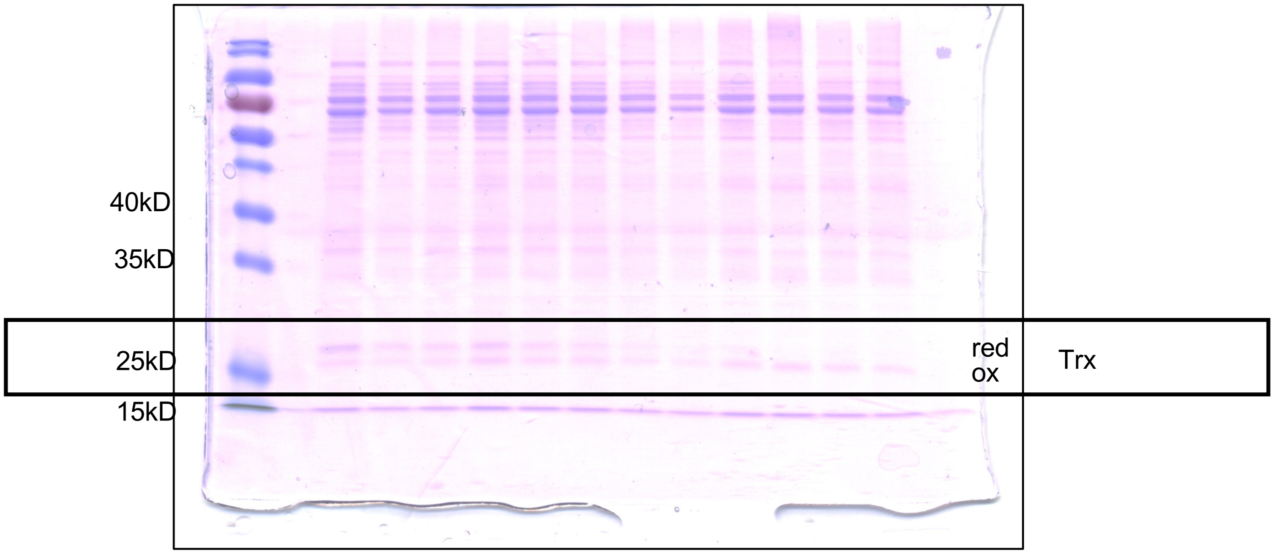


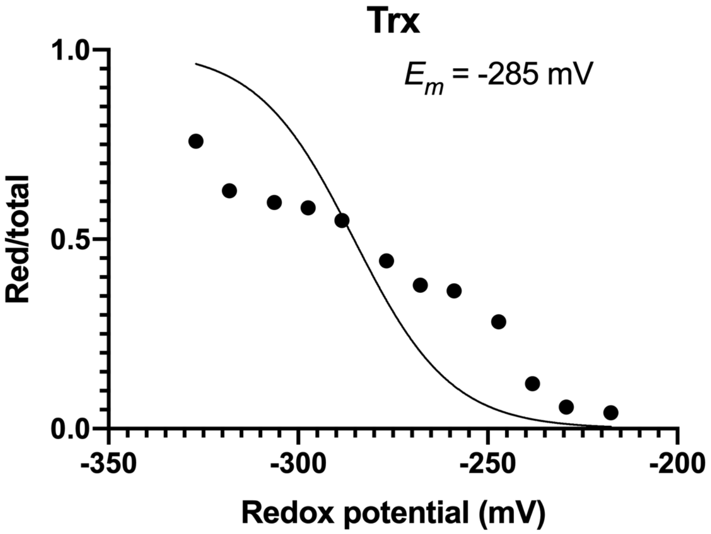


**Supporting Figure 1B-C Source Data. The SDS-polyacrylamide gel images of redox titration of Trx. Three Cys in the Trx.** Uncropped version (Replicate number 2)

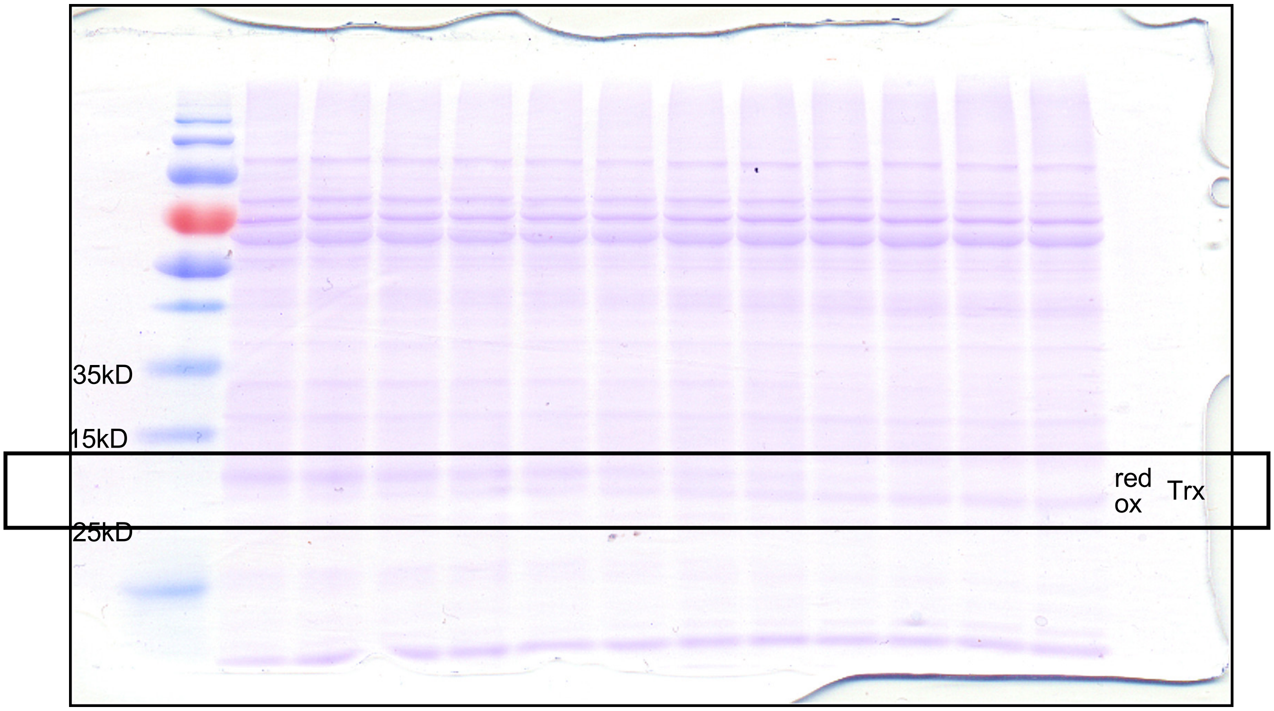


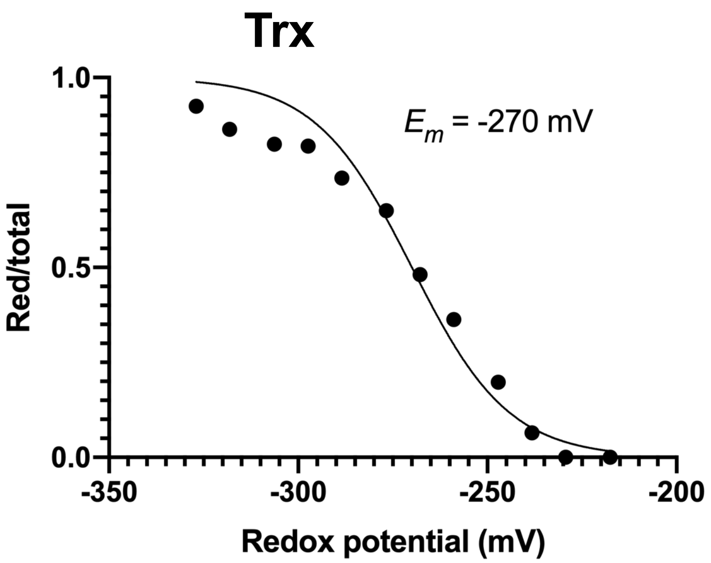


**Supporting Figure 1B-C Source Data. The SDS-polyacrylamide gel images of redox titration of Trx. Three Cys in the Trx.** Uncropped version (Replicate number 3)

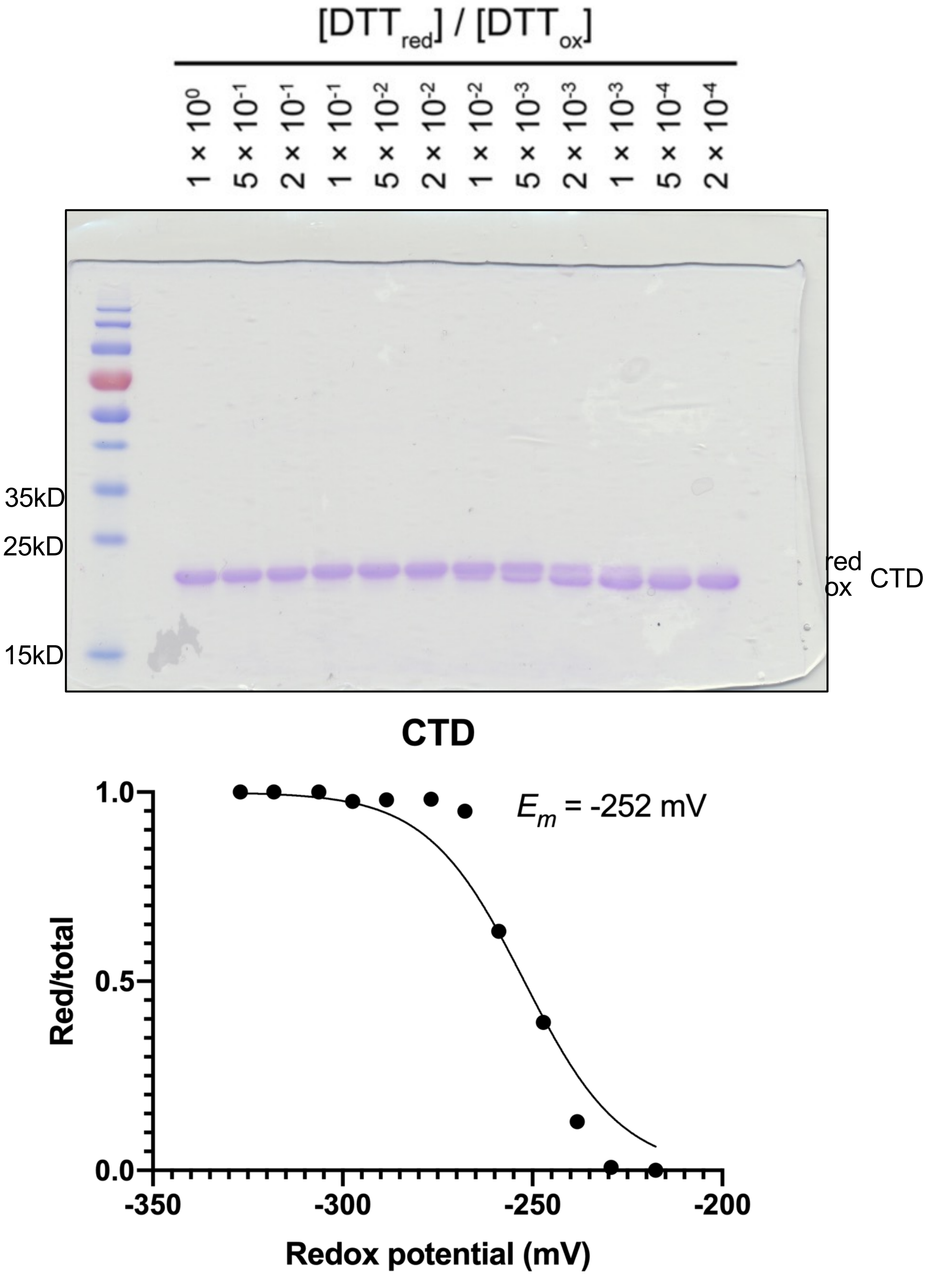


**Figure 2B-C Source Data. The SDS-polyacrylamide gel images of redox titration of CTD. Three Cys in the CTD.** Uncropped version (Replicate number 1)

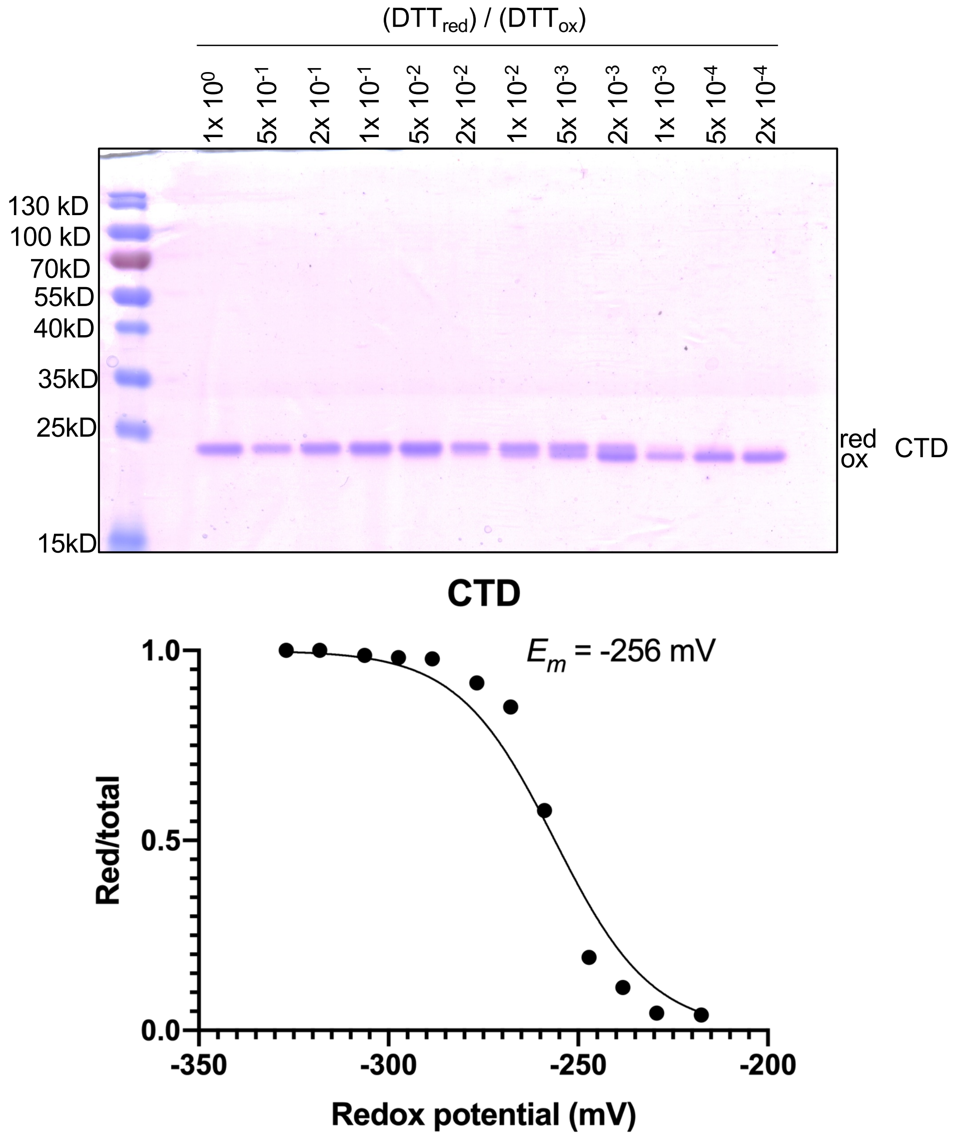


**Supporting Figure 2B-C Source Data. The SDS-polyacrylamide gel images of redox titration of CTD. Three Cys in the CTD.** Uncropped version (Replicate number 2)

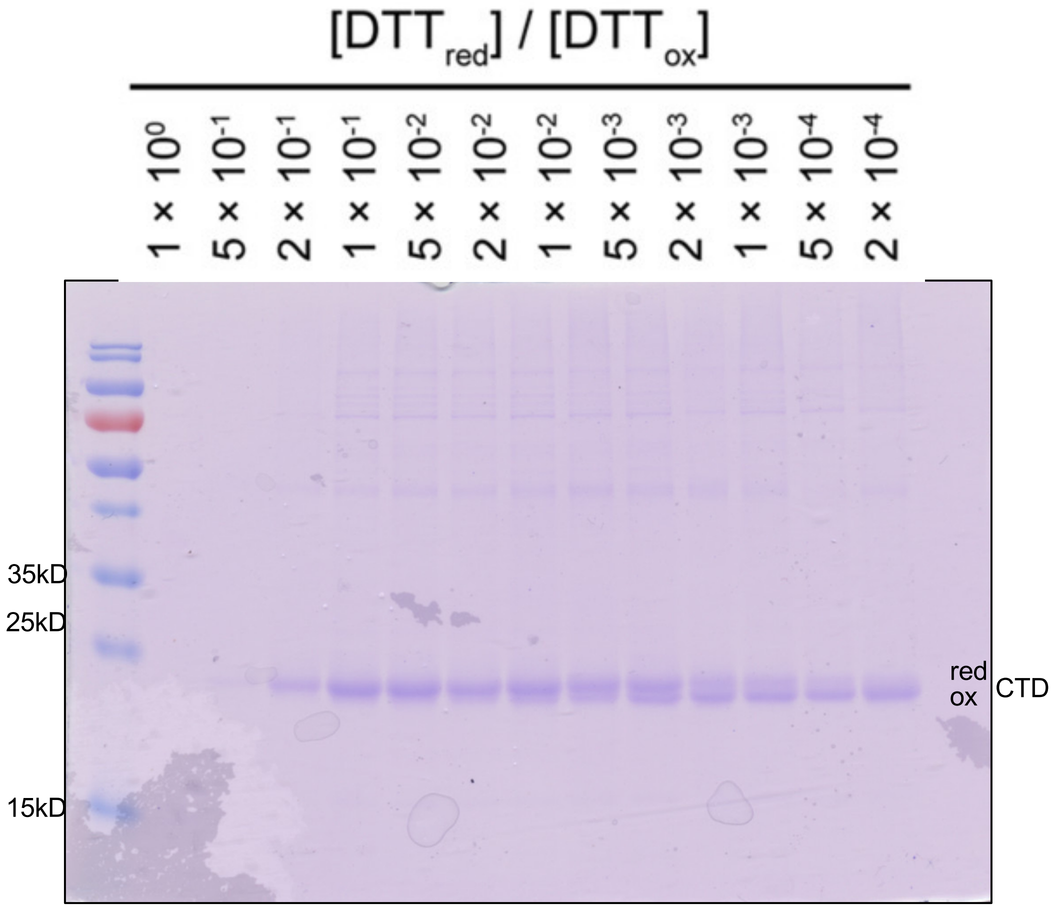


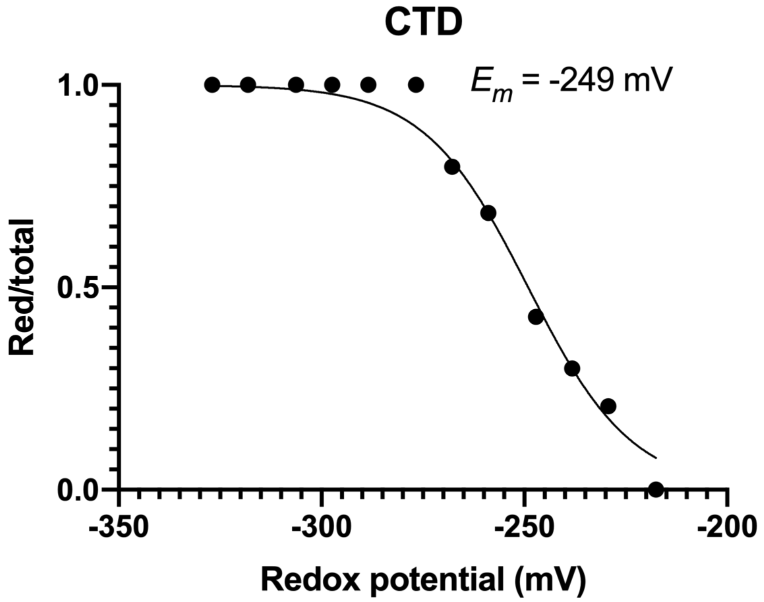


**Supporting Figure 2B-C Source Data. The SDS-polyacrylamide gel images of redox titration of CTD. Three Cys in the CTD.** Uncropped version (Replicate number 3)

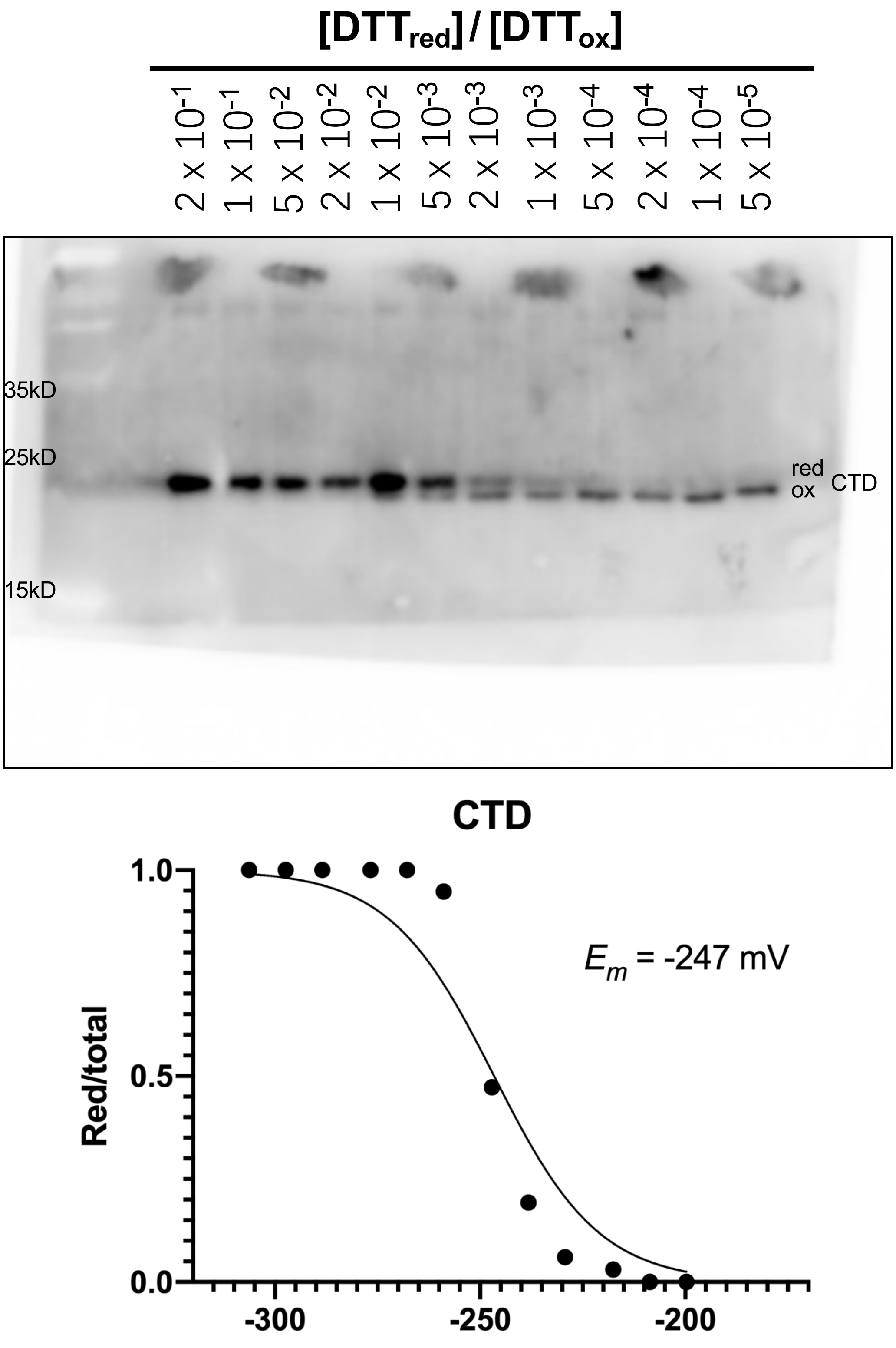


**Supporting Figure 1B-C Source Data. The immunoblot analysis of redox titration of CTD. Three Cys in the CTD.** Uncropped version, PVDF membrane was cut and probed with anti-SOQ1 antibody. (Replicate number 4)

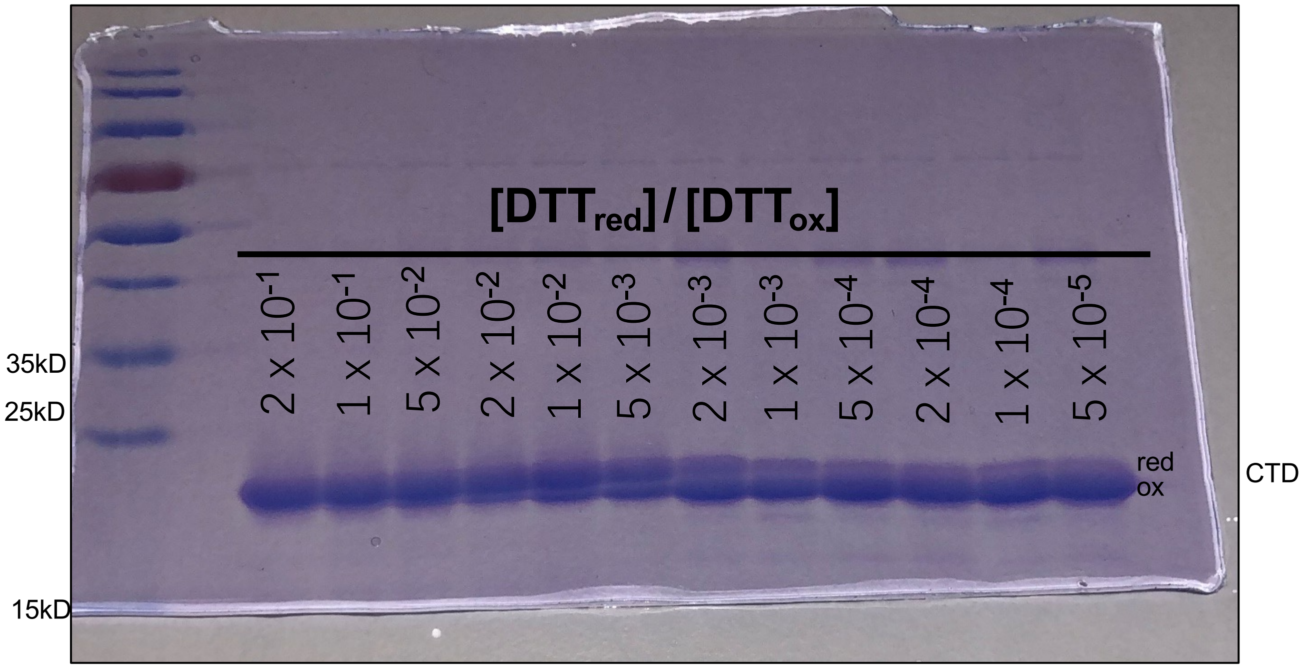


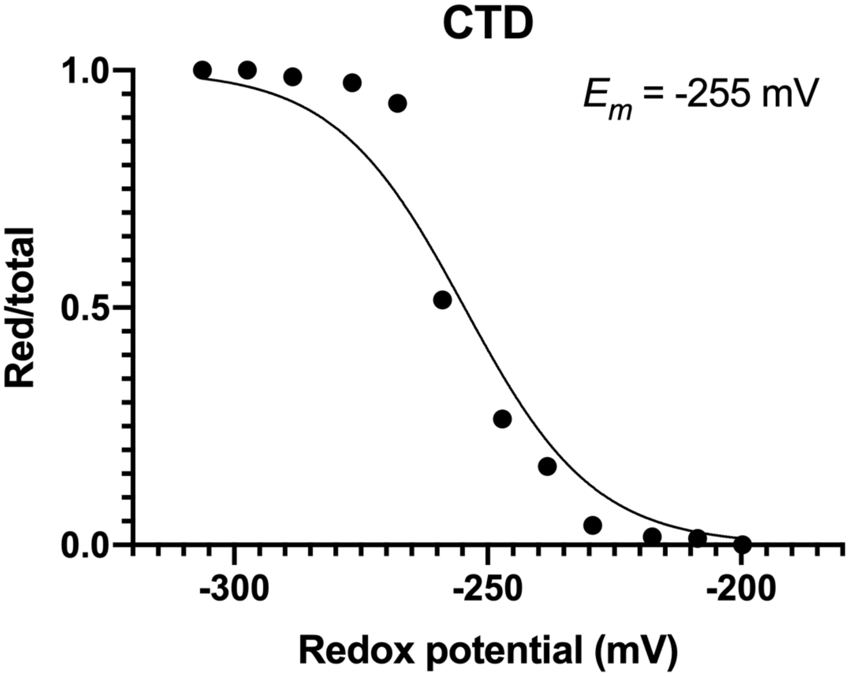


**Supporting Figure 1B-C Source Data. The SDS-polyacrylamide gel images of redox titration of CTD. Three Cys in the CTD.** Uncropped version (Replicate number 5)

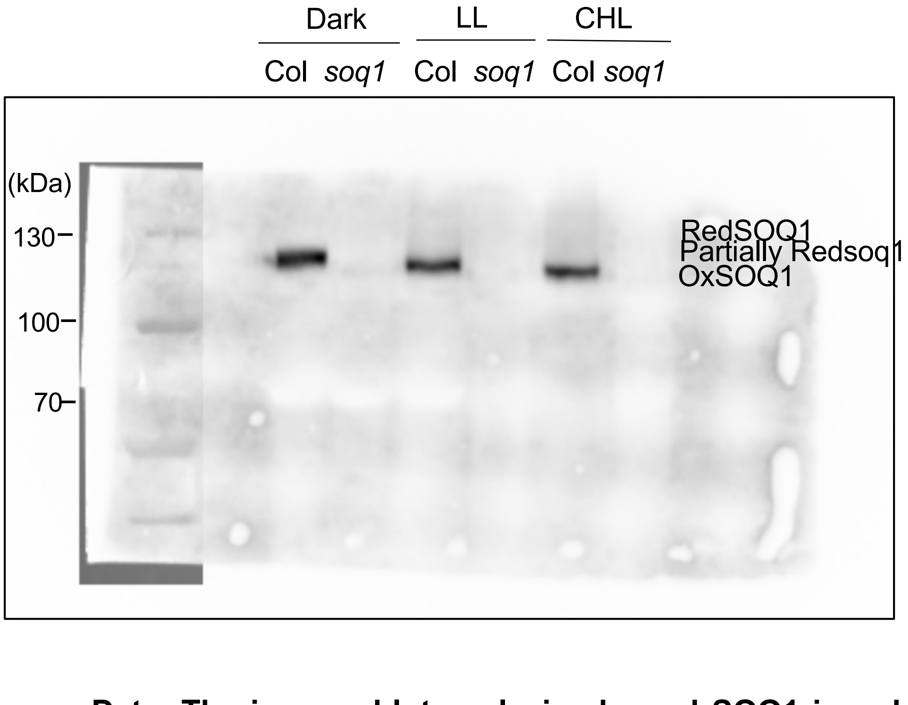


**Figure 2D Source Data. The immunoblot analysis showed SOQ1 is reduced in cold and high light conditions**. Uncropped version, PVDF membrane was probed with anti-SOQ1 antibody. (Replicate number 1)

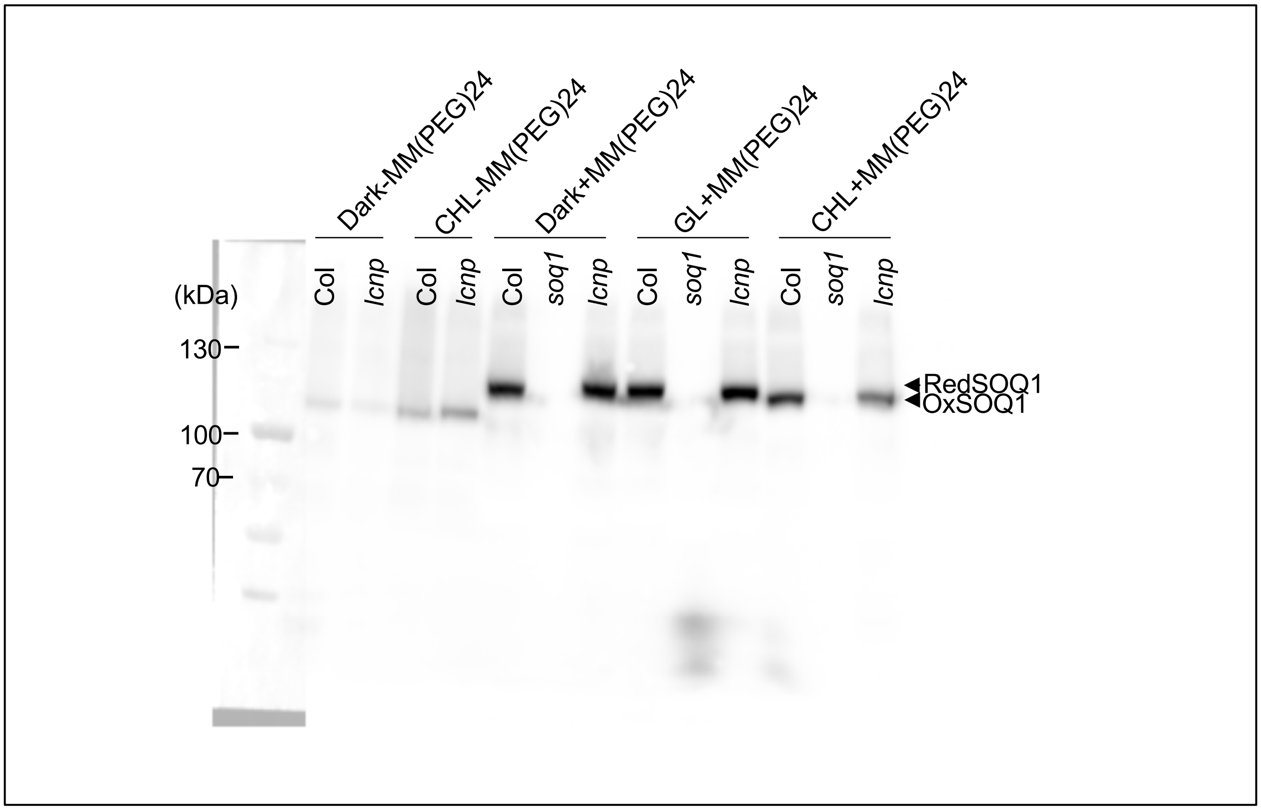


**Supporting Figure 2D Source Data. The immunoblot analysis showed SOQ1 is reduced in cold and high light conditions**. Uncropped version, PVDF membrane was probed with anti-SOQ1 antibody. (Replicate number 2)

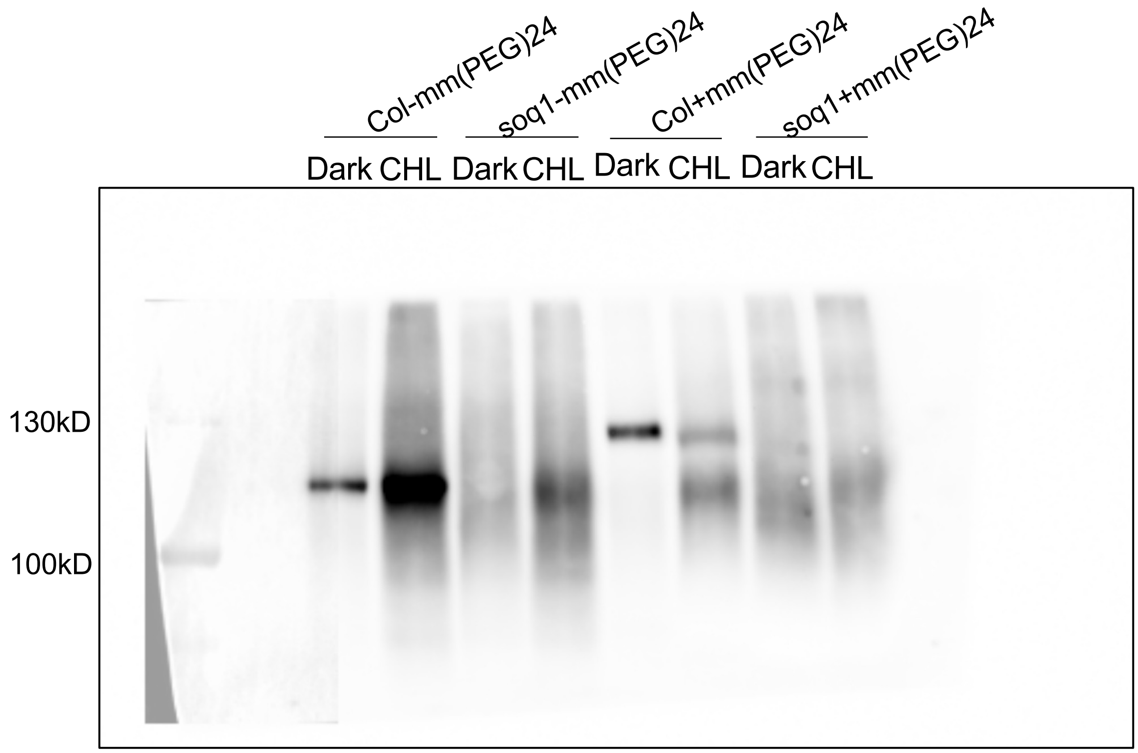


**Supporting Figure 2D Source Data. The immunoblot analysis showed SOQ1 is reduced in cold and high light conditions**. Uncropped version, PVDF membrane was probed with anti-SOQ1 antibody. (Replicate number 3)

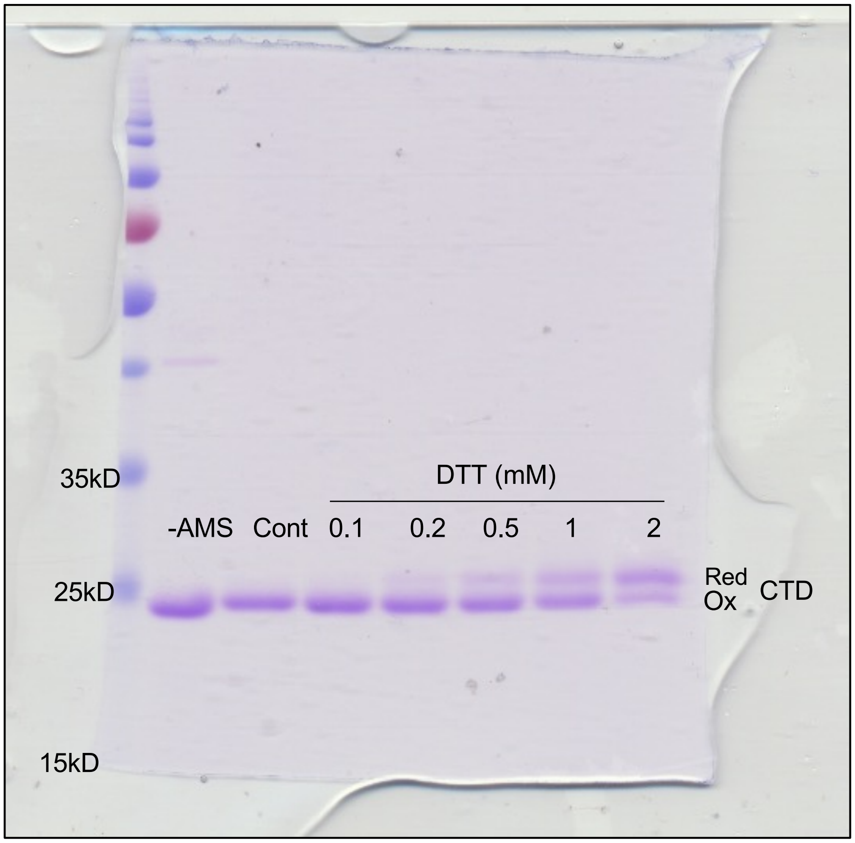


**Figure 3A upper panel Source Data. The SDS-polyacrylamide gel images to , determination of the maximum amount of DTT to mediate the electron transfer reaction. Three Cys in the CTD.** Whole-gel version (Replicate number 1). Support Figure 2A.

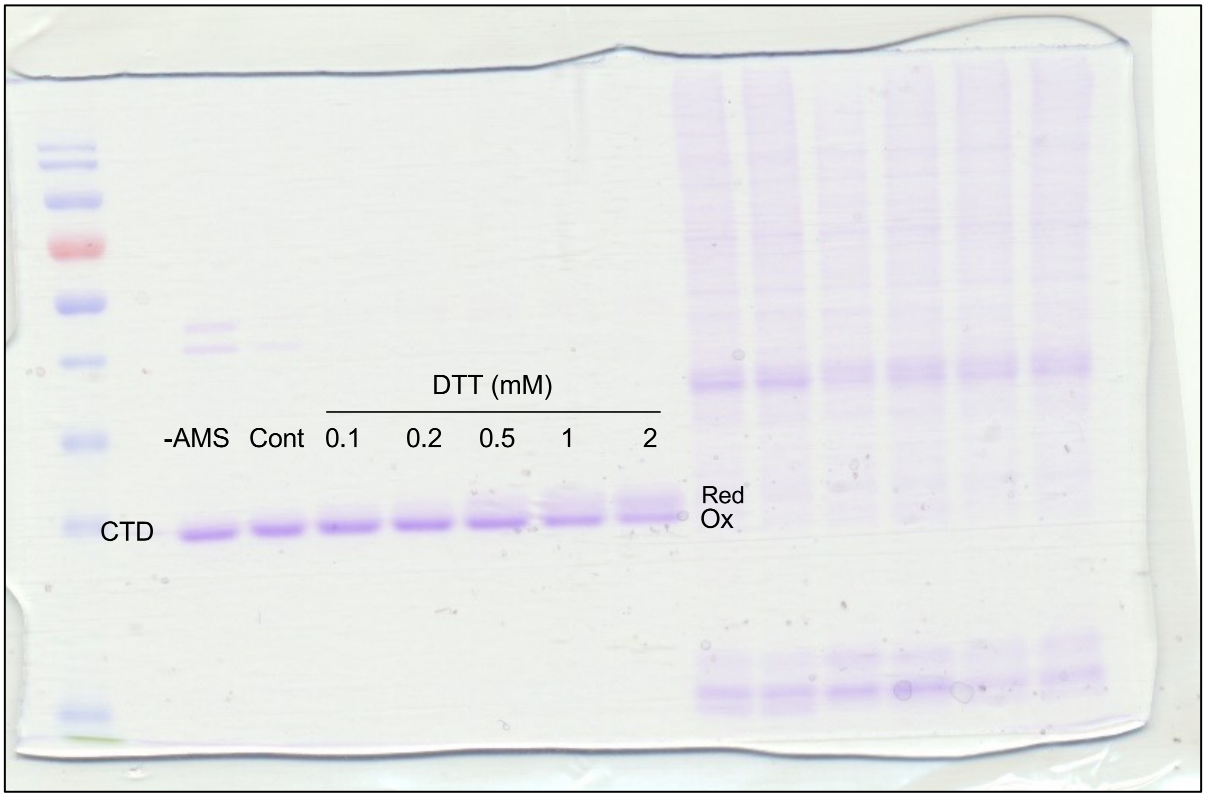


**Supporting Figure 3A Source Data. The SDS-polyacrylamide gel images determination of the maximum amount of DTT to mediate the electron transfer reaction. Three Cys in the CTD**. Whole-gel version (Replicate number 2). Support Figure 2A.

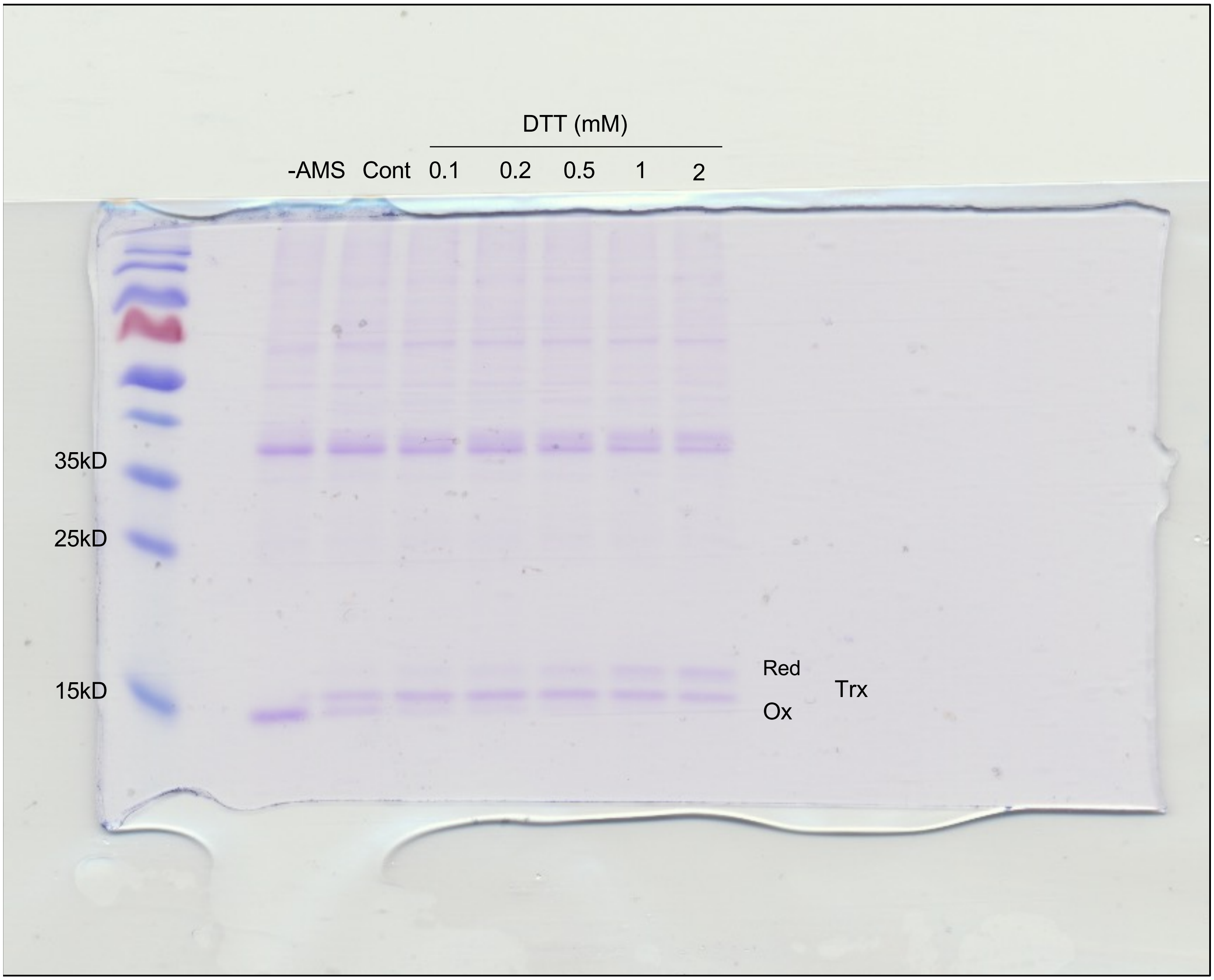


**Figure 3A Bottom panel Source Data. The SDS-polyacrylamide gel images to , determination of the maximum amount of DTT to mediate the electron transfer reaction. Three Cys in the Trx.** Whole-gel version. Support Figure 2A.

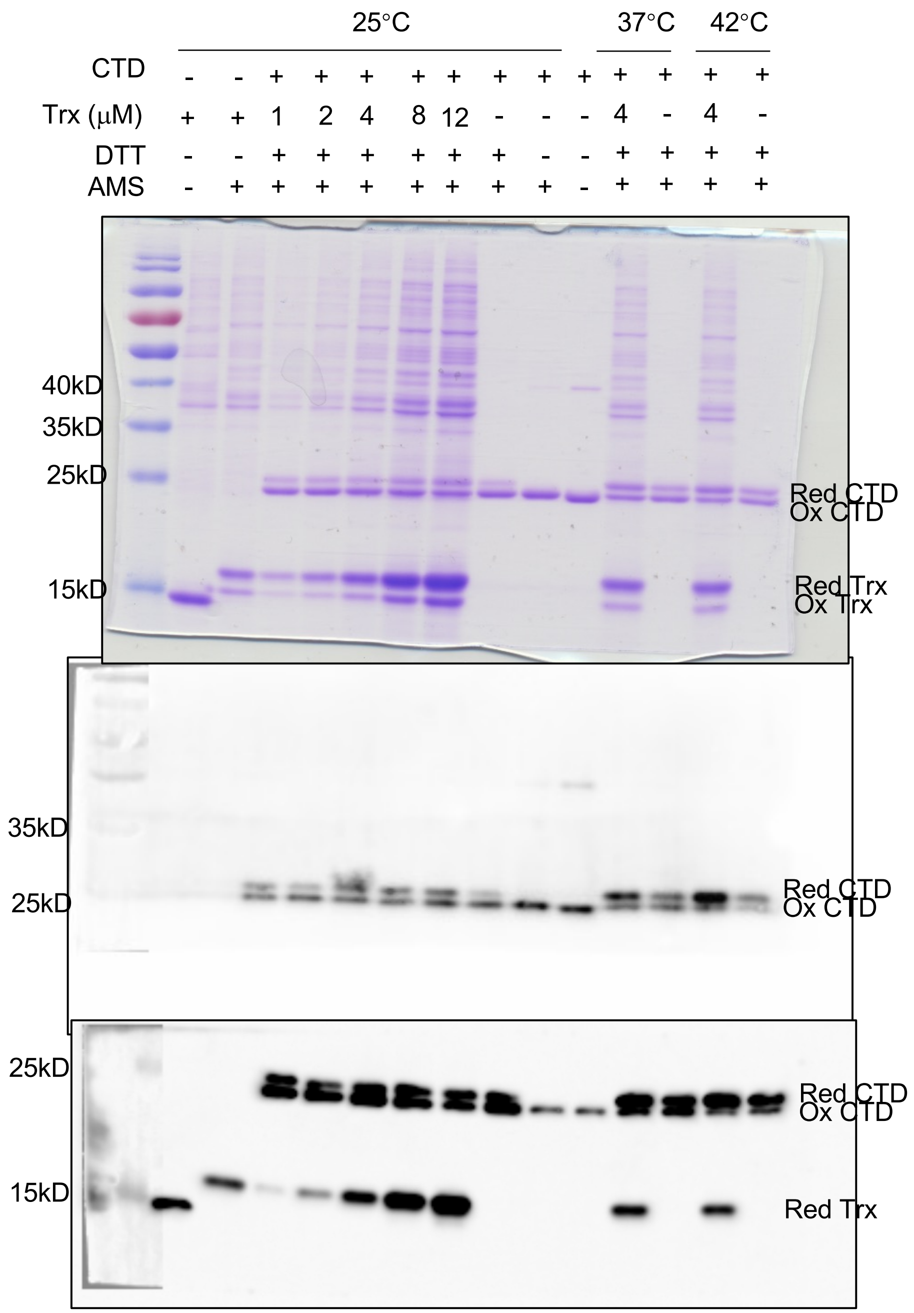


**Figure 3B Source Data. The immunoblot analysis showed Trx can reduce CTD.** Uncropped version, PVDF membrane was cut and probed with anti-SOQ1$_{CTD}$ (middle) and anti-His (bottom) antibodies. (Replicate number 1).

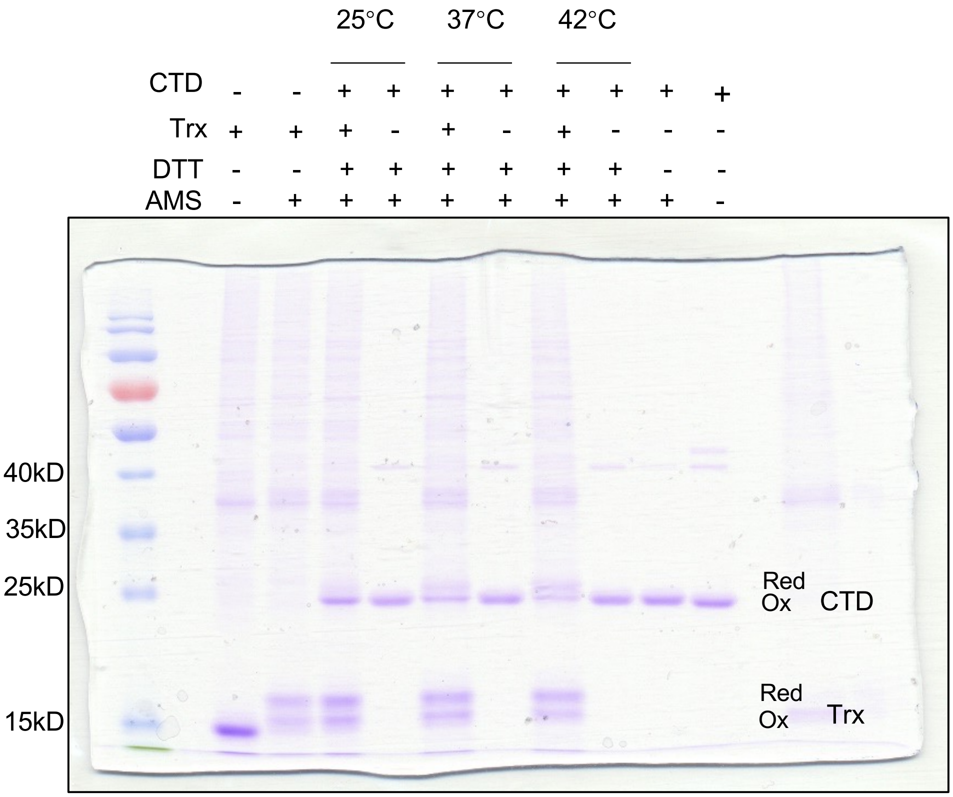


**Supporting Figure 3B Upper Panel Source Data. The SDS-polyacrylamide gel images showed Trx can reduce CTD.** Whole-gel version (Replicate number 2).

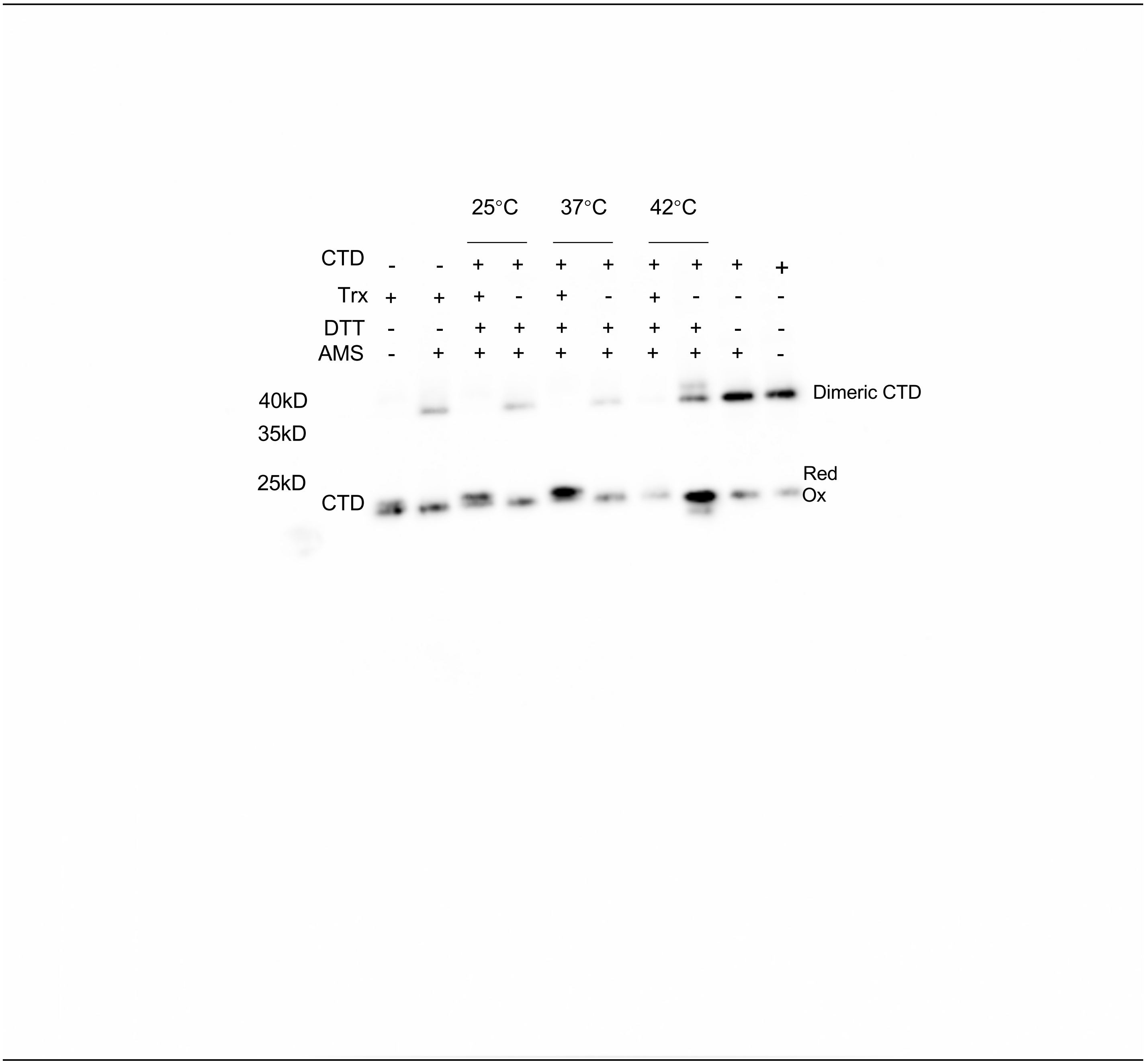


**Supporting Figure 3B Middle Panel Source Data. The immunoblot analysis showed Trx can reduce CTD.** Uncropped version, PVDF membrane was cut and probed with anti-$SOQ1_{CTD}$ antibody. (Replicate number 2).

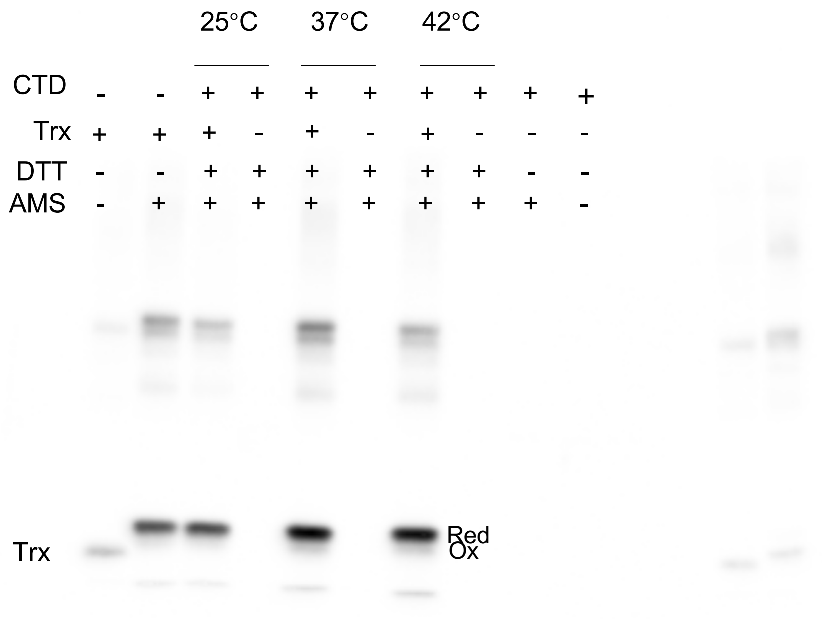


**Supporting Figure 3B Bottom Panel Source Data. The immunoblot analysis showed Trx can reduce CTD.** Uncropped version, PVDF membrane was cut and probed with anti-Trx antibody. (Replicate number 2).

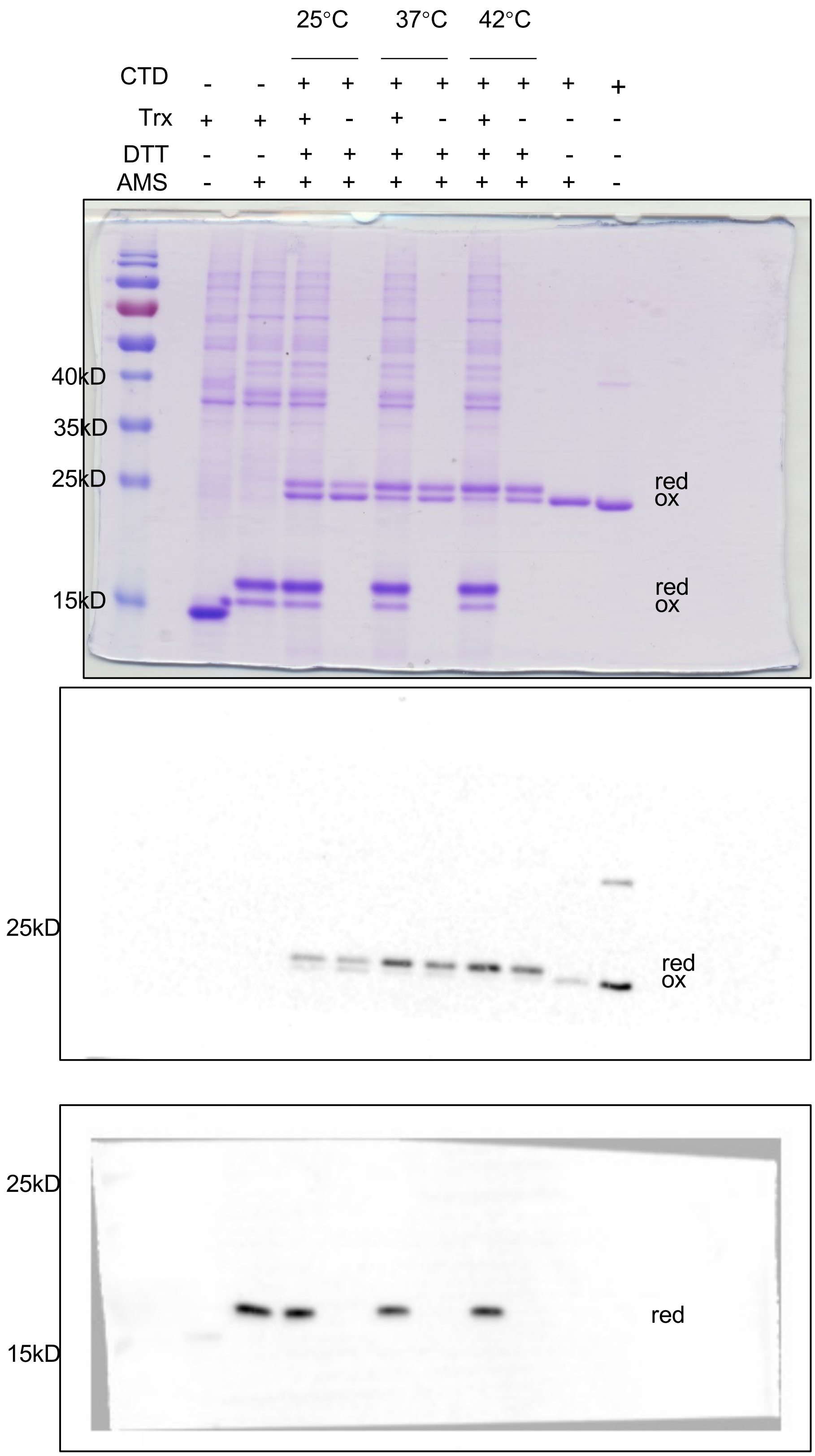


**Supporting Figure 3B Source Data. The immunoblot analysis showed Trx can reduce CTD.** Uncropped version, PVDF membrane was cut and probed with anti-Trx and CTD antibodies. (Replicate number 3).

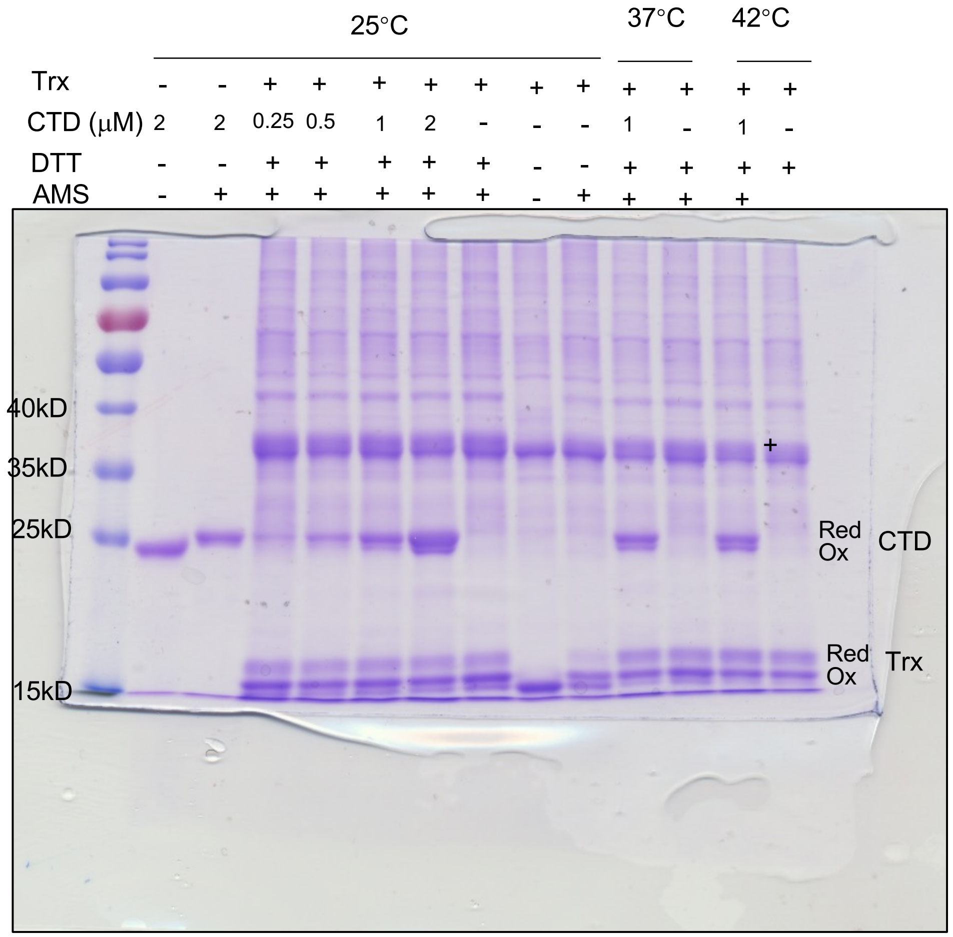


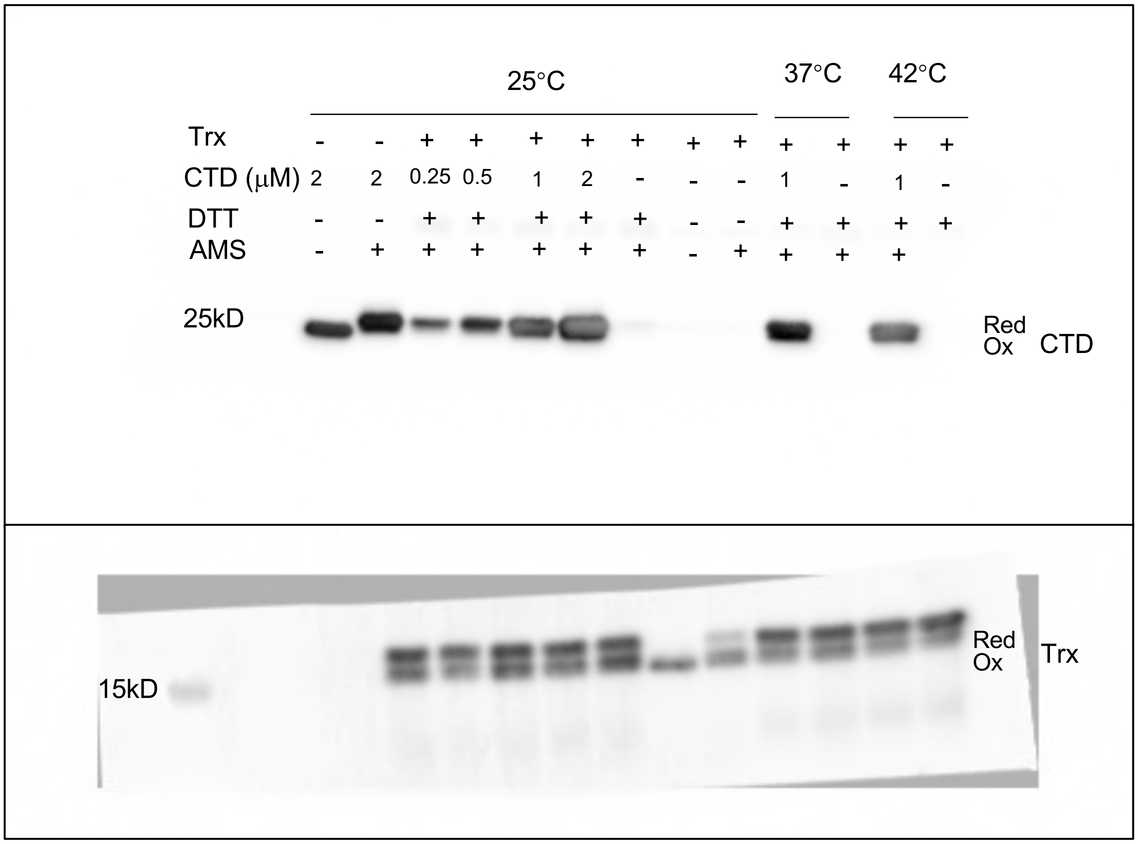


**Figure 3C Source data. CTD can not reduce Trx.** Uncropped version, PVDF membrane was cut and probed with anti-His and anti-Trx antibodies. (Replicate number 1)

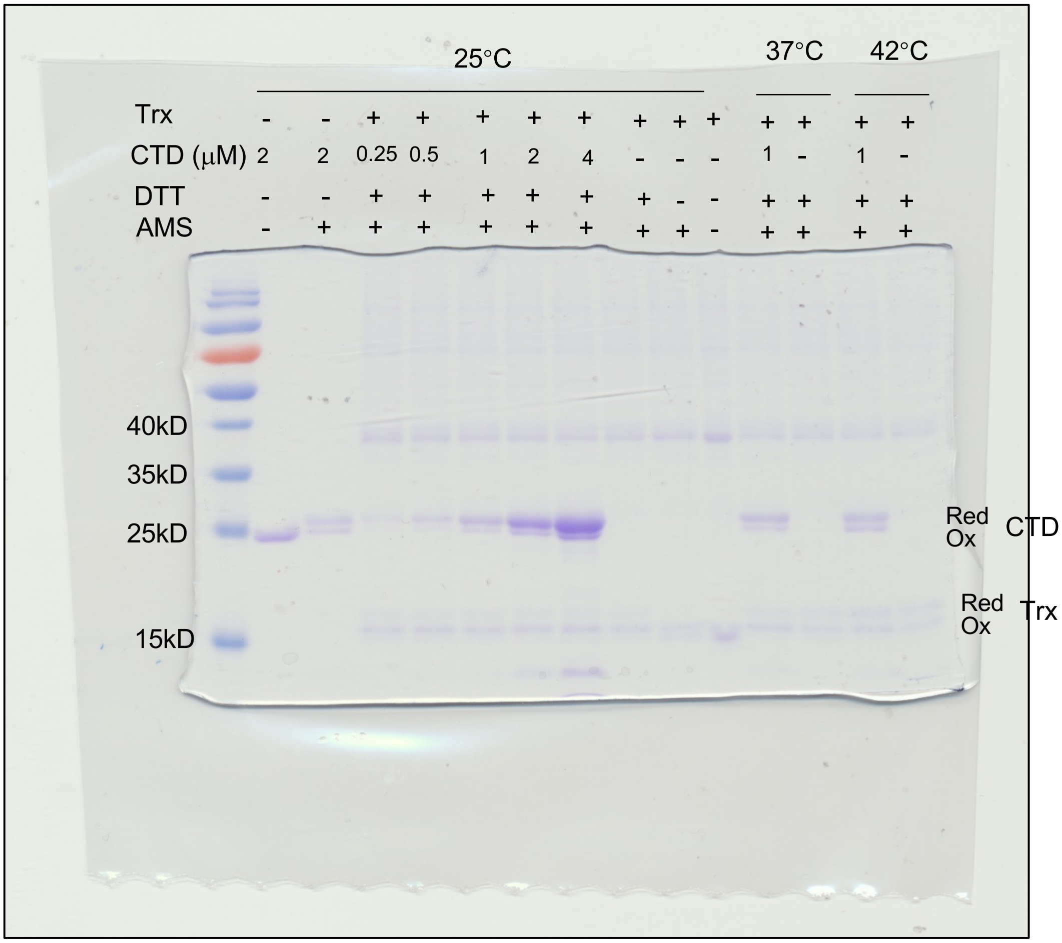


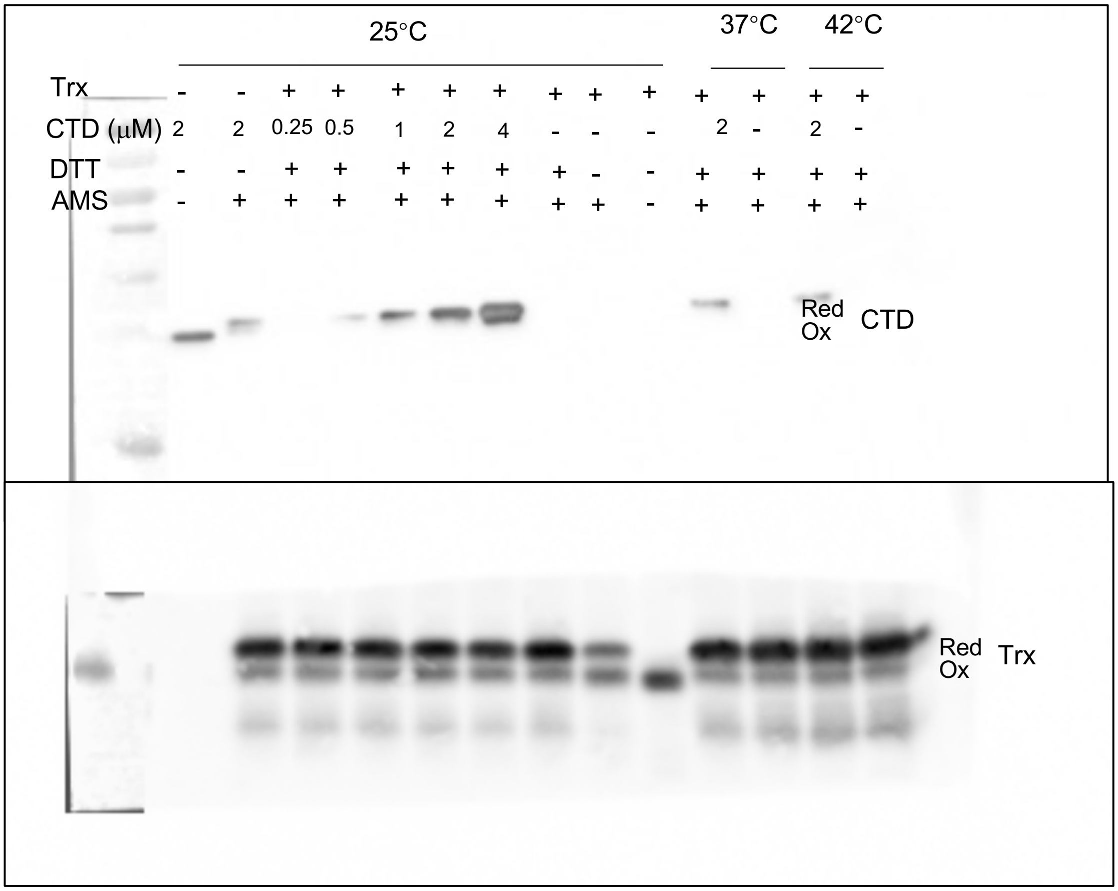


**Supporting Figure 3C Source data. CTD can not reduce Trx.** Uncropped version, PVDF membrane was cut and probed with anti-His and anti-Trx antibodies. (Replicate number 2**)**

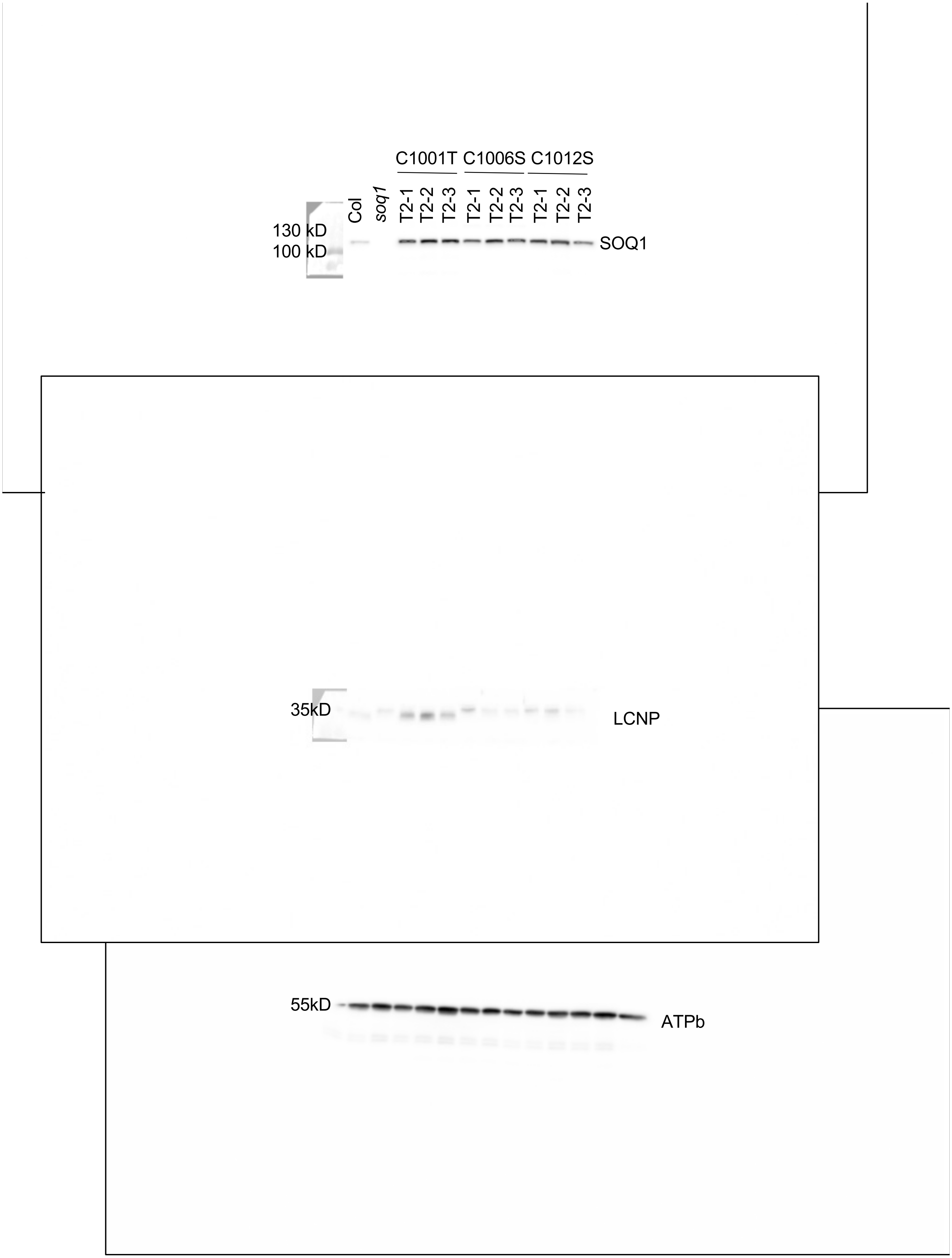


**Figure 4A Source Data. Immunoblot analysis showed the acccumulation of C1001T, C1006S and C1012S in three individual lines.** Uncropped version. PVDF Membrane was cut and probed with anti-SOQ1, anti-LCNP and anti-ATPb antibodies. (Replicate number 1)

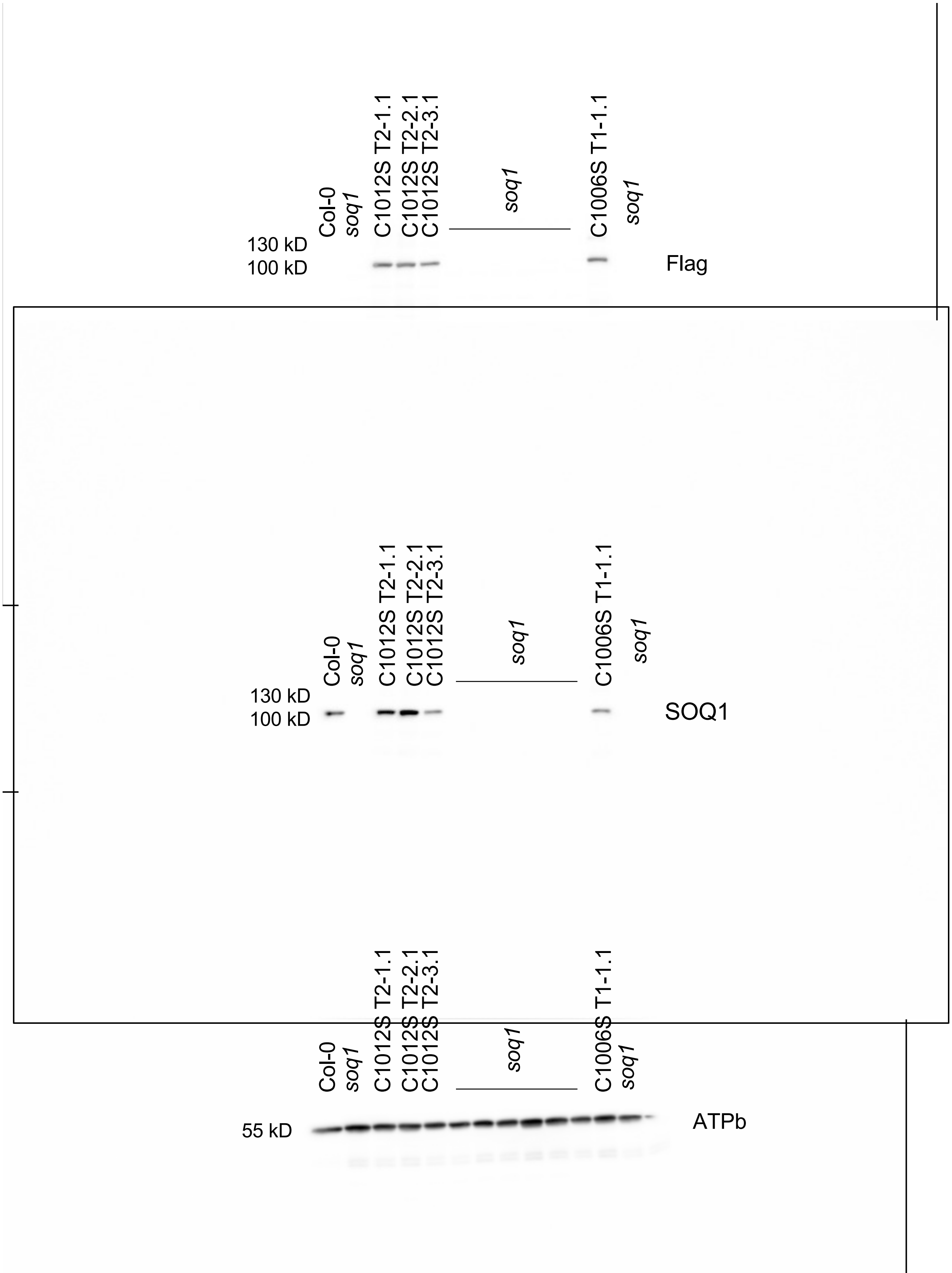


**Supporting Figure 4A Source Data. Immunoblot analysis showed the acccumulation of C1001T, C1006S and C1012S.** Uncropped version. PVDF Membrane was cut and probed with anti-SOQ1, anti-Flag and anti-ATPb antibodies. (Replicate number 2)

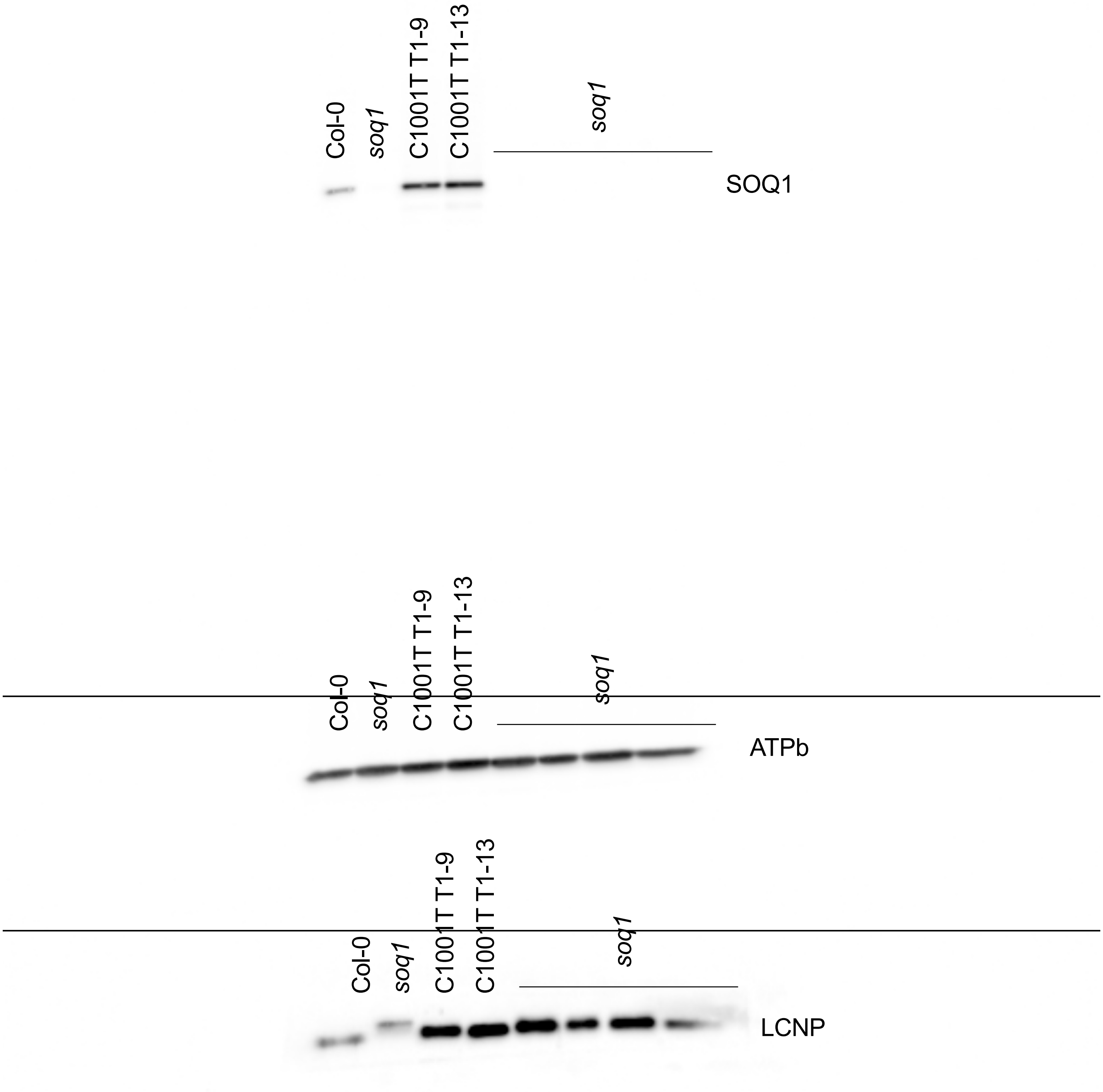


**pporting Figure 4A Source Data. Immunoblot analysis showed the acccumulation of C1001T, C**
**d C1012S in three individual lines.** Uncropped version. PVDF Membrane was cut and probed wi
Q1, anti-LCNP and anti-ATPb antibodies. (Replicate number 2)

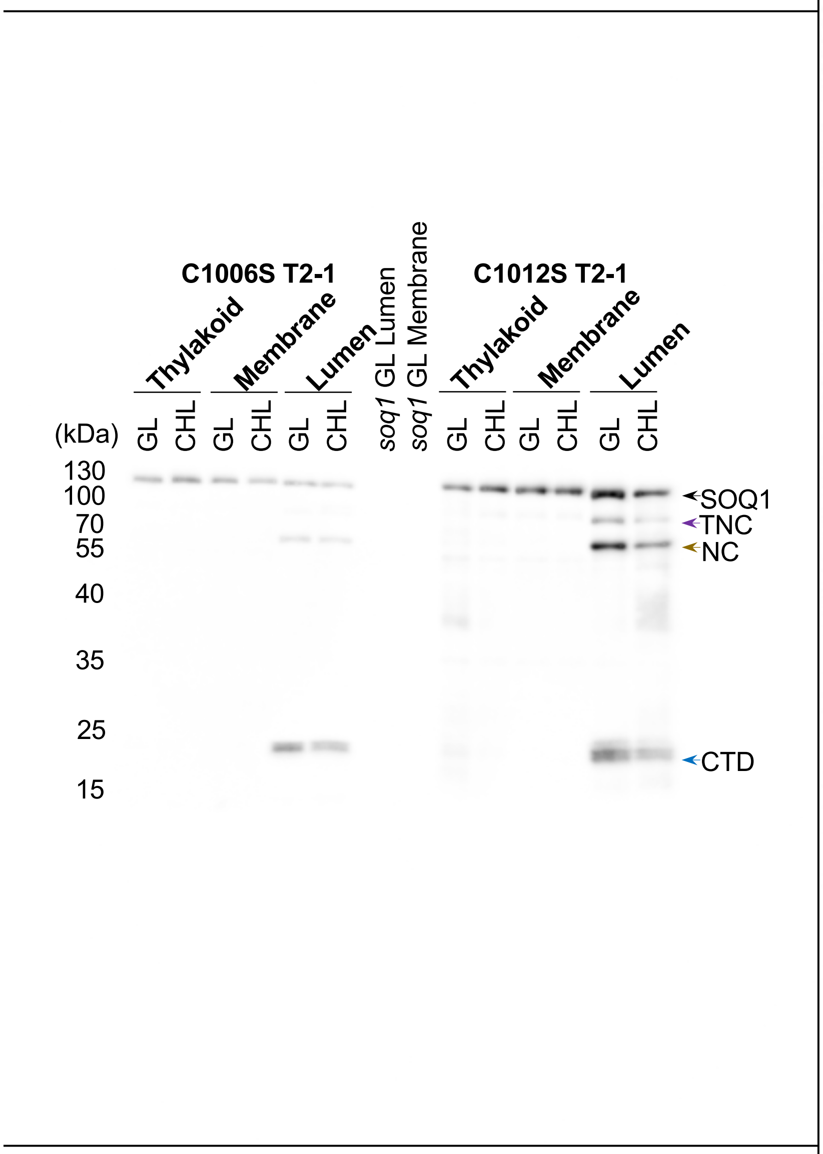


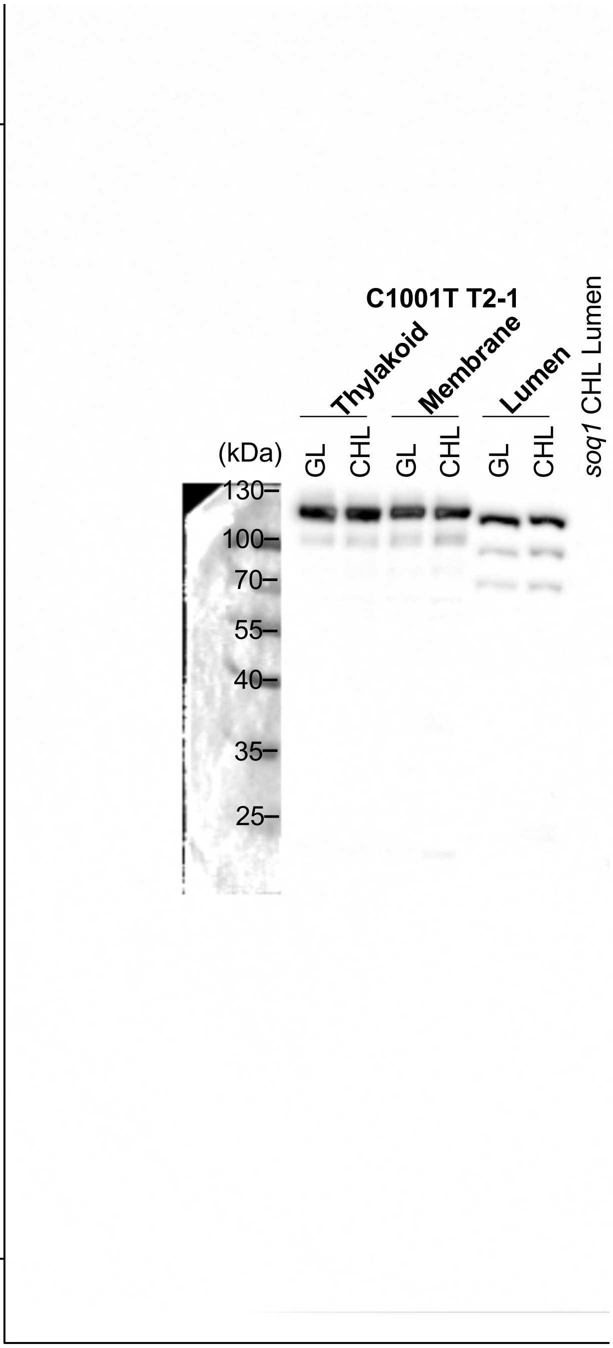


**Figure S4A and S4B Upper Panel Source Data. Immunoblot analysis showed the localization of C1001T, C1006S and C10012S.** Uncropped version. PVDF Membrane was probed with anti- $SOQ1_{CTD}$ antibody. (Replicate number 1)

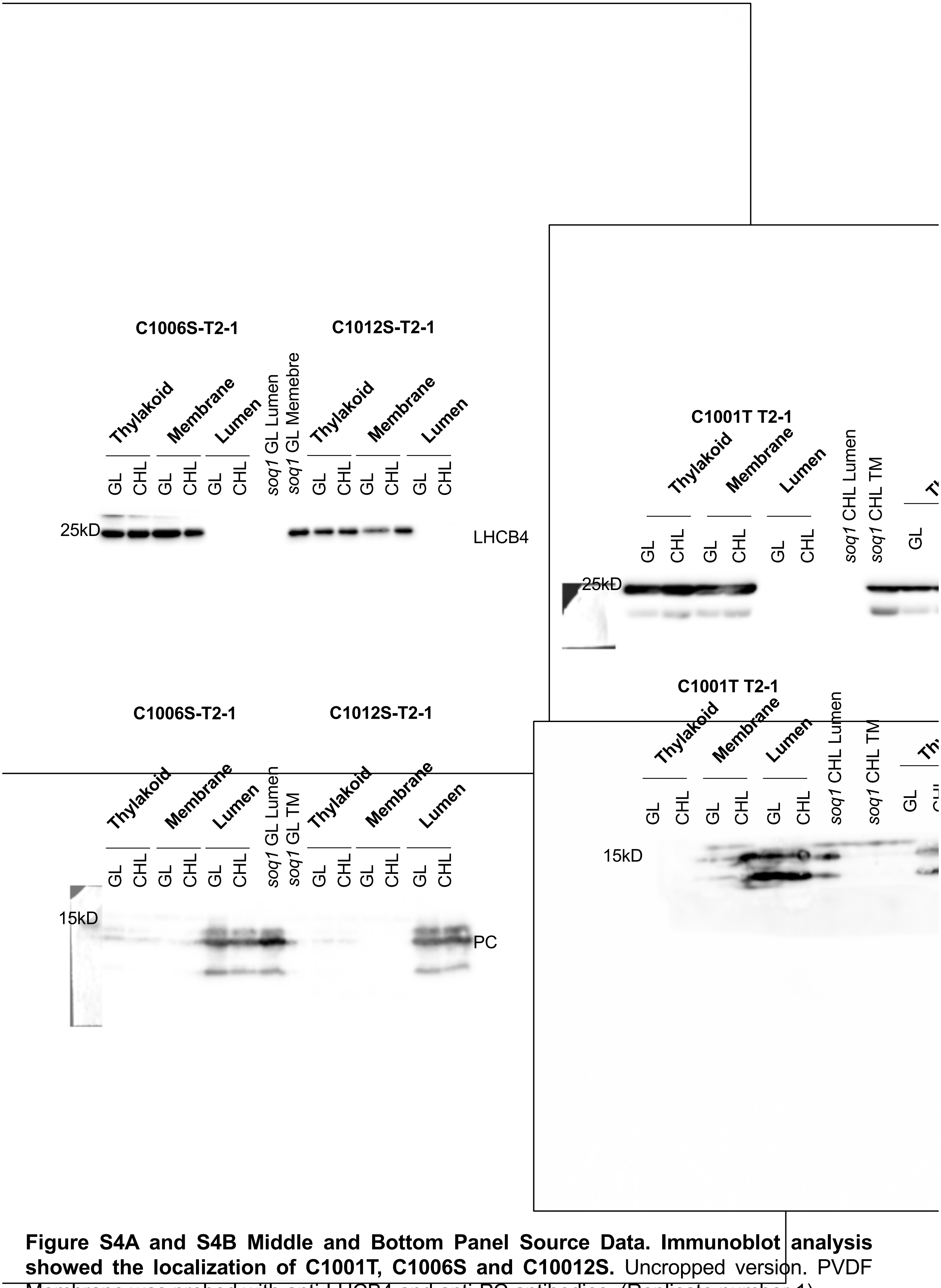


**Figure S4A and S4B Middle and Bottom Panel Source Data. Immunoblot analysis showed the localization of C1001T, C1006S and C10012S.** Uncropped version. PVDF Membrane was probed with anti-LHCB4 and anti-PC antibodies. (Replicate number 1)

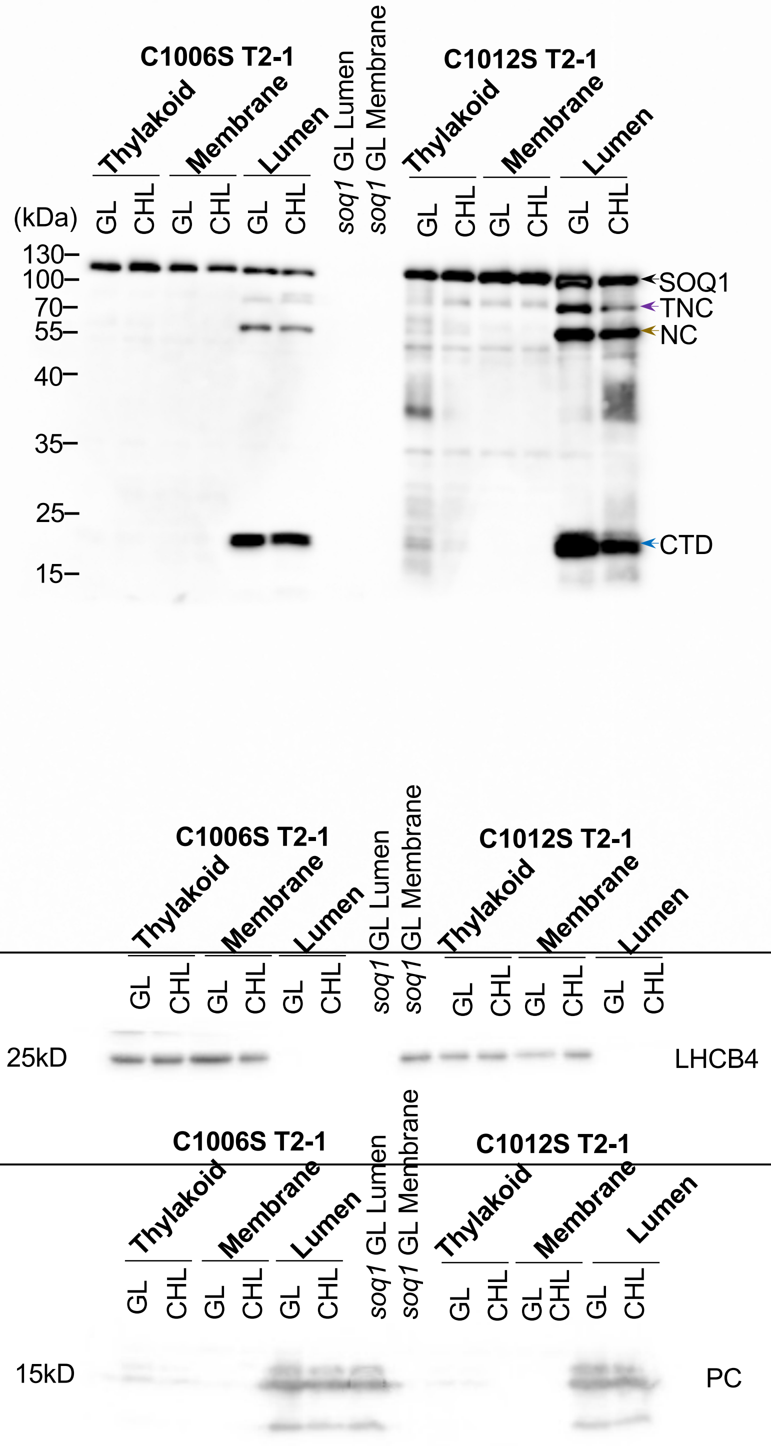


**Supporting Figure S4A Upper Panel Source Data. Immunoblot analysis showed the localization of C1006S and C10012S.** Uncropped version. PVDF Membrane was probed with anti-Flag, anti-LHCB4 and PC antibodies. (Replicate number 2)

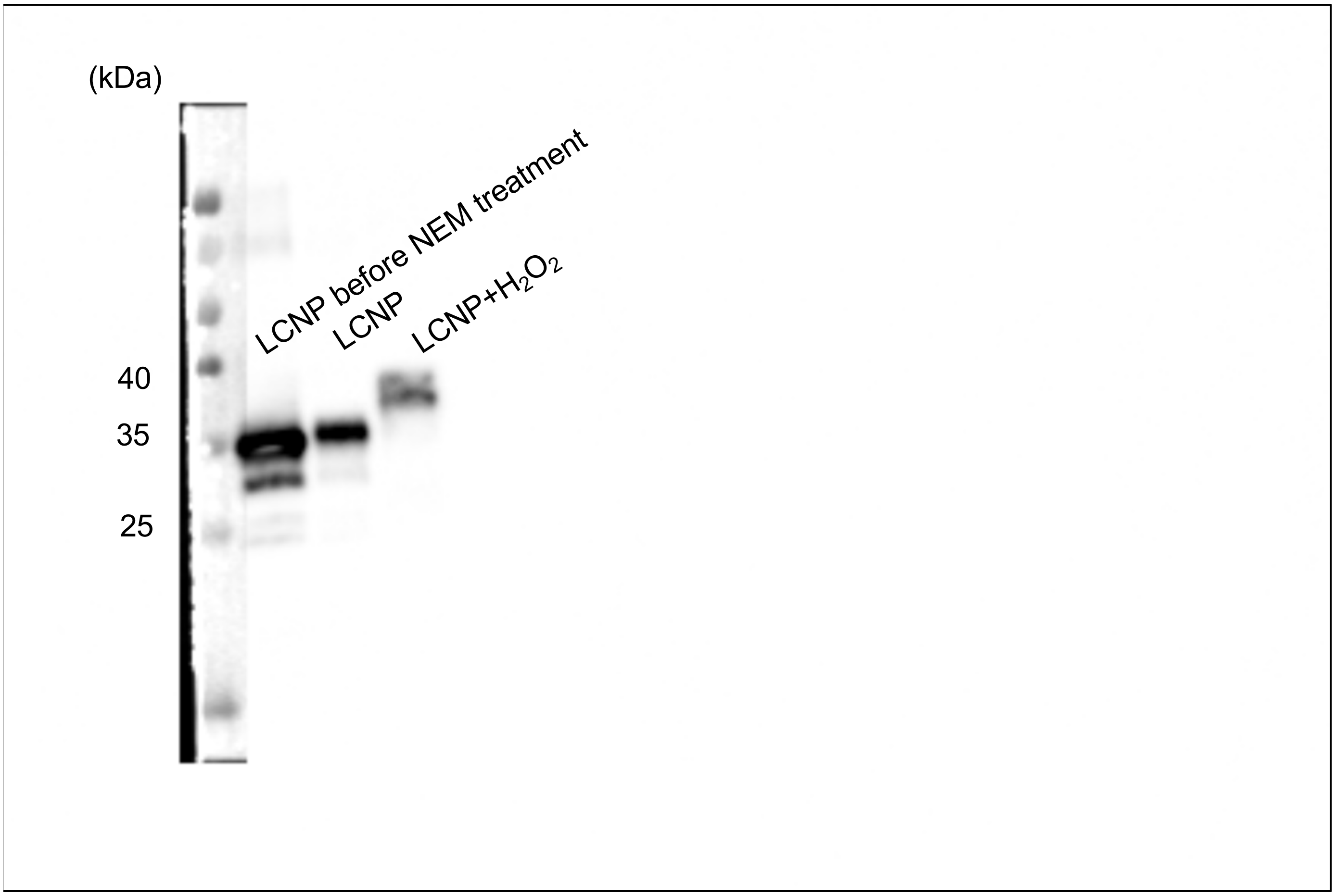


**Figure 5A Source Data. The immunoblot analysis of LCNP before NEM treatment, LCNPand LCNP-H2O2.** Uncropped version, PVDF membrane was probed with anti-LCNP antibody.

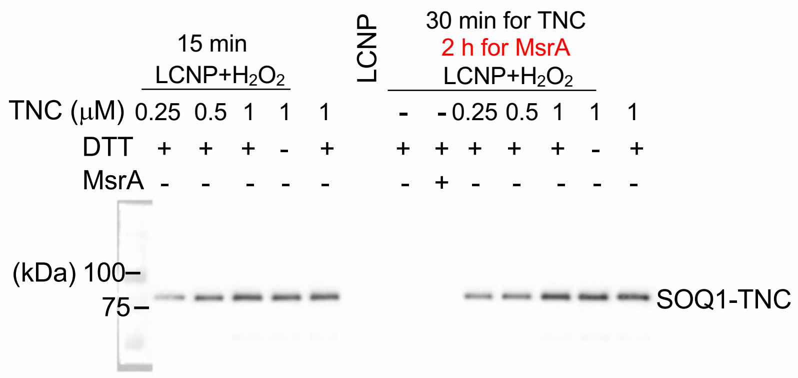


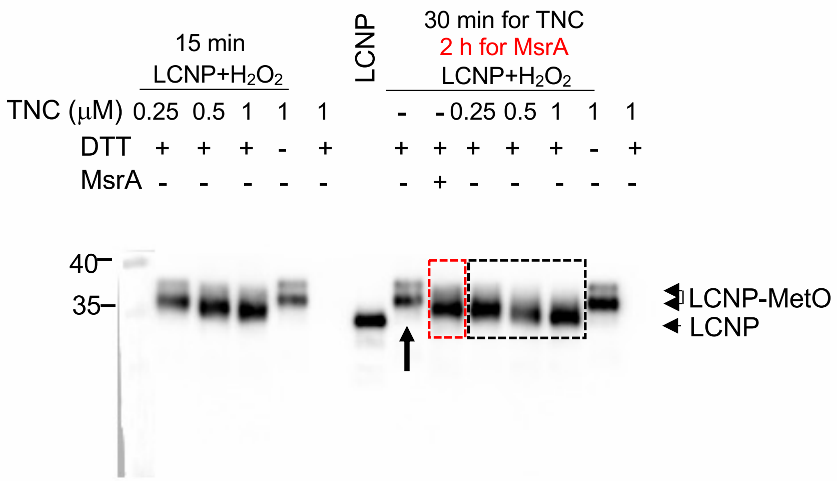


**e 5B Source Data. The immunoblot analysis showed recombinant TNC can reduce -MetSO to LCNP-Met.** Uncropped version, PVDF membrane was probed with anti-LCNP nti-SOQ1$_{CTD}$ antibodies. (Replicate number 1)

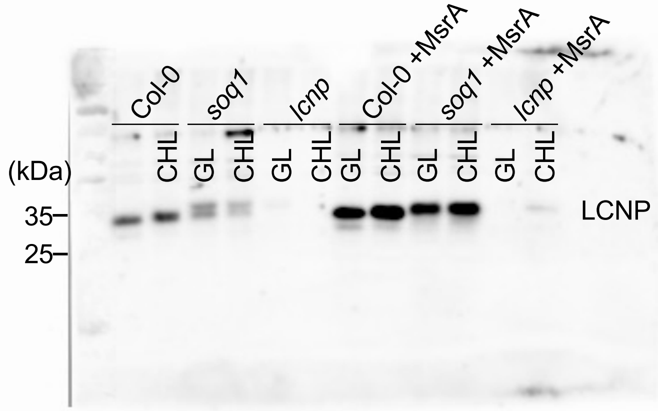


**ɡure 5C Source Data. Incubation of lumenal proteins with MsrA.** Uncropped version, PVDF ɘmbrane was probed with anti-LCNP antibody.

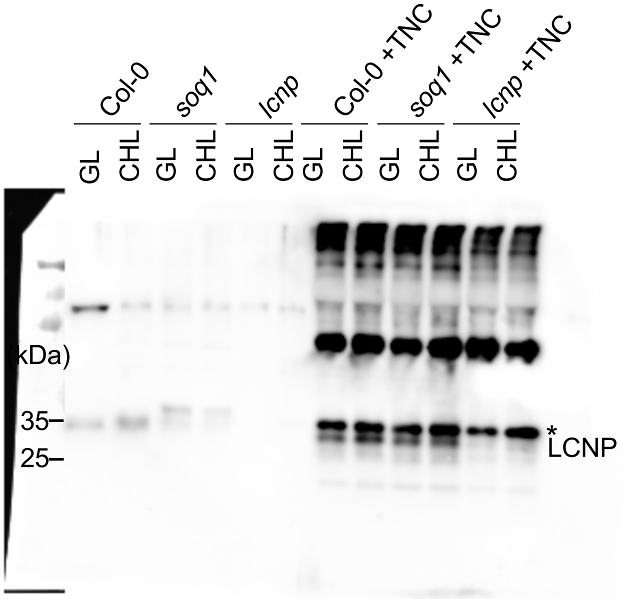


**5D Source Data. Incubation of lumenal proteins with recobimant TNC.** Star symbol (*) ɘnts a nonspecific band, based on its presence in lcnp, detected by the a-LCNP antibody ɔriginating from the TNC recombinant protein preparation. Uncropped version, PVDF rane was probed with anti-LCNP antibody.

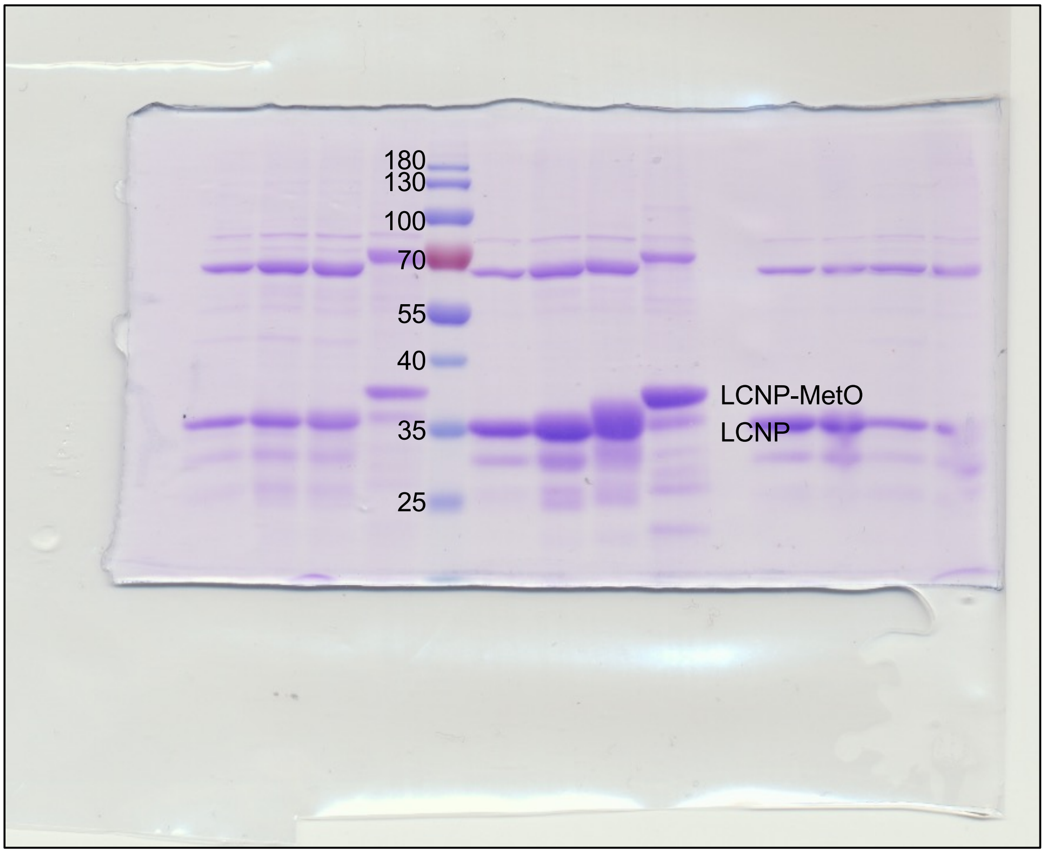


**Figure S5A Source Data.** , $H_2O_2$-treatment of recombinant LCNP $_{A104.}$ Whole-gel version.

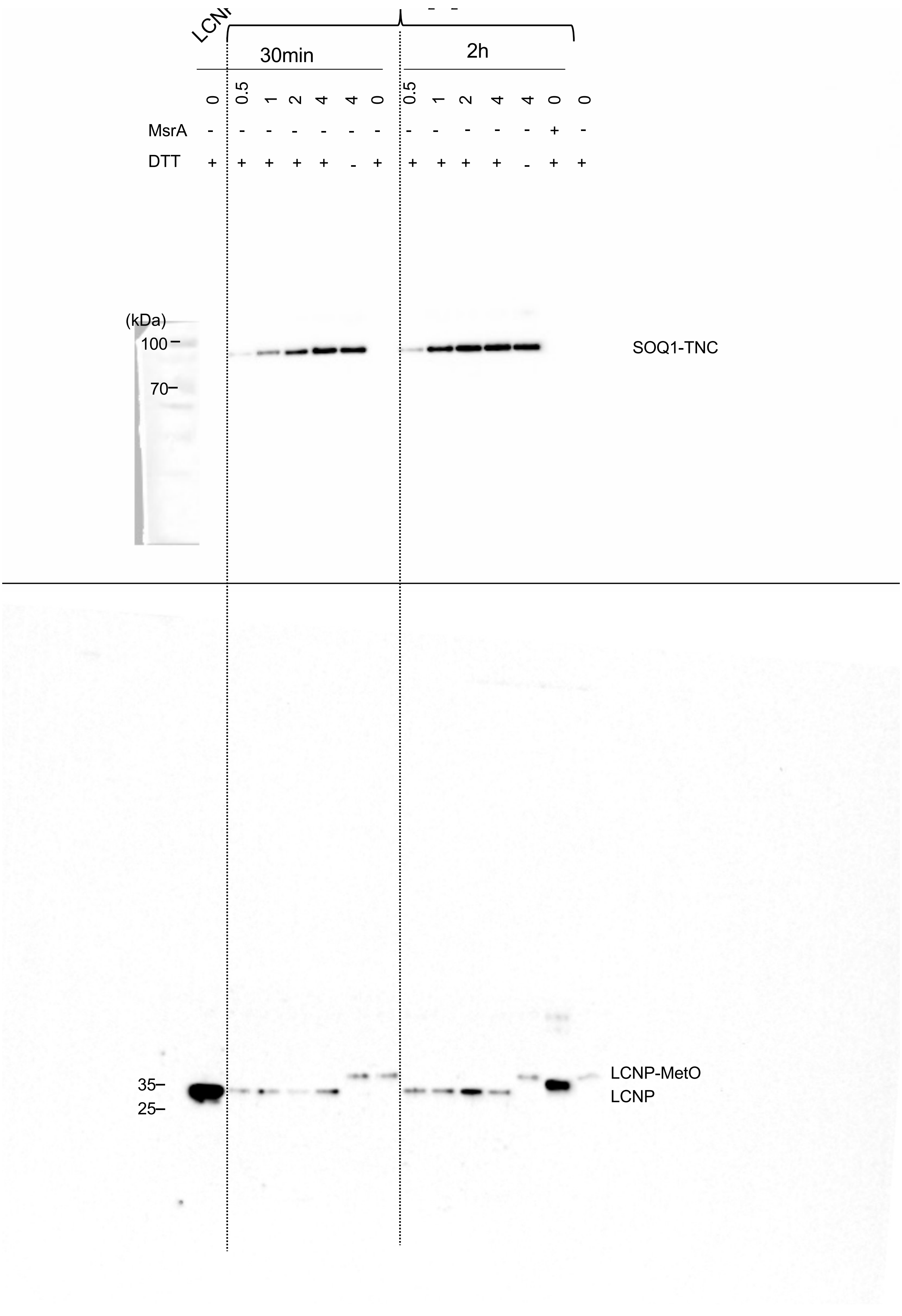

LCNP
30min
2h
0
0.5
1
2
4
4
0
0.5
1
2
4
4
0
0
MsrA
DTT
(kDa)
100
70
SOQ1-TNC
35
25
LCNP-MetO
LCNP

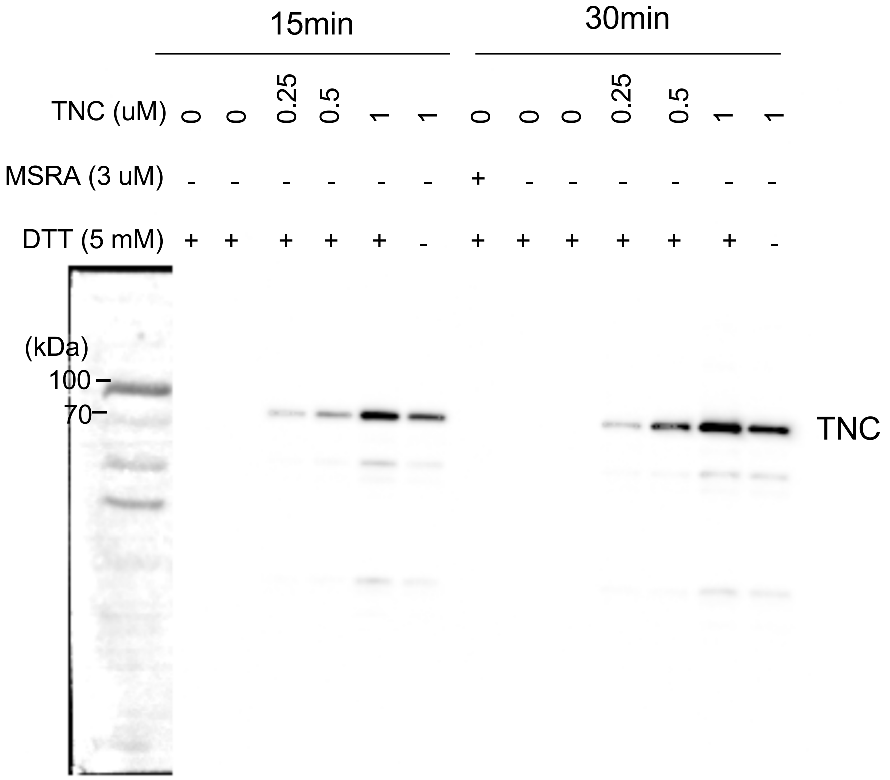


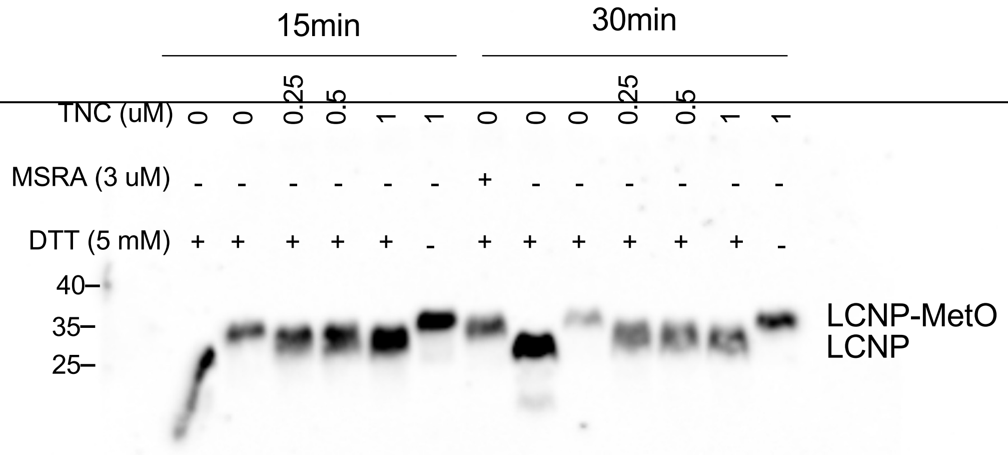


**Supporting Figure S5B Source Data. The immunoblot analysis showed recombinant TNC can reduce LCNP-MetSO to LCNP-Met.** Uncropped version, PVDF membrane was probed with [illegible]

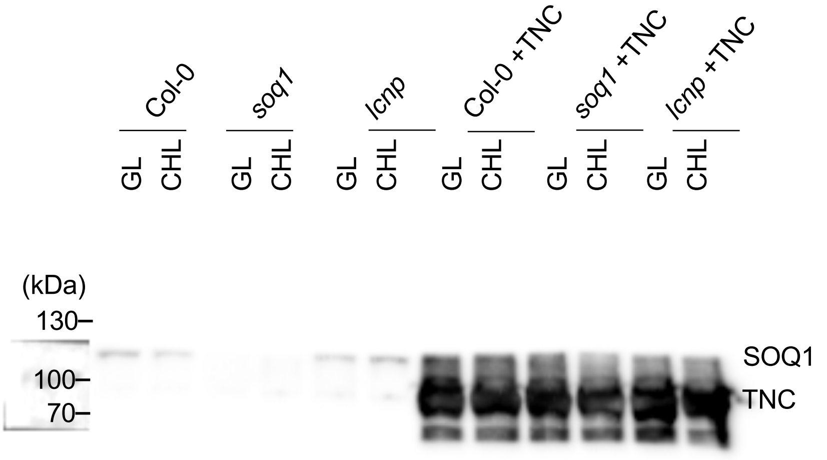


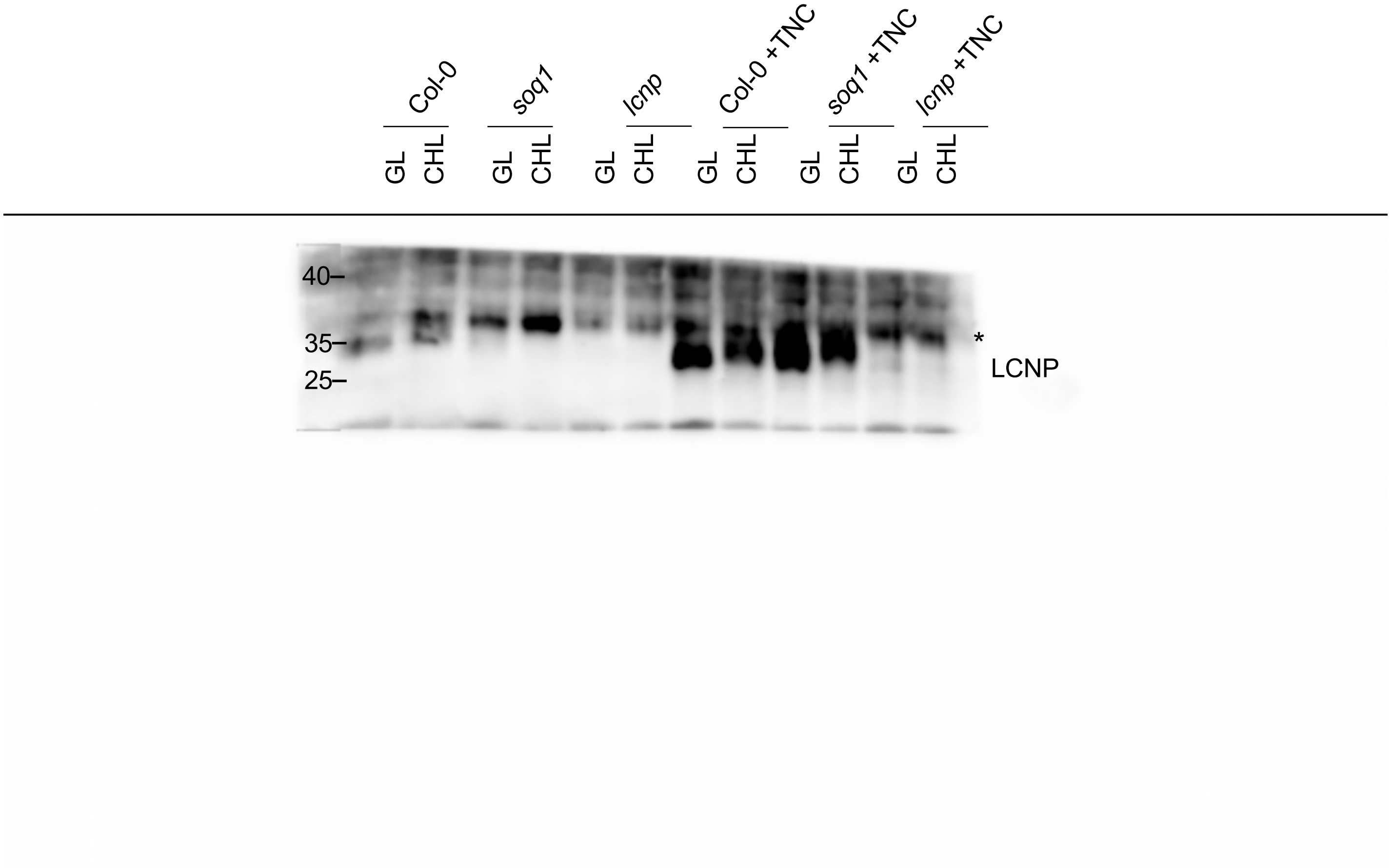


**Figure S5C Source Data. The immunoblot analysis showed recombinant TNC can reduce the shifted LCNP in *soq1* mutant.** Star symbol (*) represents nonspecific band detected by the anti-LCNP antibody, based on its absence in *lcnp* lumen samples. Uncropped version, PVDF membrane was probed with anti-SOQ1 $_{CTD}$ and anti-LCNP antibodies.

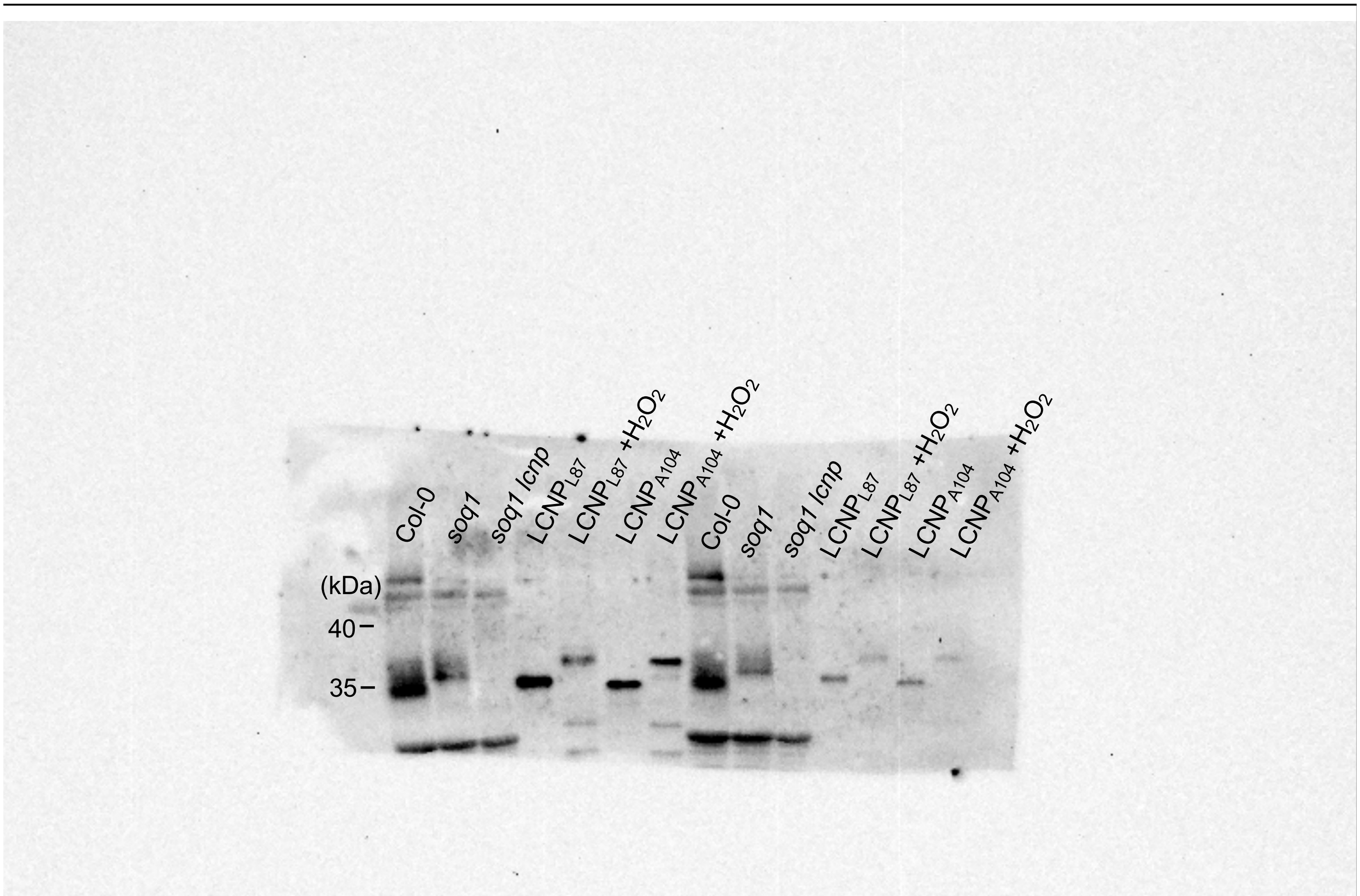


**Figure S5D Source Data. The slower electrophoretic mobility of LCNP in *soq1* is likely due to oxidation.** Uncropped version, PVDF membrane was probed with anti-LCNP antibody.

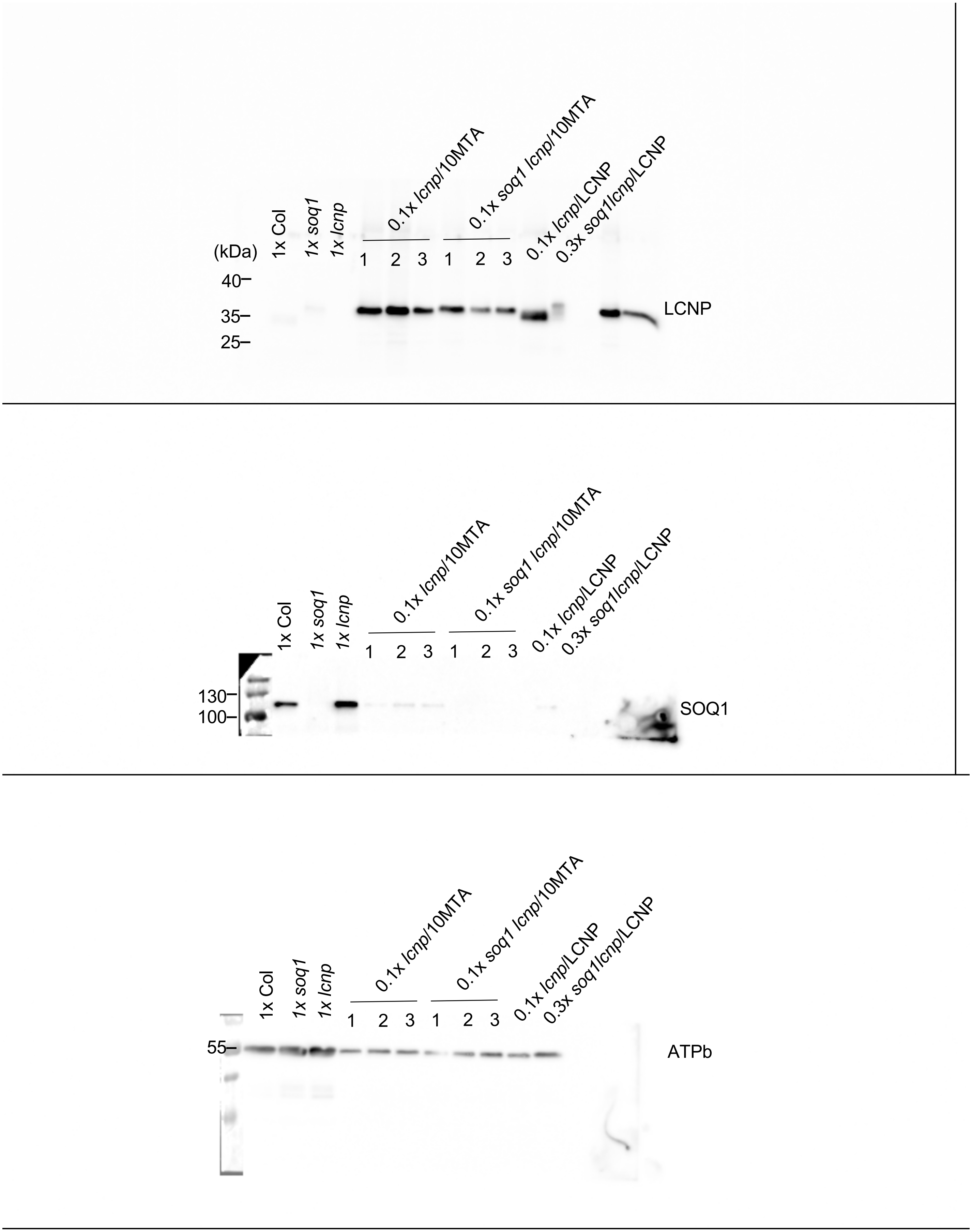


**Figure S6A Source Data. The immunoblot analysis of LCNP with 10 Met mutated to Ala in *lcnp* and *soq1 lcnp* backgroud.** Uncropped version, PVDF membrane was cut and probed with anti-LCNP, anti-SOQ1$_{CTD}$ and anti-ATPb antibodies.

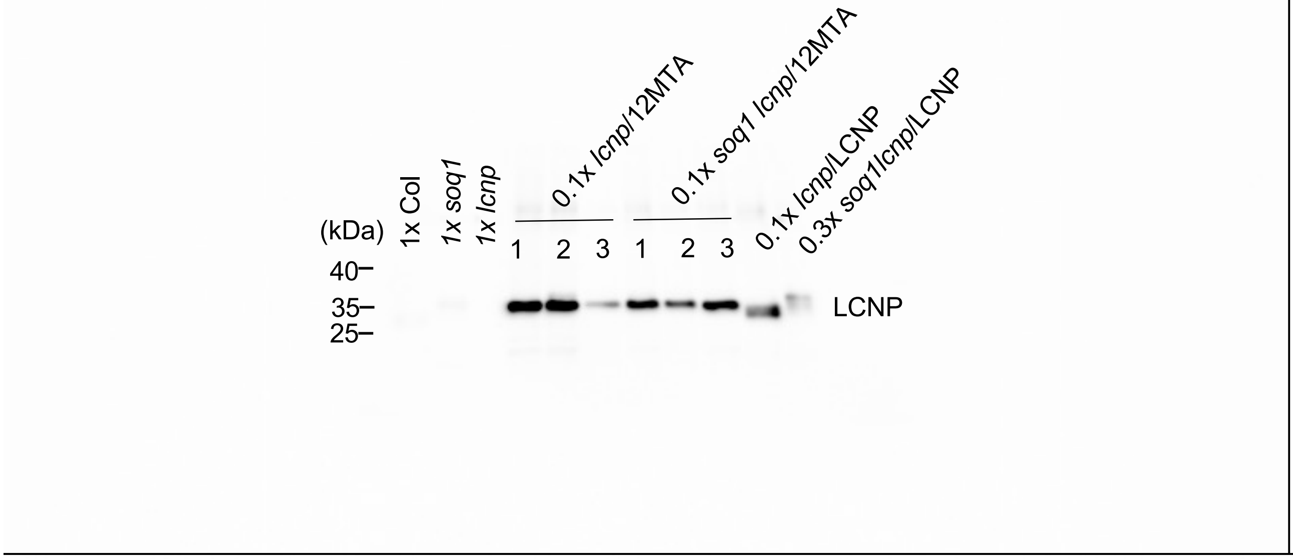


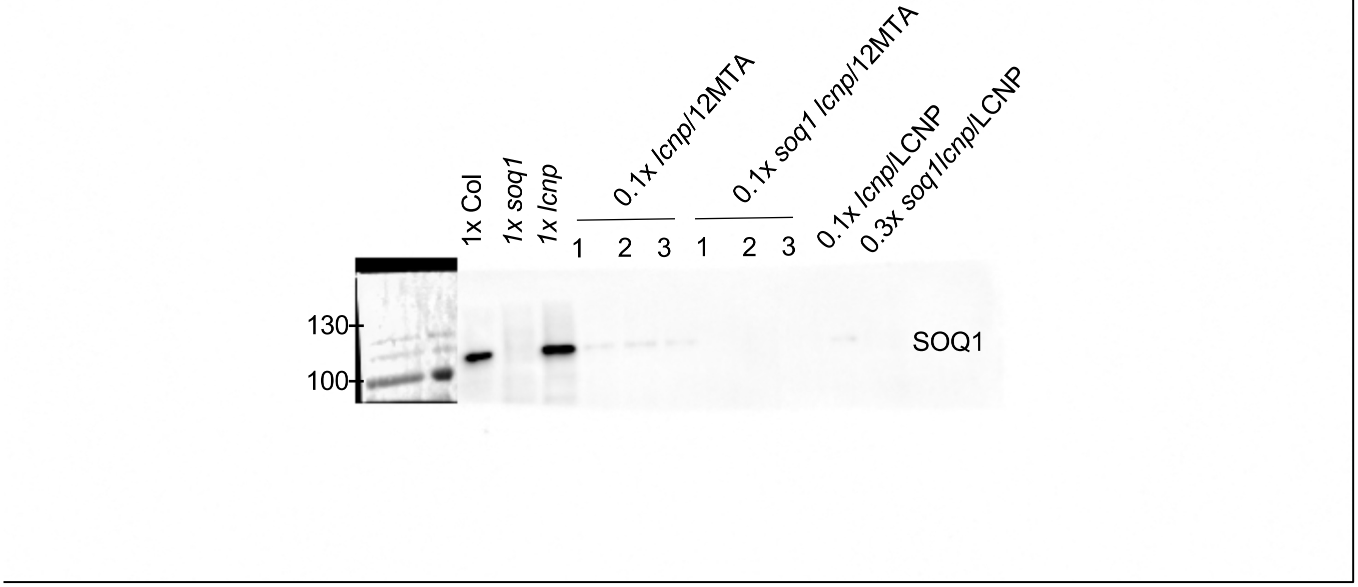


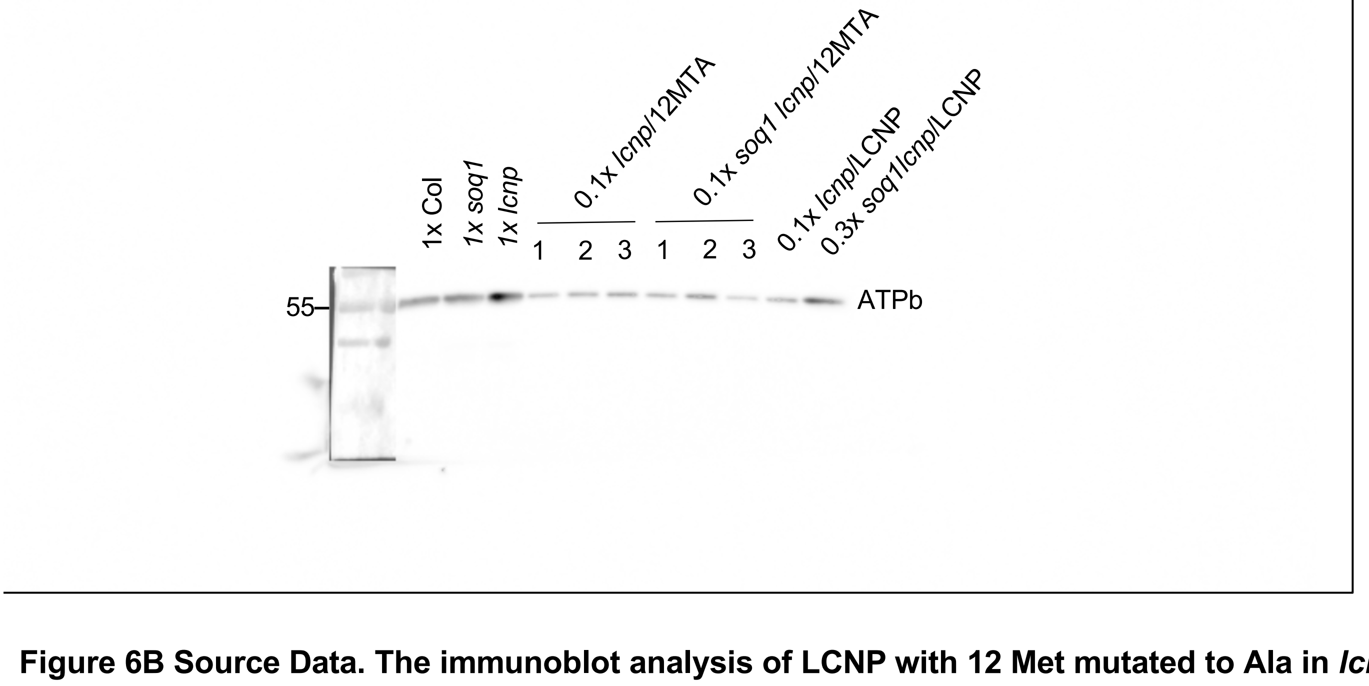


**Figure 6B Source Data. The immunoblot analysis of LCNP with 12 Met mutated to Ala in *lcnp* and *soq1 lcnp* backgroud.** Uncropped version, PVDF membrane was cut and probed with anti-LCNP, anti-$SOQ1_{CTD}$ and anti-ATPb antibodies.

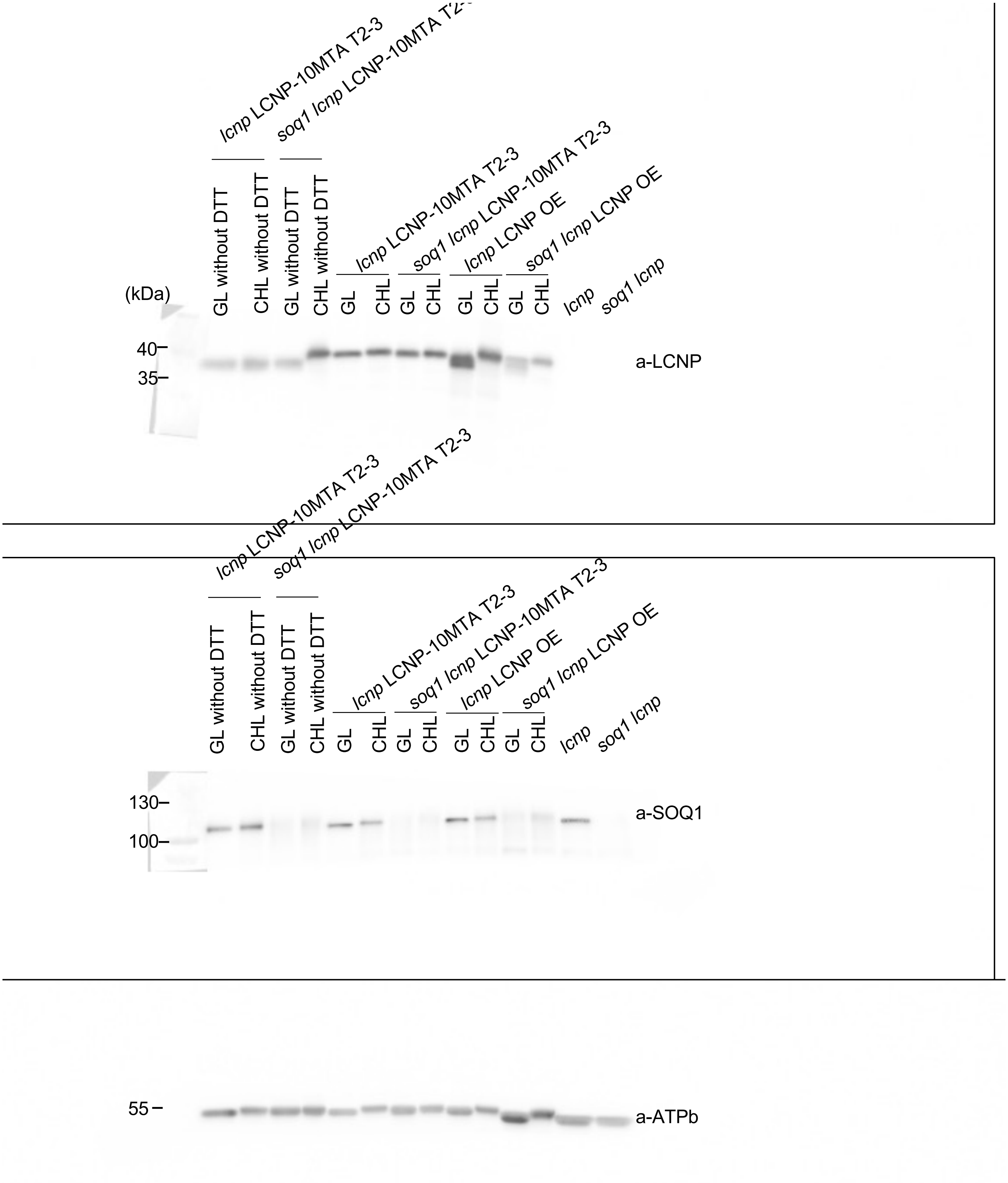


**Figure S6C Source Data. Slower electrophoretic mobility of LCNP is methionine-dependent.** Uncropped version, PVDF membrane was cut and probed with anti-LCNP, anti-$SOQ1_{CTD}$ and anti-ATPb antibodies. Support Figure 6B.

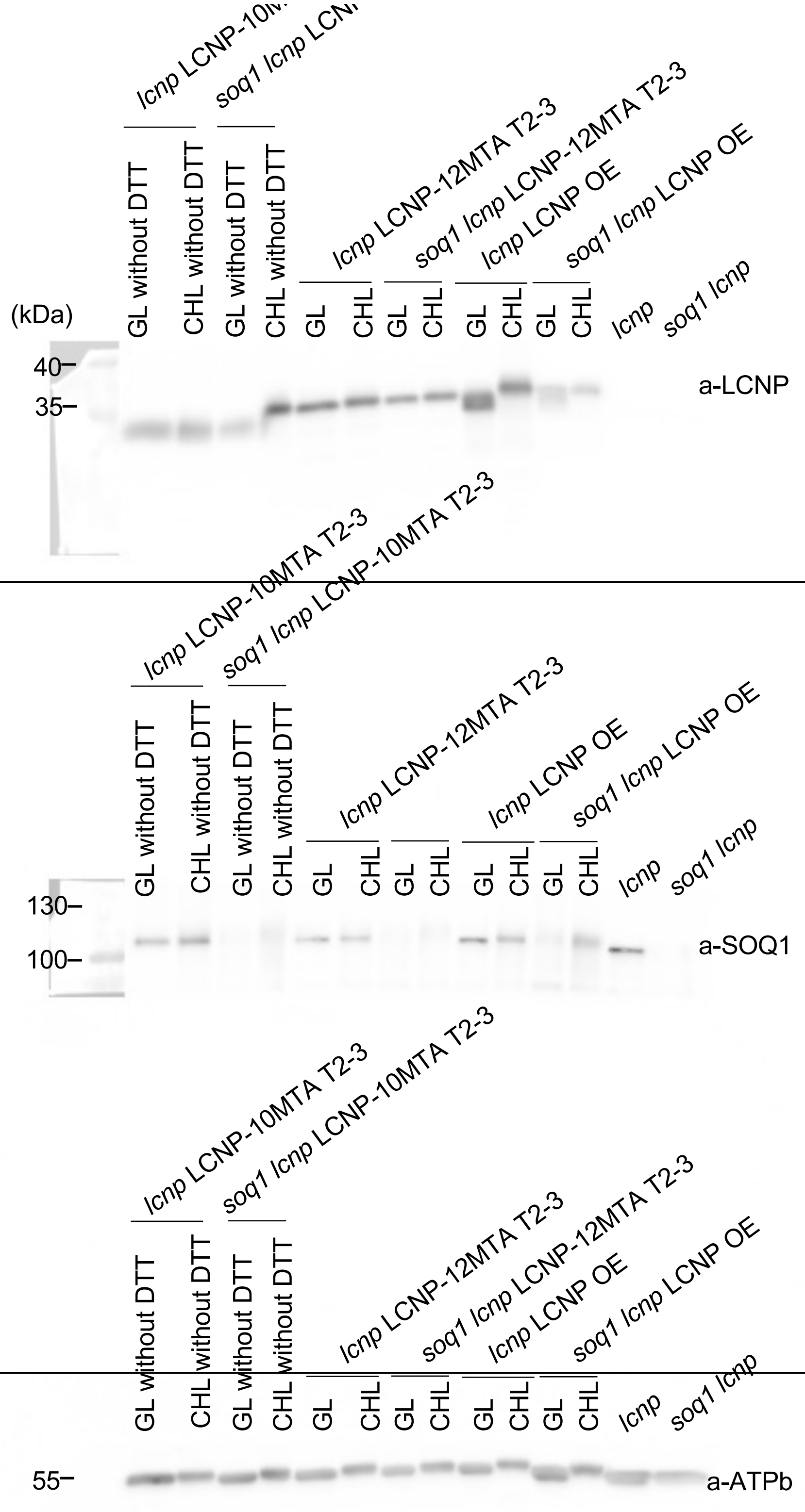


**Figure S6D Source Data. Slower electrophoretic mobility of LCNP is methionine-dependent.** Uncropped version, PVDF membrane was cut and probed with anti-LCNP, anti-SOQ1$_{CTD}$ and anti-ATPb antibodies. Support Figure 6C.
